# A single-cell transcriptome atlas of human early embryogenesis

**DOI:** 10.1101/2021.11.30.470583

**Authors:** Yichi Xu, Tengjiao Zhang, Qin Zhou, Mengzhu Hu, Yao Qi, Yifang Xue, Lihui Wang, Yuxiao Nie, Zhirong Bao, Weiyang Shi

**Affiliations:** Developmental Biology Program, Sloan Kettering Institute, New York, NY 10065, USA; Institute for Regenerative Medicine, Shanghai East Hospital, School of Life Sciences and Technology, Tongji University, Shanghai 200123, China; Developmental Biology Program, Sloan Kettering Institute, New York, NY 10065, USA; Traditional Chinese Medicine Hospital of Kunshan, Suzhou, Jiangsu 215300, China; Ministry of Education Key Laboratory of Marine Genetics and Breeding, College of Marine Life Sciences, Ocean University of China, Qingdao 266003, China; School of Pharmacy, Fudan University, Shanghai 201210, China; Ministry of Education Key Laboratory of Marine Genetics and Breeding, College of Marine Life Sciences, Ocean University of China, Qingdao 266003, China; Institute for Regenerative Medicine, Shanghai East Hospital, Tongji University, Shanghai 200123, China

## Abstract

The early window of human embryogenesis is largely a black box for developmental biologists. Here we probed the cellular diversity of 4- to 6-week human embryos when essentially all organs are just laid out. Based on over 180,000 single-cell transcriptomes, we generated a comprehensive atlas of 313 cell types in 18 developmental systems, which were annotated with a collection of ontology and markers from 157 publications. Together with spatial transcriptome on embryonic sections, we characterized the molecule and spatial architecture of previously unappreciated cell types. Combined with data from other vertebrates, the rich information shed light on spatial patterning of axes, systemic temporal regulation of developmental progression and potential human-specific regulation. Our study provides a compendium of early progenitor cells of human organs, which can serve as the root of lineage analysis in organogenesis.

## Introduction

Human embryogenesis finishes gastrulation by 2.5 week and at 4 week major embryonic organ and tissue types start to differentiate^1^. The transition from gastrulation to organogenesis is marked by sharp increase in cellular diversity generated from early progenitors. It is also at this stage that most developmental defects start to arise which could lead to miscarriage or birth defects^1^. Studies on these processes in vertebrates are mostly carried out in model systems such as the mouse and zebrafish^2–5^ but to which degree they are conserved in human embryo is unknown due to technical difficulties and ethical limitations. While single-cell data have been examined for human embryogenesis at later stages, either systematically^6,7^ or in an organ-specific manner^8–17^, the critical time window of great expansion of cellular diversity remains to be explored. Here, we examine this developmental window by studying human embryos at Carnegie stages (CS) 12-16 (4- to 6-week) and provide a comprehensive single-cell transcriptome atlas of early human embryo.

### Comprehensive map of human embryonic developmental systems and cell types

To systematically define the developmental landscape of human organogenesis, we obtained seven morphologically normal human embryos from CS12 to 16 (Fig. 1A, Extended Data Fig. 1), and used the 10x Genomics Chromium platform to obtain scRNA expression profiles. To increase sampling rate of cells, we dissected the embryos into major parts including head, trunk, viscera, and limb (Extended Data Fig. 1A). In total, we obtained 185,140 high-quality cells from 22 embryonic dissection parts. We captured an average of 7,732 transcripts and 2,338 genes per cell (Extended Data Fig. 1B). The sex of each embryo was determined by sex-specific gene expression (4 males and 3 females, Extended Data Fig. 1C). No abnormality was observed on copy number variation estimated by CopyKAT^18^, a method designed for 3’ or 5’ scRNA-seq at sparse coverage (Extended Data Fig. 1E, log2 ratios range from 0.92 to 1.06).

**Fig. 1.**
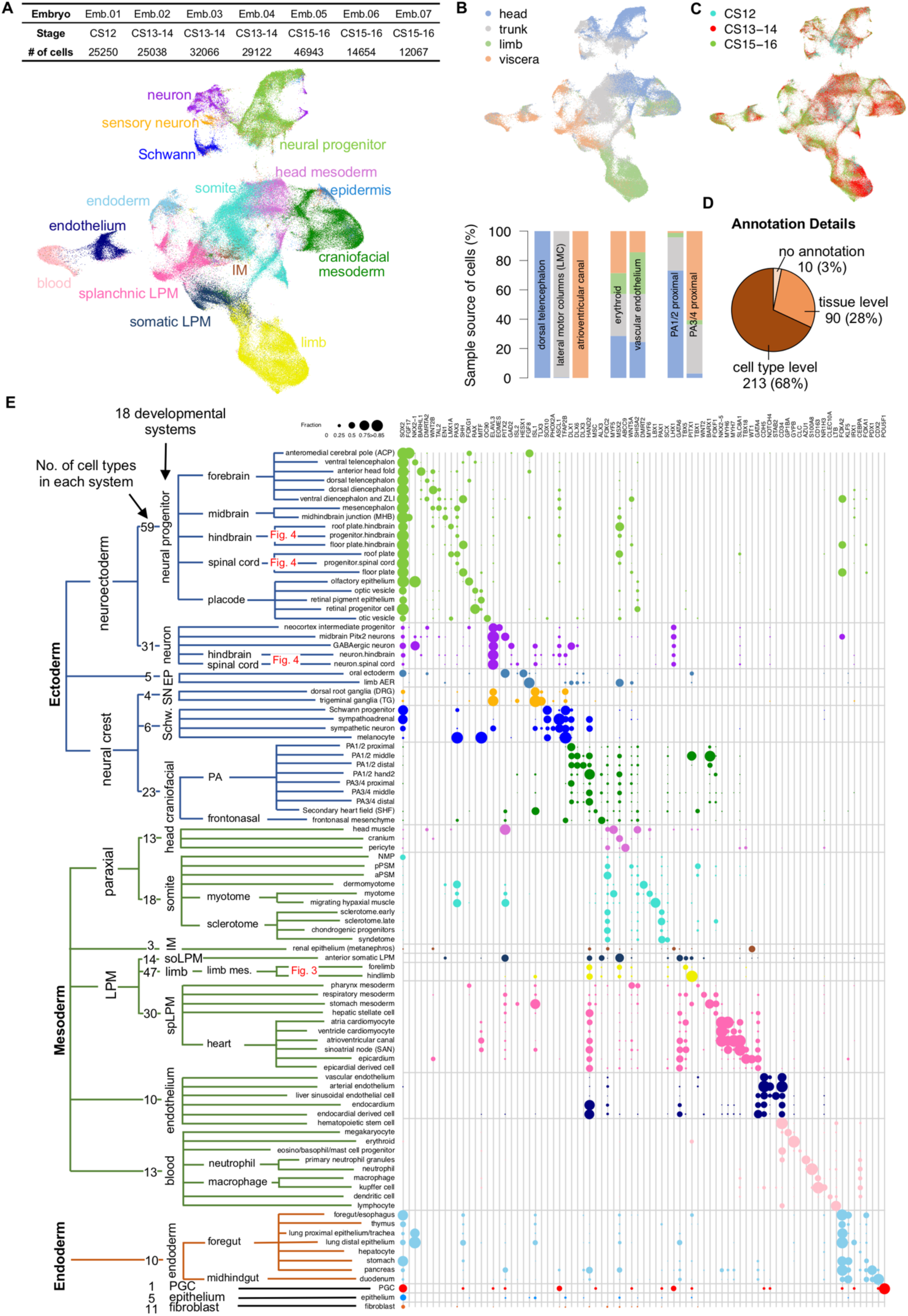
Single-cell transcriptome atlas of early human embryo. (**A**), Upper panel, Carnegie stage and number of cells of sampled embryos. Lower panel, uniform manifold approximation and projection (UMAP) visualization of high-quality cells colored by major developmental systems. IM, intermediate mesoderm; LPM, lateral plate mesoderm. (**B**), Upper panel, UMAP colored by dissection parts. Lower panel, the origin of dissection part for each cell in selected cell types. For all cell types, see Extended Data Fig. 6. (**C**), UMAP colored by embryonic stages. (**D**), Number and percentage of cell types annotated at cell type level, tissue level or without annotation. (**E**), Expression of known markers for cell types collapsed by unique terms in annotation (Supplementary Table 1C). Cell types in limb and neural tube were not showed, which are detailed in Fig. 3 and Fig. 4, respectively. Dendrogram of cell types was defined according to the lineage relationship based on annotations. The third level of dendrogram shows 18 developmental systems and number of cell types in each system. Dot size represents the fraction of expressing cells (UMI > 0) for a given gene and dot color shows developmental system. EP, epidermis; SN, sensory neuron; soLPM, somatic LPM; spLPM, splanchic LPM; PGC, primordial germ cell; PA, pharyngeal arch; ZLI, zona limitans intrathalamica; AER, apical ectodermal ridge; NMP, neuro-mesodermal progenitor; a/pPSM, anterior/posterior presomitic mesoderm.

To identify cell types, we conducted semi-supervised clustering, where knowledge of human and mammalian lineage differentiation was used to optimize *de novo* clustering to best recapitulate the known lineage hierarchy. We curated a total of 157 publications that define the developmental systems (major lineages, organs, and tissues) and the known cell types within each system at human CS12 to 16 or the corresponding mouse stages (E9.5 to E11.5), with a total of 234 diagnostic markers (Supplementary Table 1). To first resolve the developmental systems and then cell types within each, we applied iterative clustering (Extended Data Fig. 2-4, Supplementary Note 1) and identified a total of 313 cell types/clusters in 18 developmental systems (Fig. 1A). Notably, the mesoderm displays the most tissue-type and transcriptional diversity at this stage. Eight mesodermal systems are present, including two paraxial mesoderm (head mesoderm and somite), the intermediate mesoderm, three lateral plate mesoderm (LPM; limb, somatic LPM and splanchnic LPM), as well as blood and endothelium. Technically, we found that clustering based on transcription factors (TFs) better resolves the developmental systems than using all highly variable genes (HVGs) (Extended Data Fig. 2, Methods), possibly because of convergent expression of genes such as epithelial and extracellular matrix pathways across lineages.

Among the 313 cell types, we assigned 213 to known cell type identity and another 90 less specifically to a tissue identity (e.g., telencephalon) (Fig. 1D), generating 177 unique terms in annotation (Fig. 1E, Supplementary Table 1). The identified cell types include small anatomical structures such as 9 signaling centers (see below) and sensory placodes, as well as migratory cell types that would have been difficult to identify without the whole-embryo approach, such as the neural crest derived second heart field (SHF)^19^ marked by *MSC* and *ISL1*. Notably, many cell types were annotated only at the tissue level in head mesoderm and somatic LPM, reflecting our lack of understanding of these systems compared to the others. Based on our annotation, 47 pairs of cell types in this dataset are known to have close lineage relationship, such as from Schwann progenitor to melanocyte and from epicardium to epicardial derived cell (Supplementary Table 1C). Thus, we provide a deep annotation of cell types of human early organogenesis.

In addition to the literature and marker support, we further assessed the quality of our cell types/clusters through a series of quantitative tests (Methods). First, from technical standpoint, cross-validation at cell type level in each developmental system by scPred^20^ showed a mean of 0.89 on AUROC (area under the receiver operating characteristic) (Extended Data Fig. 4F), indicating clusters are well separated and cells can be robustly recalled to its own cluster. Also, cell types are not biased by batch, sequencing depth, or cell cycle states of cells (Extended Data Fig. 4G, 4H). Second, comparing with single-nucleus RNA-seq in whole mouse embryos at the corresponding stages^2^, 92% of human cell types can match to at least one mouse cell type (specificity score > 0.05) with 83% of matched mouse cell types agreeing on developmental systems (Extended Data Fig. 5, Supplementary Table 2), indicating that most of human cell types are supported by in an independently annotated dataset despite species and technical variations. Third, the spatial and temporal information preserved in sample collection can be used to test the quality of cell type identification. We found that more than 98% of all cells came from the expected dissection parts of the annotated cell type (Extended Data Fig. 6A, Supplementary Table 1). For example, dorsal telencephalon and lateral motor columns were only from head and trunk, respectively, whereas embryo-wide cell types such as endothelium and erythroid cells were from all dissection parts (Fig. 1B). Cell types from dissection boundaries show multiple origins, such as pharyngeal arches. We also examined stage distribution of cells in each cell type. Only 1 cell type may have bias on sampling or clustering because the middle stage CS13-14 is underrepresented (Extended Data Fig. 6B). These results suggest that computational artifacts are rare in the defined cell types.

To characterize cell types from the perspective of gene expression, we identified 3,698 differentially expressed genes (DEGs) across all cell types (Methods), including 206 (94%) of the canonical markers we collected. The number of DEGs for each cell type ranged from less than 40 in epidermis to more than 200 in neurons (Extended Data Fig. 6C). To identify cell type specific markers, we ranked DEGs by z-score for each cell type. 88% of the canonical markers belong to the top 20 most specific DEGs in each cell type (Supplementary Table 1). Many of the top DEGs by z-score have not been identified as cell specific markers in prior studies, e.g., *FOXL2* and *MMP23B* for SHF. More specifically, we interrogated ligands^21^ in the DEGs of 9 known signaling centers identified in our data, including the ZPA (zone of polarizing activity) and AER (apical ectodermal ridge) in limb patterning^22^, and 7 in the neural tube and brain^23^, which consists of less than 1.8% of the cells sampled. Our data captured the known signaling molecules in these 9 signaling centers, such as *SHH* in the floor plate^23^, *SHH* and *BMP4* in ZPA^22^ and *FGF8/9* in AER^22^, demonstrating the fidelity of our data in recovering small anatomical structures (Extended Data Fig. 6D). Moreover, we identified 33 additional signaling molecules from these centers^24^, ranging from 2 to 11 per signaling center. Compared to scRNA-seq data in mouse embryos^25–27^, almost all known signaling molecules are conserved despite potential batch effect and 57% of additional signaling molecules are expressed in the corresponding mouse cell types, which may implicate human mouse difference (Extended Data Fig. 6E).

To allow easy access of the expression data, we constructed a web-based database (Human early organogenesis atlas, HEOA, https://heoa.shinyapps.io/base/) for visualization, extraction of cell type specific expression profiles as well as comparison of gene expression between cell types.

### Orthogonal investigation by spatial transcriptome

To elucidate fine-grained spatial organization of cell types beyond dissection parts, we performed spatial transcriptome (ST) with 55-μm diameter spot on two sagittal sections (one center plane and one off-center plane) from another human embryo at CS13 (Fig. 2A, Extended Data Fig. 7A, 7B, Methods). Comparing with spatial transcriptome in mouse embryos at E10.5^28^, human sections show similar architecture with mouse sections in unsupervised co-clustering of spots (median r = 0.76, Extended Data Fig. 7C, 7D). To resolve the localization of cell types, we deconvoluted each spot in ST by RCTD^29^ with 239 cell types from scRNA-seq that are expected to be in the vicinity of the sections (Methods, Supplementary Note 2 and 3). The two sections are mainly mesodermal and ectodermal structures, respectively, which have all developmental systems in place (Fig. 2B) and in total detected 93% of cell types in deconvolution (Fig. 2F). On the cell type level, the deconvolution results confirmed the localization of recognizable structures on hematoxylin and eosin (H&E) staining, such as optic vesicle and renal epithelium (Fig. 2A, Extended Data Fig. 7E). Many structures that are difficult to be recognized only by H&E staining were also revealed, such as head muscle and pancreas (Extended Data Fig. 7F). Specifically, in heart, as expected, atria and ventricle cardiomyocyte are arranged from cranial to caudal, with endocardium inside and epicardium outside^13^ (Fig. 2C). Brain vesicles appear in an expected orientation on both rostral-caudal and dorsal-ventral axes (Fig. 2D). Migrating cell type SHF was detected in both origin and destination (Extended Data Fig. 8A). Other types of cases were also resolved, such as differentiation-related localization of sclerotome and systemic distribution of endothelium (Extended Data Fig. 8A). In summary, our scRNA-seq and ST data are coherent, thus providing a compendium of cell types with spatial information.

**Fig. 2.**
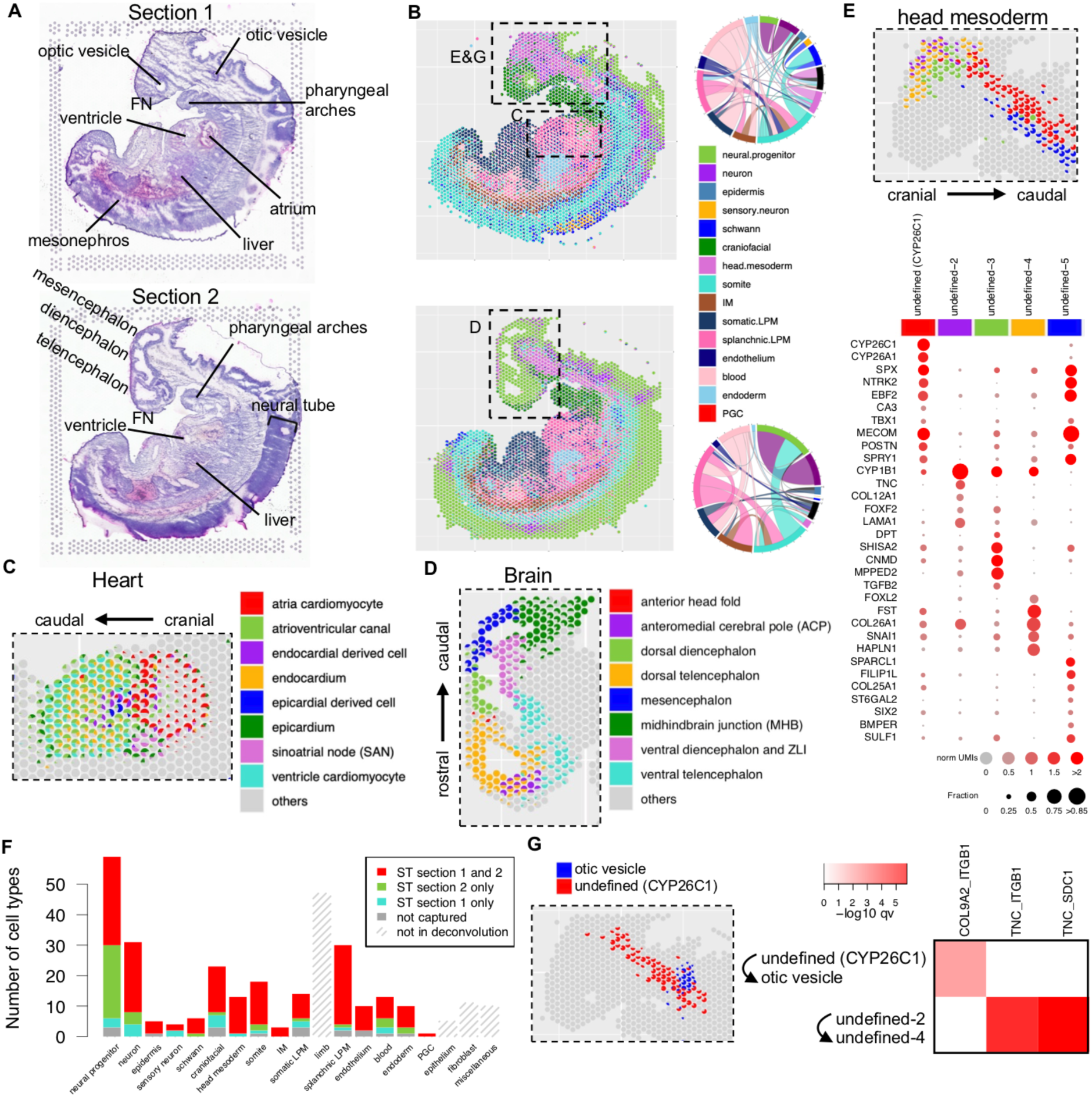
Spatial transcriptome of a CS13 embryo. (**A**), Two sagittal sections of a human embryo at CS13. Recognizable structures based on H&E staining are marked. FN, frontonasal mesenchyme. (**B**), The proportion of developmental systems in each spot by deconvolution (Methods) is showed as a pie chart. Black boxes denote the regions that are showed in panels C, D, E, and G. The chord diagram shows the number of spots with co-localization of developmental systems (proportion > 0.4 for both systems) on each section. (**C**), The proportion of heart cell types in each spot at the heart region of section 1 is showed as a pie chart. (**D**), The proportion of brain cell types in each spot at the brain region of section 2 is showed as a pie chart. (**E**), Upper panel, The proportion of 5 undefined cell types from head mesoderm in each spot on section 1 is showed as a pie chart. Lower panel, top DEGs of the 5 undefined cell types from scRNA-seq. (**F**), The detection of cell types on two sections by developmental system. At least one spot with proportion > 0.05 (equivalent to 1 cell) is required for the detection of a cell type. (**G**), Left panel, the proportion of undefined (*CYP26C1*) from head mesoderm and otic vesicle in each spot of section 1 is showed as a pie chart. Right panel, the significant signaling interactions involved in head mesoderm (Methods). The heatmap shows -log10 adjusted p value of hypergeometric test of a ligand-receptor pair (column) in a cell type pair (row).

We next explored whether previously uncharacterized developmental systems can be better understood with ST. Head mesoderm at this developmental stage contains 7 somitomeres^30^ but its cell types are not clear. We found 5 undefined cell types in head mesoderm have abundant detection in ST (Fig. 2E and Extended Data Fig. 8B). Importantly, they appear in the region of somitomeres in ST and have distinct distribution along anterior-posterior and dorsal-ventral axes. These 5 undefined cell types have robust DEGs between each other (Fig. 2E). The expression of DEGs largely agree with deconvolution, suggesting localizations of these cell types by deconvolution are rational (Extended Data Fig. 8C). They have clear separation in cross-validation (mean of AUROC 0.91). The most anterior cell types, undefined-2, has a AUROC score of 0.84 although its localization partially overlaps with undefined-4. The detection of the 5 cell types is correlated between two modalities, scRNA-seq and ST (Extended Data Fig. 8D, section with mainly mesodermal tissues r = 0.92, section with mainly ectodermal tissues r = -0.29 as control). These results indicate that the undefined cell types we found in head mesoderm are most likely real cell types that have not been described before. To further characterize these cell types, we asked whether there are signaling interactions involving them, giving mesodermal tissues usually send signals for the differentiation of surrounding tissues. We developed a statistical procedure to infer signaling interaction using scRNA-seq and ST (Extended Data Fig. 8E, Methods). In total, 134 significant signaling interactions were identified using this approach (adjusted p value of hypergeometric test < 0.01, Supplementary Table 1), including known events such as *SHH* from floor plate^31^, *FGF8/17* from anteromedial cerebral pole^32^, *BMP2/7* from myocardium^33^ (Extended Data Fig. 8F). Although spinal neurons and sclerotome both receive sonic hedgehog signaling from floor plate^31^, we found its antagonist HHIP is specific to sclerotome. Undefined cell types in head mesoderm were found to have interactions with otic vesicle and interactions between them (Fig. 2G, Extended Data Fig. 8G). These signaling events most likely contribute to morphogenesis, including collagen-integrin interaction^34^ and TNC, an extracellular matrix protein associates with morphogenetic events of anterior head formation^35^. Together, our scRNA and ST uncovered previously unappreciated cell types in head mesoderm and potential interactions with neighboring tissues.

### Patterning of axis formation

Bilaterian body plan is laid out along three main body axes, the proximal-distal (PD), anterior-posterior (AP) and dorsal-ventral (DV) axes. Axis patterning is a prominent and ubiquitous process during embryogenesis as each organ elaborates the three axes via diverse mechanisms. Studies have provided classic paradigms of spatial patterning albeit in model organisms. Here we examine axis formation in limb and neural tube development in human.

Morphologically, the limb buds start to appear at CS12. During this time, PD, AP and DV axes are established through interactions of signaling gradients along different directions^36^. However, cell types in early limb bud are underdefined despite the extensive *in situ* data due to difficulties in inferring the boundaries between *in situ* expression domains. To this end, we annotated cell types in our scRNA-seq data with *in situ* results in mouse^37–46^ to locate cell types in the forelimb mesenchyme at each stage. Altogether, we defined 6, 10, 15 spatial domains (‘domain’, a cell type with identified spatial location) along the PD and AP axes for CS12, CS13-14 and CS15-16 forelimb, respectively (Fig. 3A, Extended Data Fig. 9-11, Methods). The spatial organization of cell types was largely recapitulated by UMAP, where domains are displayed following the PD and AP axes and D-to-P flows in RNA velocity^47^ analysis matches previous lineage tracing results^48^ (Fig. 3B). The clustering of domains is not biased by embryo (Fig. 3C, Extended Data Fig. 9B, Mann-Whitney U test p < 10^-16^). As *post hoc* validation, the expression of HOXA genes on our domain map recapitulated known HOX pattern^4^^9^ with minor exception (Fig. 3D). Importantly, 17 out of 20 patterning genes (e.g., *HAND1*, *ZIC3*, *IRX3*) independently selected by Zhang and colleagues in spatial transcriptome of human limb bud at similar stage^50^ display consistent expression pattern on our domain map at CS15-16 (Supplementary Note 1), indicating the high fidelity of our spatial reconstruction.

**Fig. 3.**
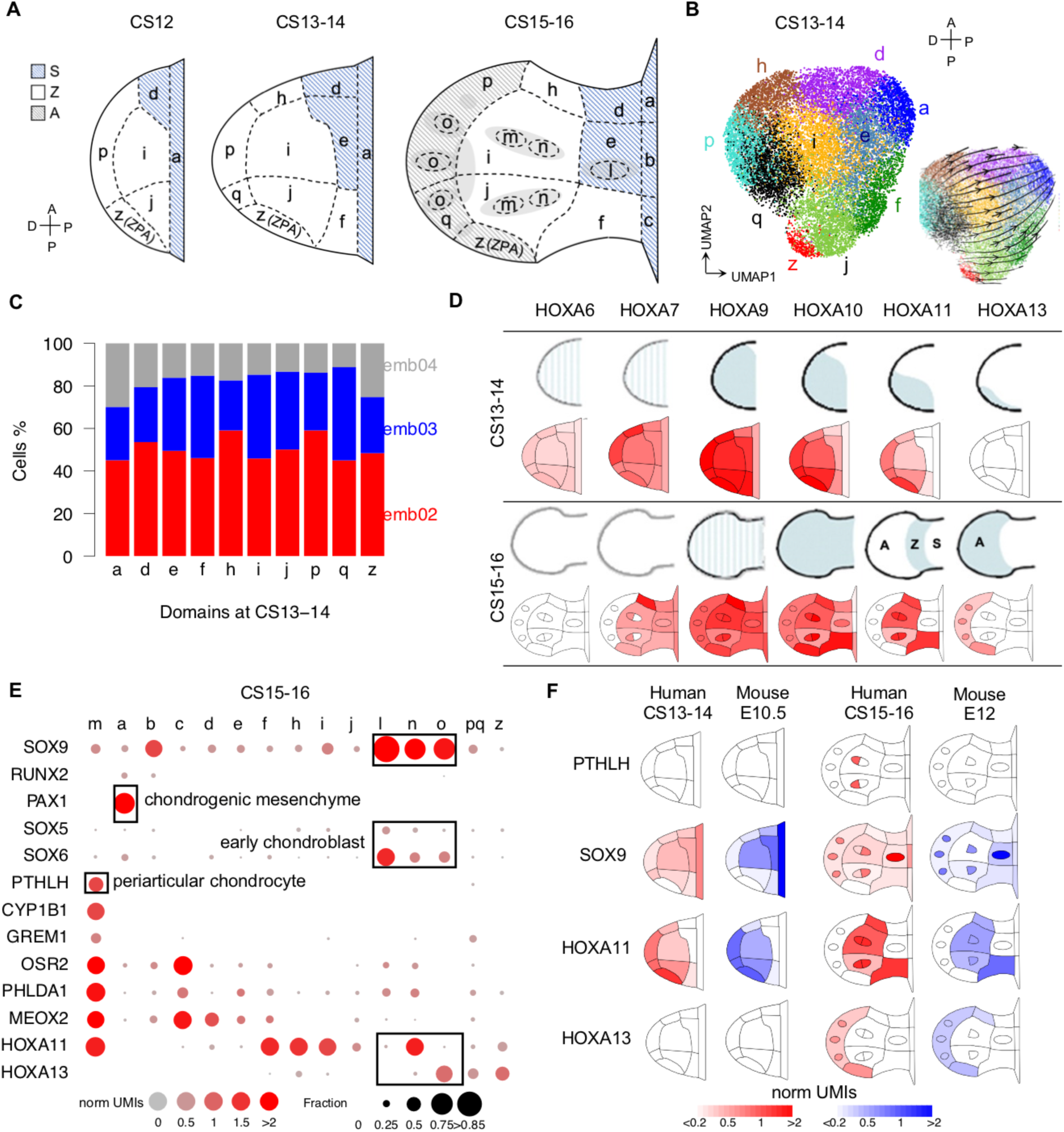
Spatial domains in limb bud. (**A**), Diagram depicting domains in forelimb mesenchyme defined in this study at each developmental stage. Domains from stylopod (S), zeugopod (Z) and autopod (A) are indicated by different colors. D-P, distal-proximal; A-P, anterior-posterior. (**B**), UMAP visualization of cell types and RNA velocity analysis in forelimb at CS13-14. Cells are colored by domains, as labeled in panel A. (**C**), The percentage of cells from each embryo in each domain at CS13-14. (**D**), Comparison of expression pattern of HOXA genes between *in situ* and scRNA-seq. The expression in scRNA-seq was summarized as mean of normalized unique molecular identifiers (UMIs) in each domain. (**E**), The expression of marker genes in chondrocyte differentiation (boxes) and selected DEGs of domain *m* at CS15-16. (**F**), The comparison of expression pattern of *PTHLH*, *SOX9*, and HOXA genes between human (red) and mouse (blue) at two stages.

At CS15-16, we found that five domains (*a*, *l*, *m*, *n*, and *o*) are at different degrees of chondrogenesis^51,52^ (Fig. 3A, E), including domain *m* in the zeugopod territory that is distinguished by *PTHLH* and other DEGs (AUROC = 0.94). Of note, *PTHLH* has not been characterized by *in situ* in mouse at corresponding stage. Domain *m* is robustly identified in all three embryos (82∼135 cells per embryo) and its DEGs are robustly expressed in each embryo (Extended Data Fig. 9C). To compare to mouse at the corresponding stages, we annotated domains by reclustering a published scRNA-seq data in mouse forelimb at E10.5 and E12^26^ for human CS13-14 and CS15-16, respectively. All domains except domain *m* were identified in mouse by the same set of marker genes used in human data (Extended Data Fig. 9D-E, Supplementary Table 2). The master regulator of chondrocyte *SOX9* shows consistent expression pattern between human and mouse (Fig. 3F). However, specific markers of domain *m* (*PTHLH* and *CYP1B1*) were almost not co-expressed in any cell of E12 mouse (Extended Data Fig. 9F, 9G). To exclude potential artifacts in clustering, we examined the co-embedded space of human and mouse cells (Extended Data Fig. 9H). Indeed, cells in domain *m* have significantly fewer mouse cells in nearest neighbors (Mann-Whitney U test p = 10^-9^). Domains *a* and *b* in human also have fewer mouse correspondences but to a lesser extent, likely because proximal structures were partially lost during dissection of limb in mouse dataset. These results suggest domain *m* in human is not present in mouse at E12. Domain *m* may appear later in mouse because *Pthlh* was reported to have localized expression in limb at E12.5 but not earlier, and is attributed to a cell type called periarticular chondrocyte^53^. The stage alignment between species was justified by the expression of HOX genes (Fig. 3F), where the switch of HOX pattern is under strict temporal regulation in limb bud^54^. Therefore, based on the fine map of limb bud, we found heterochrony on chondrogenesis between human and mouse.

The neural tube is patterned along the AP axis by HOX genes^55^ and the DV axis by opposing signaling gradients to form regionalized cell types^56^. We examined these patterns in human in contrast to those in mouse.

Based on the expression of HOX genes, we reconstructed the AP axis for the neural progenitors of the hindbrain and the spinal cord (Fig. 4A). HOX expression along the pseudo AP axis agreed with dissection parts of cells. In addition, cell types were dispersed along the pseudo AP axis, demonstrating that our cell type identification was not affected by the AP gradient of gene expression. We then identified 21 genes whose expression level display an AP gradient besides the HOX genes (Fig. 4B). Among these, the *CYP26C1* gene is known to be involved in establishing the retinoic acid gradient that is responsible for activating some of the HOX genes along the AP axis^57^. Notably, these genes include five long non-coding RNAs (lncRNA) situated in the HOX gene clusters, namely *HOTAIRM1*, *RP11-357H14.17*, *RP11-834C11.4*, *RP11-834C11.6* and *FLJ12825*. Other than *HOTAIRM1*, these lncRNAs are unique to the human genome^58^, indicating human-specific regulatory networks in neural tube AP patterning. The spatial pattern of the 5 lncRNAs were validated by spatial transcriptome (Fig. 4C, Pearson’s correlation 0.82∼0.91, Methods). Furthermore, we found that *HOTAIRM1* expression is not similar with the adjacent HOX genes (*HOXA2*) as by the general rule of lncRNA expression in the HOX gene cluster^59^ and the cases of the other four lncRNAs with AP gradient (Extended Data Fig. 12A). Instead, it appears to be correlated to the more distant *HOXA4*. The discordant expression between *HOTAIRM1* and adjacent HOX genes was consistent with previous finding in myeloid lineage^60^. Thus, our analysis expands the known AP gradient gene expression in vertebrate neural tube patterning and sheds light on human-specific regulation.

**Fig. 4.**
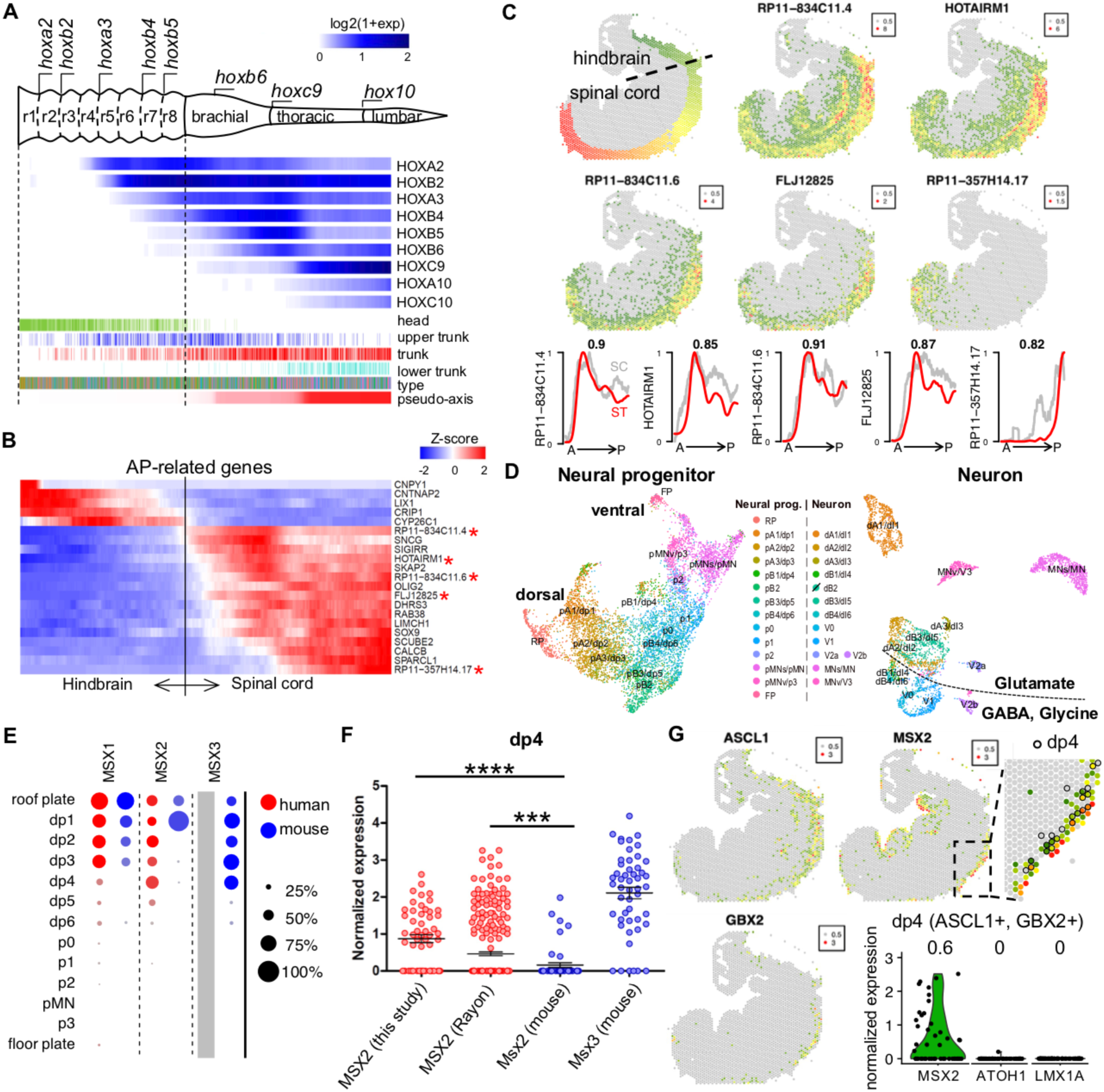
Spatial patterning of neural tube. (**A**), Schematic diagram modified from Philippidou *et al*.^55^ (upper panel) illustrating HOX gene expression along the vertebrate anterior-posterior axis. Heatmap (lower panel) displaying the expression of HOX genes along the pseudo-axis constructed with neural progenitor cells from hindbrain and spinal cord. Origin of dissection parts, cell types (see panel C for color code), and pseudo-axis value of each cell were shown below. (**B**), Expression patterns of AP-related genes based on pseudo-axis. Asterisks denote lncRNAs. (**C**), Top-left panel, AP values on section 2 of spatial transcriptome is showed as heatmap (Methods). The division between hindbrain and spinal cord was determined by morphology on section 2. Top panels, normalized UMIs of 5 lncRNAs on section 2 of ST. Bottom panels, expression of the 5 lncRNAs along pseudo-axis of scRNA-seq (grey) and AP axis of ST (red). Pearson’s correlations between two curves are marked on each panel. (**D**), UMAP visualization of neural progenitors (left) and neurons (right) from hindbrain and spinal cord. Cells are colored by cell types. ‘/’ on dB2 legend denotes dB2 is not detected in this dataset. (**E**), Comparison of expression of MSX paralogs in neural progenitor between human and mouse. Grey box denotes no *MSX3* in the human genome. **(F**), The expression of *MSX2* in dp4 in human (our dataset and Rayon *et al.*^63^) and mouse, as well as the expression of *Msx3* in dp4 in mouse. Each dot denotes a cell with confident dp4 identity from Extended Data Fig. 12D. ****, p value < 10^-7^; ***, p value < 0.001 (Mann-Whitney U test). **(G**), The expression of *ASCL1*, *GBX2*, *MSX2* on section 2 of ST and violin plot of *MSX2*, *ATOH1*, *LMX1A* expression in the dp4 spots (*ASCL1* > 0.5 and *GBX2* > 0.5 in neural tube, i.e., spots with AP values in panel C). In the zoom in area of *MSX2* heatmap, black circles denote dp4 spots. Each dot in the violin plot denotes a dp4 spot. The value above each bar indicates mean of normalized UMIs for each gene.

Vertebrate neuronal diversity is most evident along the DV axis^61^. We identified 13 neural progenitor and 13 neuronal cell types in the neural tube which can be recognized by the known marker expression in mouse^62^ (Fig. 4D). Given the conservation of neuronal cell types, we next asked to what extent homologous cell types have conserved gene expression between mouse and human. For this, we compared our data with scRNA-seq in mouse neural tube^62^. Neurons display virtually the same expression patterns of canonical markers for each type (Extended Data Fig. 12B). In contrast, progenitors show more difference (Extended Data Fig. 12B). Most of the differences are quantitative, *i.e.*, markers display the same DV spatial range but at different levels of expression, such as *MSX1* and *OLIG3*. A recent scRNA-seq study of human neural tube^63^ identified two TFs that are specific to human, namely *PAX7* and *NKX6-*2, which are confirmed in our dataset (Extended Data Fig. 12C). More importantly, we found that *MSX2* expands its expression ventrally in human compared to mouse (Fig. 4E, roof plate to dp4 in human and roof plate to dp1 in mouse). The expression of *MSX2* in human is similar to its paralog *MSX3* in mouse (roof plate to dp4), a gene that is lost in the human genome, suggesting *MSX2* compensates the function of *MSX3* in human dp2-4 (Fig. 4E). The expansion of *MSX2* expression is supported by the recent human dataset^63^ (Fig. 4F) and this is not caused by erroneous clustering in human or mouse datasets (Extended Data Fig. 12D). Lastly, we sought to examine *MSX2* expression in dp4 on section 2 of ST in a way that is independent with scRNA-seq. We located dp4 in the spots in neural tube that co-express dp4 markers (*ASCL1* > 0.5 and *GBX2* > 0.5). Indeed, *MSX2* is expressed in the dp4 spots in ST (Fig. 4G, mean = 0.6). *MSX2* expression in these spots are not from cell types known to express *MSX2*, namely dp1 or roof plate, as their specific markers *ATOH1* and *LMX1A* are not co-expressed with *MSX2* in these spots, indicating the spatial resolution of our ST is sufficient to validate the expansion of *MSX2* expression in human. Thus, master TFs can have considerably different expression between species even in a broadly conserved system like neural tube.

### Systemic temporal regulation of vertebrate embryogenesis

Vertebrate embryogenesis progresses through stages with characteristic developmental events, but it is not clear whether there exist conserved regulatory mechanisms that may control stage transitions. Here we examine systemic gene expression changes in our data in comparison to other vertebrate models, and implicated the potential role of LIN28A, a conserved RNA binding protein known to regulate developmental stage transition in *C. elegans*^64, 65^, in the regulation of vertebrate embryonic stage transition. While examining temporal gene changes in the early human embryo, we observed a sharp decrease of *LIN28A* expression (>2 fold) from CS12-14 to CS15-16 embryo in 96% cell types (Fig. 5A). Similarly, at the corresponding stages in mouse embryos (E9.5-10.5 vs. E11.5), *Lin28a* showed >2 fold decrease in 90% of reported cell types^2^ (Fig. 5B, 5C). Combining published bulk and single-cell RNA-seq in other vertebrate embryos in a broader time window^2,4,5,66–69^, we observed a conserved pattern of *LIN28A* expression: *LIN28A* is dramatically up-regulated at the beginning of gastrulation, peaking at early organogenesis, and diminishing at hatching in zebrafish/frog or E11.5/CS15-16 in mouse/human (Fig. 5D). Previous studies of LIN28A protein in mouse^70^ and in frog^71^ showed protein dynamics of LIN28A is consistent with mRNA dynamics. Thus, LIN28A displays a conserved and systemic dynamics in vertebrate embryogenesis.

**Fig. 5.**
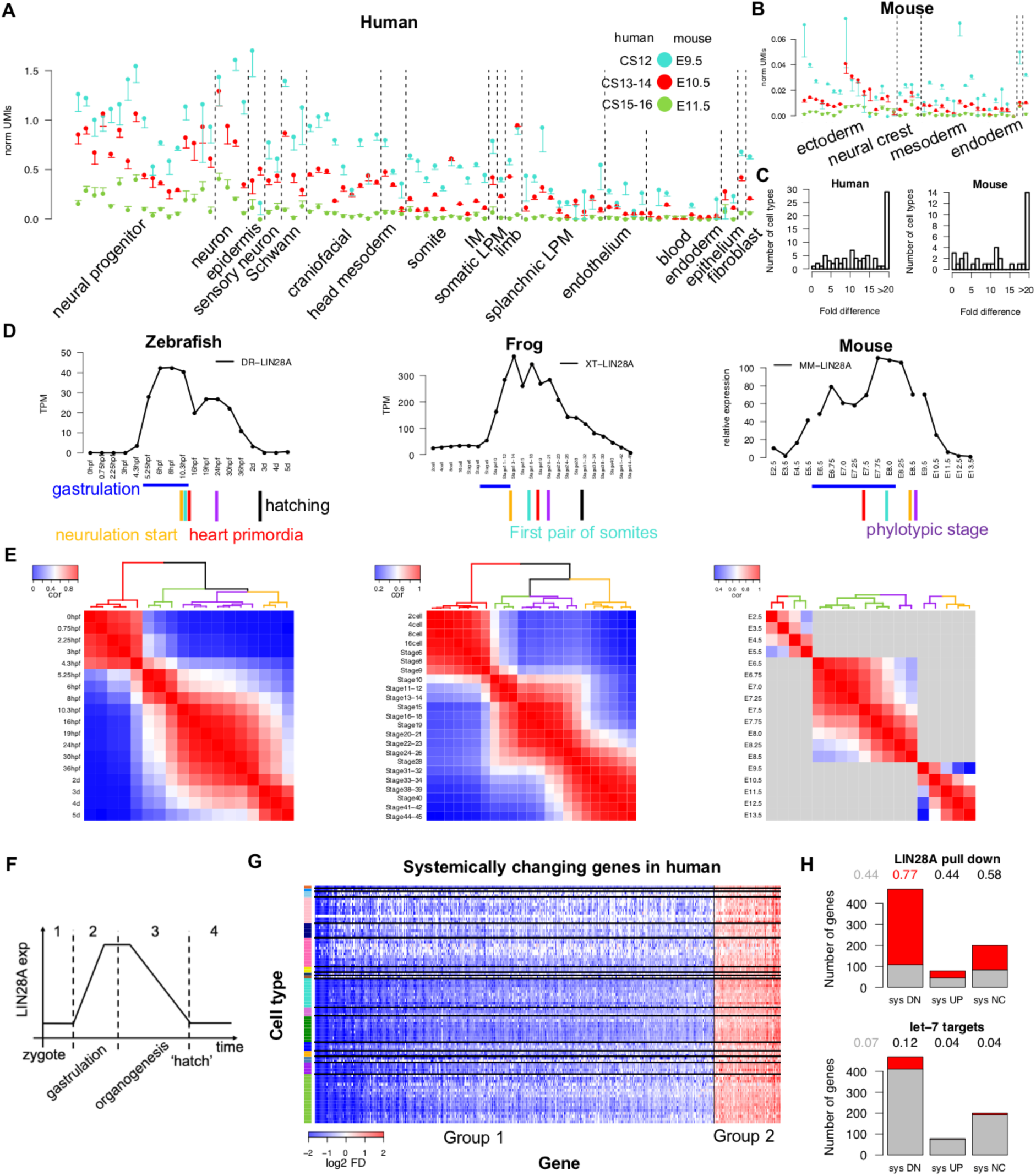
Systemic temporal regulation in vertebrate embryogenesis. (**A**), *LIN28A* expression (normalized UMIs +/-SE) in human embryos at CS12, CS13-14, and CS15-16 by cell type. (**B**), *Lin28a* expression (normalized UMIs +/-SE) in mouse embryos at E9.5, E10.5, and E11.5 by cell type. (**C**), The distribution of fold difference of *LIN28A* in each cell type between early and late stages in human (left) and mouse (right). (**D**), *LIN28A* expression in zebrafish, frog, and mouse embryos by time point. For mouse, lines between time points from different datasets were not drawn. Vertical lines below show the timing of major developmental events. (**E**), Heatmaps of Pearson’s correlation between time points in each species. Colors of dendrogram show clusters of time points. For mouse, pairwise correlation and clustering were only performed within each of the three mouse datasets (grey, inter-dataset entries). (**F**), A schematic diagram showing the dynamics of *LIN28A* across four developmental stages. (**G**), Heatmap showing log2 fold differences of systemically changing genes in each cell type in human from stage 3 to stage 4. Down-regulated and up-regulated genes were separated by vertical lines. For the color bar of cell types, see Fig. 1 for conventions. (**H**), Enrichment analysis of LIN28A pull-down targets and *let-7* targets in systemically down-regulated (‘sys DN’, group 1 in panel G), systemically up-regulated (‘sys UP’, group 2 in panel G), and unchanged genes (‘sys NC’) in human from stage 3 to stage 4. Red portion of bars represent targets in each group. Numbers above indicate the percentage of targets in whole genome (grey) and each group (red, hypergeometric test p value < 0.001 and odds ratio > 2).

Furthermore, we found that gene expression through vertebrate embryogenesis displays distinct temporal boundaries that correspond to landmarks in development, which we name punctuated developmental stages. Four such stages were revealed in each species by hierarchical clustering based on pairwise correlations of TF expression between timepoints (Fig. 5E, Extended Data Fig. 13A, Supplementary Table 3). In zebrafish and frog, stage 2 corresponds to gastrulation, stage 3 to organogenesis, and hatching at the boundary between stage 3 and stage 4^72, 73^. Similarly, 3 stage boundaries were observed in mouse (Fig. 5E). In addition, these 4 stages between any two species show strong pairwise correlations of expression of homologous TFs (Extended Data Fig. 13B), which is consistent with the temporal alignment based on sparsely sampled time series^74^. Intriguingly, the beginning of the 4th stage in mouse, E11.5, corresponds to birth in opossum based on tissue-level transcriptomes^75^. These results suggest that vertebrate embryogenesis consists of conserved and punctuated stages and raise an intriguing possibility that over the course of evolution of mammals some of the post embryonic development was shifted in uterus.

Given that the up- and down-regulation of *LIN28A* coincide with the transition from stage 1 to 2 and 3 to 4, respectively (Fig. 5F), we next exploited the potential of LIN28A on stage transition. Several lines of evidence suggest that LIN28A may regulate development during stage transition. First, three of the four major processes known to be regulated by LIN28A^76^ stood out among systemically changing genes (genes that show consistent changes across cell types) from stage 1 to stage 2 and from stage 3 to stage 4 (Fig. 5G, Extended Data Fig. 13C, Supplementary Table 4, Methods), namely cell cycle, mRNA splicing, and translation (Supplementary Table 5). The three pathways have 17, 36, and 10 genes, respectively, that are positively correlated to *LIN28A* expression in both zebrafish and frog (Extended Data Fig. 13D, Supplementary Table 5). Second, LIN28A may regulate systemically changing genes through direct binding of mRNAs, instead of the well-studied *let-7*-dependent pathway. The systemically down-regulated genes from stage 3 to stage 4 in human are enriched for LIN28A binding^77^ (Fig. 5H, hypergeometric test p = 10^-50^), while up-regulated genes and ubiquitously expressed genes with no significant changes in the time window are not. We found that none of these categories is enriched for the known *let-7* binding motif in human^78^ (Fig. 5H). Third, growth phenotypes of *Lin28a*(-) mouse embryos suggest that *Lin28a* is necessary and sufficient to promote growth no later than E9.5^79^, which corresponds to our stages 3 (organogenesis). Perturbation of *Lin28a* in mouse tailbud^80^ and lung^81^ led to heterochronic phenotypes, where loss of function caused precocious development and prolonged expression caused retarded development. Taken together, our results suggest punctuated stage transitions in vertebrate embryogenesis and regulation by a conserved mechanism via LIN28A.

### Integration with later stage human data

Due to practical limitations of specimen availability, our dataset provides the earliest timepoint in terms of human organogenesis when major organs are being laid out. It provides the root for inferring the developmental trajectories of different organs. In this regard, we conducted systematic data integration with scRNA-seq from later stages of human fetus (10-19 week old)^7^, which profiled fetal organs covering diverse systems but with some absent, such as limb and head mesoderm. Joint embedding after batch correction shows a global alignment of datasets between early and late stages in which developmental systems are aligned with distinct temporal trajectories (Fig. 6A, Extended Data Fig. 14A-B, Methods).

**Fig. 6.**
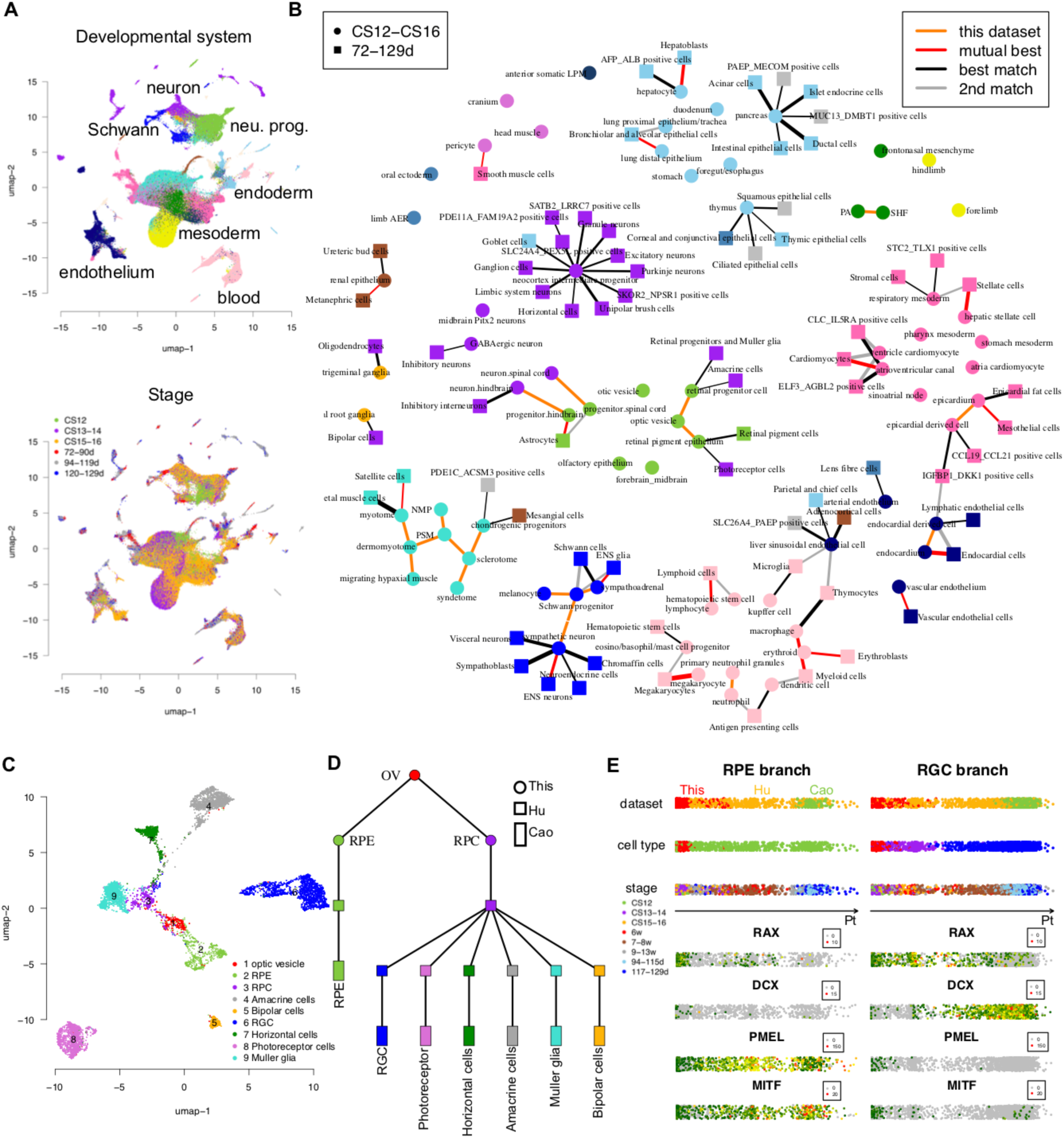
Systemic data integration. (**A**), UMAP of visualization of our dataset and Cao’s dataset^7^ colored by developmental system (upper panel, see Fig. 1 for conventions) and stage (lower panel). (**B**), The mutual best match (red edge), best match (black edge) and 2nd-best match (grey edge) in our dataset for each cell type in Cao’s dataset by Slingshot (also see Supplementary Table 6). Cell types are colored by developmental system (see Fig. 1 for conventions). The thickness of lines is inversely proportional to z-score across distances between a cell type in Cao’s dataset and all cell types in our dataset. The lineage relationship between cell types within our dataset (orange edge) was determined by annotation. (**C**), UMAP visualization of cell types of the developing eye in three studies colored by cell type. (**D**), Inferred cell trajectory in eye development by Slingshot and temporal information (Methods). Nodes are colored by cell type as in panel C. The shapes of nodes denote data source. (**E**), Pseudo-time ordering of cells in RPE branch and RGC branch by Slingshot. The color codes from top to bottom are dataset, cell type (see panel C), and stage. The heatmaps below show normalized UMIs of 4 marker genes along pseudo-time.

An effective alignment between datasets should reflect the lineage relationship between cell types. To evaluate this, we assigned one or two best matched cell types in our dataset for each of 73 cell types at the later stages based on the distances calculated by Slingshot^82^ (Fig. 6B, Supplementary Table 6, Methods). These mapping generally respect the unfolding of cell types in embryos from three aspects. First, almost all cell types in our dataset from the structures that were not sampled at the later stages were not matched to any cell type at the later stage, e.g., limb, craniofacial mesoderm, and somatic LPM. Second, 88% cell types of later stage are linked to the correct developmental system at the early stage (Supplementary Table 6). Some mismatches on system could be due to common epithelial features (corneal and conjunctival epithelial cells to thymus) and common neuronal features (bipolar cells to dorsal root ganglia). Third, on the cell type level, although most cell types do not have known lineage to be referenced, there are many unambiguous cases standing out. For example, endoderm-originated hepatoblasts and mesoderm-originated stellate cells from liver samples at the later stages were matched to hepatocyte and stellate cells in our dataset, respectively. Blood and endothelium at the later stages were identified from multiple organs, contributing to 6 mutual best matches from the same lineage of early cell types, which are erythrocytes to erythroblasts, megakaryocyte to megakaryocyte, lymphocyte to lymphoid cells, macrophage to myeloid cells, endocardium to endocardial cells and vascular endothelium to vascular endothelium. However, a few exceptions exist. For example, in eye development, only 2 out of 6 downstream cell types of retinal progenitor cells at the later stages were matched to retinal progenitor cells in our dataset. This analysis shows that one could establish eligible but imperfect mapping of cell types between early organogenesis and fetal stage.

Data from intermediate developmental stages would likely improve trajectories. In this regard, we further examined the trajectory of eye development, where organ-specific data is available at the intermediate stages. We collected scRNA-seq data on human eye field spanning from 5- to 13-week^9^ to fill the stage gap and integrate cell types in eye from three datasets (Fig. 6C, Extended Data Fig. 14C-D). Vertebrate eye forms from the optic vesicle that subsequently develops into a two-layer structure, i.e., retinal pigmented epithelium (RPE) and retinal progenitor cells (RPC). The multipotent RPC in turn gives rise to various types of retinal neurons, such as retinal ganglion cells (RGC)^83^ (Extended Data Fig. 14C). Using Chu-Liu/Edmonds’ algorithm^84^ to incorporate temporal information on transcriptome similarity, we reconstructed the trajectories of cell types in eye, which recapitulated the known lineage (Fig. 6D, Methods). Pseudo-time analysis on the RPE and RGC branches confirmed the known expression patterns of key regulators (Fig. 6E). *RAX* is initially expressed in the entire vesicle and then restricted to RPC^85^. The expression of *PMEL* and *MITF* in RPE is important for the differentiation of pigment cells^86, 87^. As shown in Fig. 6E, data from consecutive stages convey contiguous trajectories. Furthermore, we identified key genes and TFs that are specifically upregulated in each cell type, which predicts the key regulators that underlie the specification of each lineage from the optic vesicle as well as the temporal evolution within each lineage (Supplementary Table 6, Methods). More broadly, with improved temporal resolution in data collection, one could systematically infer organ-specific trajectories rooted by our dataset.

## Discussion

In this study, we provide the single-cell transcriptional landscape of 4- to 6-week human embryos, which is the critical period of early organogenesis when diverse embryonic tissue and cell types appear^1^. Combining with spatial transcriptome, the locations of most cell types are resolved. As demonstrated in the case studies, our study provides an opportunity to further our understanding of the developmental schemes of vertebrate embryogenesis and the molecular mechanisms that drive the specification of organs and the founding cell types. In developmental systems where our current understanding lacks depth, such as the head mesoderm, the cell types defined in our study with spatial and molecular architecture resolved are particularly valuable. Subsequent single-cell characterization of these systems at slightly later stages will bring in the lineage context to further understand the significance of these cell types. In the meantime, the systematic nature of the dataset also provides the opportunity to gain global views that are not obvious from studying individual developmental processes or organs, such as temporal gene regulation on whole-embryo level by *LIN28A* that is conserved in vertebrates.

Powerful as it may be, scRNA-seq relies on cell dissociation during which the spatial information is lost. However, in some cases the spatial information may be inferred from gene expression profiles^88^. In our studies, we successfully reconstructed the spatial axes and domains of the developing neural tube and limb bud. The reconstructed AP axis in neural tube is independently verified by spatial transcriptome. We reason that such success reflects the developmental mechanisms of axes specification that generate a gradient of cell differentiation states at the initial stage of axes formation, such as molecular gradients and the coupling of time and space through a differentiation front in limb bud development. Since axis specification is prominent at this stage of embryogenesis and occurs in many organs and developmental systems, systematic examination of gene expression gradients will not only enrich our understanding of the molecular mechanisms of axis specification and pattern formation across the embryo, but also help focus future efforts of spatial transcriptomics^13^.

With the collection of whole embryo datasets in vertebrates, cross species analysis becomes increasingly powerful. In one such analysis, we showed that vertebrate embryogenesis consists of 4 conserved developmental stages with sharp temporal boundaries, which implies an unexpected concept that vertebrate embryogenesis may require systemic regulation of stage transition. LIN28A appears to regulate stage-specific growth through direct binding of target mRNAs. Further studies are needed to examine potential heterochronic phenotypes of differentiation across organs and tissues. Besides systemic insights, detailed comparison of individual organs and cell types can reveal potential human-specific regulation, such as the human-mouse differences in the patterning of the neural tube and chondrogenesis in the limb bud.

Accumulation of single-cell studies and data integration are key toward consortium efforts such as the human cell atlas^89, 90^. In terms of human development, our data covers a critical period. As demonstrated in the developing retina, by defining the founding cell population and early progenitor cell types of organs, our data provide the root of organ-specific differentiation trajectories for most of the organs. Meanwhile, our attempt for systematic integration with fetal organ data from 10 to 26 weeks suggest that weeks 6 to 10 is a critical gap to be filled in terms of single-cell omics to achieve informative trajectories of lineage differentiation systematically. In contrast, clinical specimens for stages prior to 4 weeks are extremely rare^91^, and our best hope in the foreseeable future to study the earliest stages may be the *in vitro* models that are being perfected such as the organoids and gastruloids^92^. In this regard, our dataset provides the *in vivo* benchmark to evaluate how well different models mimic development: the end product of such a model should match the *in vivo* cell types. Similarly, our dataset provides valuable guidance for iPS and ESC-based stem cell engineering for regenerative medicine.

## Acknowledgements

We thank Drs. Rachna Sheth, Hongbo Zhang, Sarah Teichmann, Chaolin Zhang, Danwei Huangfu, Alexandra Joyner, Lorenz Studer, and Gufa Lin for discussions. We thank Leo Xiao for website design. This work is supported by Funding Project of National Key Research and Development Program of China (2018YFD0900604), Natural Science Foundation of China (41676119 and 41476120), and Fundamental Research Funds for the Central Universities from the Ocean University of China to W.S.. Z.B. is supported by funding from Memorial Sloan Kettering Cancer Center. T.Z. is a visiting student at Memorial Sloan Kettering Cancer Center sponsored by the China Scholarship Council.

## Author Contributions

W.S. and Z.B. conceived the project. T.Z., M.H., Y.Q., L.W., Q.Z., and Y.X. collected the embryos and generated the atlas dataset. Y.X., T.Z. and Y.N. performed data processing, clustering, and all the bioinformatics analysis. Y.X. created the atlas website. W.S., Z.B., Y.X., and T.Z. wrote the manuscript.

## Methods

### Ethics statement

This study of single-cell transcriptome analysis of human embryos between 4 to 6 weeks was approved by the Ethics Committee of Tongji University School of Medicine and Life Sciences (No. 2019tjdx280) and the Medical Ethics Committee of Traditional Chinese Medicine Hospital of Kunshan (No. 2019-17). The collection and use of human embryos are compliant to current ISSCR guidelines. Informed consent was agreed and signed by each patient only after each patient was provided with all the necessary information about the study. Embryos were collected only after voluntary informed consent was obtained from each patient undergoing legal pregnancy termination. All procedures were performed in strict accordance with the ‘Management of Human Genetic Resources’ formulated by the Ministry of Science and Technology of China (No. 717, effective July 1, 2019).

### Embryo collection and sequencing

Aborted embryos from donations were washed with PBS and immediately transported to the laboratory in cold Hibernate-E medium (Gibco). Embryos with good quality were staged based on anatomical characters^93^. Seven embryos were collected for scRNA-seq with 1 embryo at CS12, 3 at CS13-14, and 3 at CS15-16. We dissected each embryo on ice into four main parts, *i.e.*, head, trunk, viscera, and limb except for Emb. 01 of CS12. Considering the small size of Emb. 01, we dissected it into two parts at the transverse plane, head and upper trunk as one part (ht), lower trunk and viscera as another part (tv) (Extended Data Fig. 1A). The dissection parts of embryo were digested with 0.05mg/ml Liberase TM (Roche) in Hibernate-E medium at 37°C for 10 minutes and stopped with FBS. Single cells were subsequently washed twice with cold PBS, filtered through a 35 μm cell strainer and loaded onto 10x Genomics Chromium system using Single Cell 3’ Reagent Kits v2 (Emb. 01-02, Emb. 05-07) or v3 (Emb. 03-04) at a concentration of 16,000 single cells per sample. Head sample of Emb. 06 was made into two libraries as technical replicates. For all other dissection parts, each was made as one library. Libraries were sequenced on Illumina NovaSeq system in PE150 mode.

For spatial transcriptome, a CS13 embryo was embedded in optimal cutting temperature compound. The embryo was cut into 10 μm sections through the sagittal planes. One section at the center and another section off the center about 1/4 embryo thickness to the edge were chosen. Two sections were processed using Visium Spatial Gene Expression Kit (10x Genomics) according to the manufacturer’s instructions. First, embryonic sections were permeabilized with 12 mins that is optimized by Visium Spatial Tissue Optimization Kit. Second, sections were stained with hematoxylin and eosin (H&E) and imaged using PANNORAMIC MIDI scanner. Then, reverse transcription, spatially barcoding, and cDNA amplification were performed on sections. The resulted libraries were examined for quality control and sequenced on Illumina NovaSeq system in PE150 mode.

### Cell type identification

#### 1. Collection of ontology and diagnostic markers

The quality of annotation of cell types is the key for a valuable single-cell database. Towards a high-quality annotation, we need precise and comprehensive ontology and diagnostic markers. Two principles were considered during the literature search. First, the ontology should follow a hierarchical structure from developmental systems (organs, tissues and major sublineages) to cell types to capture the known lineage hierarchy. Second, the developmental stage of the diagnostic markers should match the time window we study.

Because studies on human embryos from CS12-CS16 are scarce, we expanded our search to studies in mouse embryos at the corresponding stages (E9.5-E11.5)^94^. In total, 157 references for developmental systems and cell types were collected, which describe their organization on lineage/anatomy (ontology), as well as 234 diagnostic markers for developmental systems and cell types (Supplementary Table 1). Diagnostic markers were defined as genes with specific expression on global (developmental systems) or local (cell types) scale at the time window we studied.

#### 2. Knowledge-based semi-supervised clustering

##### 2.1 Overview

Embryos in our study are at the beginning of organogenesis with a dramatic increase of cell types. To handle such a complex system, we designed an iterative clustering approach to first resolve the developmental systems and then cluster cell types within each system. Some developmental systems do not have well-documented ontology information to be referenced, such as splanchnic LPM. To cover known cell types and allow for searching unknown cell types, we built a semi-supervised approach that applied knowledge to guide and optimize the iterative clustering. This approach combines the advantage of unsupervised clustering^2^ and knowledge-guided identification^62^. Details of the approach follow.

##### 2.2 Preprocessing of scRNA-seq data

###### 2.2.1 The determination of sex of embryos

The expression of *XIST* and *RSP4Y1* were used to determine the sex of embryos. Embryos with high *XIST* expression and no *RSP4Y1* expression were defined as female, and *vice versa*, as male.

###### 2.2.2 Quality control on cells

Raw reads were demultiplexed, filtered and mapped to human reference genome (GRCh38) using Cell Ranger 3.0.2 with default settings. The filtered feature-barcode matrices generated by Cell Ranger were used for downstream analysis. Four libraries were considered as outliers and the entire library was excluded from downstream analysis because of low number of detected genes (head sample of Emb. 03 and head, trunk samples of Emb. 07) or contamination of erythroid cells (>80% erythroid cells in viscera sample of Emb. 06). To exclude low-quality cells, cells with fewer than 1,000 genes or over the 90th quantile of total UMIs in each library or having higher than 10% UMIs from mitochondrial genome were discarded. The doublets were estimated by scrublet pipeline with ‘expected_doublet_rate’=0.1 and thresholding at scrublet score 0.4 according to the bimodal distribution of score^95^. In total, 41% of filtered cells by Cell Ranger were considered as low-quality cells or doublets and excluded from downstream analysis. The expression matrix was normalized into UMIs per 10,000 UMIs.

###### 2.2.3 Isolation of erythroid cells

We observed that erythroid cells are the major source of lysis contamination. To prevent potential influence of erythroid cells on downstream analysis, we identified and isolated them using canonical markers before clustering. To do that, the total UMIs of hemoglobin genes (*HBA1*, *HBA2*, *HBE1*, *HBG1*, *HBG2*, *HBZ*) were plotted against total UMIs in each cell (Extended Data Fig. 1D). The threshold of erythroid cells was set as the upper boundary of 99% confidence interval of linear regression on cells with low UMIs of hemoglobin genes (Extended Data Fig. 1D, dash green line). The cells with total UMIs of hemoglobin genes above the threshold were considered as erythroid cells (Extended Data Fig. 1D, red dots) and excluded from clustering. A total of 1,000 randomly selected red blood cells were included for the downstream analysis after clustering.

##### 2.3 Identification of developmental systems

Cells from all embryos were pooled before clustering. Aggregate information from pooled samples increases power of clustering especially for cell types with low differentiated status^96^. As shown in the section of quality control, the temporal difference within a cell type does not have a profound effect on the identification of cell types (except limb mesenchyme, see section of *Special cases* below).

The top level of clustering is aimed to identify super-clusters (we termed super-cluster as a cluster resolved from top-level of clustering) at the level of developmental system. To reduce batch effect between samples, the detection of HVGs were performed per sample based on Poisson distribution^96^ and then merged from each sample. TFs in HVGs were used in Louvain clustering in R package Seurat 3.0^97^ (‘FindClusters’, r=0.5, PCs=30). Batch correction was performed by reciprocal PCA with two batches (v2 kit: Emb. 01-02, Emb. 05-07; v3 kit: Emb. 03-04). Stripped nucleus removal was performed as described in Pijuan-Sala *et al.* ^5^. 25,040 cells from three clusters were found to have considerably lower mitochondrial gene expression (total UMIs of 13 mitochondrial protein-coding genes 58±70 in them vs. 179±133 in other cells, mean ± standard deviation). A total of 185,140 high-quality cells were obtained after removing presumable stripped nuclei. Sixteen super-clusters were annotated with canonical markers in the clustering of high-quality cells (Extended Data Fig. 2A).

Only TFs in HVGs were used in the top level of clustering because we found TFs in HVGs are better in preserving lineage organization than all HVGs (Extended Data Fig. 2). The distribution of expression of canonical markers across super-clusters show that markers of intermediate mesoderm (*WT1*) and endoderm (*FOXA2*) significantly lose their specificity in the clustering by HVGs compared to that by TFs (Kolmogorov-Smirnov test p < 0.05). The global mapping between two sets of super-clusters shows that super-clusters 3 and 7-12 by HVGs do not have clear match to super-clusters by TFs (Extended Data Fig. 2C). Super-cluster 3 by HVGs is composed of cells from many super-clusters by TFs, including almost all the *FOXA2*+ endoderm (Extended Data Fig. 2C). The non-TF HVGs shared in super-cluster 3 are likely the common feature of epithelium, causing the mixture of epithelial cells from different germ layers in super-cluster 3. Furthermore, super-clusters 10-12 by HVGs are strongly biased to embryonic stage (Extended Data Fig. 2D), which leads to the mismatch on mesodermal super-clusters. The mismatch on mesodermal super-clusters is also observed in the clustering with downsampling of HVGs to the number of TFs in HVGs (Extended Data Fig. 2E).

##### 2.4 Identification of cell types

The low level of clustering is aimed to identify clusters at the level of cell type, which is performed within each super-cluster.

###### 2.4.1 HVGs

HVGs were calculated per sample based on Poisson distribution^96^ and then merged from each sample. Ubiquitously expressed HVGs in each sample (>70% cells) were excluded. Also, hemoglobin genes, cell cycle genes^98, 99^, sex-specific genes (*XIST*, *RPS4Y1*, *RPS4Y2*), and batch-effect genes^13,100^ were excluded from HVGs during clustering.

###### 2.4.2.1 Rounds of clustering

The rounds of clustering were controlled so that known cell types were well-resolved according to diagnostic markers and clustering is not too fragile. For relatively complex super-clusters, i.e., brain, neural progenitor, neuron, craniofacial, splanchnic LPM, and endoderm, we performed two rounds of clustering. For other super-clusters, one round of clustering was performed. We considered whether the second round of clustering is needed according to the support of divisions of the second round by diagnostic markers (Extended Data Fig. 4A-B). For example, in brain, 9 clusters were derived from first round of clustering, each of which was further divided into 1-4 clusters in the second round of clustering. All the divisions in the second round were supported by 2 or more markers. In comparison, in endothelium, only 1 cluster from the first round was supported to be further divided. Thus, two rounds of clustering were chosen for brain and one round of clustering were chosen for endothelium.

###### 2.4.2.2 Parameters in Louvain clustering

The resolution ‘r’ in ‘FindClusters’ (Seurat) were determined by systems with more prior knowledge and then applied to systems with less prior knowledge. The ‘r’ was initially trained in spinal neuron, which is composed of 12 well-studied cell types. A list of ‘r’ value spanning from 0.1 to 1 with interval of 0.2 were examined to see which one gives the closest number of clusters to expected number (Extended Data Fig. 4C). Under the condition of 10, 20, and 30 PCs, the number of clusters resulted from r = 0.5 is closest to the expected number 12. The clusters start to be convergent when r > 0.5 (Extended Data Fig. 4D, adjusted Rand index, ARI). As a *post hoc* examination, known cell types in blood were resolved under this resolution (Supplementary Note 1). Thus, r = 0.5 was propagated to the clustering of other super-clusters except special cases. The number of PCs were determined by ranking of PCs based on the percentage of variance explained by each one (elbow point, PCs = 20).

The robustness of clustering was tested by comparing the result of the chosen r or PCs to results from a range of values in each super-cluster (Extended Data Fig. 4E), which is measured by ARI. PC = 20 was compared to a series of PCs (15-25, increment 1, excluding 20) when r was fixed to 0.5. Resolution ‘r’ = 0.5 was compared to a series of ‘r’ (0.3-0.7, increment 0.1, excluding 0.5) when PCs was fixed to 20.

###### 2.4.3 Special cases: untangling axes in the identification of cell types

During clustering, we found different developmental systems may have different profound factors (or axes) entangling upon the axis of cell type. Axis formation is very common among organs at this stage. These axes are reflected in the single-cell data and sometimes orthogonal to the axis of cell type. They will not be sequentially decomposed by clustering, which may cause mistakes on cell type identification in blind clustering. Thus, untangling axes properly could improve the correctness of cell type identification. This section will introduce two cases we handled, which can be implication for other studies with similar characteristics.

###### 2.4.3.1 Temporal axis in limb bud

Temporal axis is one of profound axes in the mesenchyme of limb bud. We found removal of temporal axis improved cell type identification at CS12 in limb bud due to profound axes and disproportional cell number between stages. Limb bud is fast growing in the time window of our samples, during which anterior-posterior (AP) axis, proximal-distal (PD) axis, and differentiation axis are tangled, such as, the change of HOX genes expression on both AP/PD axes and the start of chondrogenesis at CS15-16 (Fig. 3). The limb cells harvested at later stages are 10 folds more than cells at CS12. It is likely that signals from the relatively low number of cells at CS12 were overshadowed by other stages in pooled clustering. Thus, clustering in the super-cluster limb was performed per stage.

###### 2.4.3.2 HOX genes

Cell types running along the AP axis adapt HOX code, which is supposed to make cell type dissimilar by AP position. To untangle it, we excluded HOX genes in HVGs for developmental systems running along AP axis, i.e., neural progenitor, neuron, Schwann, somites, IM, and LPM. Furthermore, for systems with known AP segmentation (neural progenitor and neuron), HOX genes were included at next round of clustering to distinguish neural cell types between rhombomere and spinal cord, e.g., dA1 in rhombomere and dl1 in spinal cord. In this way, distinguished segmentations on AP axis can be identified.

#### 3. Annotation

##### 3.1 Annotation of cell types

The biological identity of each cluster was manually annotated with 2-5 marker genes (Supplementary Table 1B). Further detail can be found in Supplementary Note 1 “annotation_vignette.pdf” using blood as an example. For all other systems, the detail can be found at https://heoa.shinyapps.io/code/. In total, we obtained 313 cell types.

###### 3.2 Post-processing

To organize resulted cell types in reflecting lineage history, we regrouped 16 super-clusters into 18 developmental systems (Fig. 1E). Cell types in super-cluster brain at this stage are all neural progenitors, and thus were merged with the neural progenitor super-cluster. Sensory neurons have different embryonic origin with other neurons in spinal cord and were separated as a new system “sensory neuron”. Cell types in super-cluster heart were merged with splanchnic LPM. Because of few number of cells, PGC was not resolved as a cluster. We identified presumable PGC cells directly by markers (Supplementary Table 1A, *POU5F1*, *DDX4*, *DPPA3, DAZL*, *NANO3*, *DND1*, *ALPL*, *SALL4*, total UMIs >= 9, total detected markers >= 3 in trunk and viscera samples) as a new system “PGC”. Cell types with epithelium feature (*KRT18/19*, *CLDN4*) but without concrete identity of any system were regrouped as a new system “epithelium”. Cell types with fibroblast feature (*COL1A2*, *COL3A1*) but without concrete identity of any system were regrouped as “fibroblast”. Ten cell types without clear belonging of developmental system were regrouped as “miscellaneous”.

##### 4. Quality control on cell types

###### 4.1 Cross-validation

Cross validation on cell identity at multiple levels was performed in R package scPred^20^. For the identity of developmental system, 2,000 randomly sampled cells in each system were used to train a classifier if the number of cells in this system is greater than 2,000. Within each developmental system, both cell type annotation (“Celltype_annotation” in Supplementary Table 1C) and final identity (“Final_annotation” in Supplementary Table 1C) were trained and tested (red and blue dots in Extended Data Fig. 4F, respectively). Developmental system PGC is not included because there is only 1 cell type in this system. In each testing, a Seurat object was constructed using the corresponding cells and setting as that in clustering, which is served as the input of scPred. As negative control, randomly shuffled system identity or final identity was tested in the same pipeline. All results are evaluated by AUROC (area under receiver operating curve).

###### 4.2 Estimation of batch effect

Batch effects on clustering were estimated by the entropy of mixing^101^. Four types of batch effects were tested: embryo, technical replicates (Emb. 06 head, see Methods section “embryo collection”), cell cycle phase (from Seurat “CellCycleScoring”), and total UMIs (stratified by 4 quantiles). For each test, 100 cells were randomly sampled from each developmental system and the labels of 100 nearest neighbors of each sampled cell were scored in the reduced-dimension space as mixing entropy^101^. The entropy of mixing was calculated under positive control (cluster identity as label), this batch effect (batch as label), and negative control (random label), respectively (Extended Data Fig. 4G).

###### 4.3 Comparison with snRNA-seq of whole mouse embryos

We compared our human dataset (except cell types in miscellaneous) to scRNA-seq data in whole mouse embryos^2^. First, 644 cell types out of total 655 cell types from mouse dataset were taken, which contain cells from E9.5-E11.5, corresponding to the stages of our human dataset. Second, the match of cell types between human and mouse were calculated by non-negative least squares (NNLS) regression based approach from Cao *et al.* ^2^ with slight changes. Briefly, a set of informative genes was selected for NNLS across all comparisons between cell types, which are the union of DEGs of human cell types excluding genes that are expressed in >50% human or mouse cell types. The mean expression of the gene set in each human cell type was predicted by NNLS with the gene expression of all mouse cell types. Reciprocally, the mean expression of the gene set in each mouse cell type was predicted by NNLS with the gene expression of all human cell types. The specificity score is defined as beta1 + beta2, which were resulted from two NNLS regressions, respectively. Third, to filter matched mouse cell types, we retained top 4 mouse cell types if more than 4 are best matched to the same human cell type. We noticed that neural cell types are overrepresented in mouse dataset (308 out of 644 cell types), we removed neural cell types that are best matched to mesodermal cell type in human. We removed mouse cell types that have specificity score > 0.1 with > 20 human cell types. Three steps of filtering resulted 298 mouse cell types, which is comparable to the number of cell types in human dataset (Extended Data Fig. 5, Supplementary Table 2A). Final, to evaluate the comparison, we determined the developmental system of mouse cell type according to annotation from Cao *et al.* ^2^ and compared to the developmental system of its best matched human cell type. Mesodermal cell types in mouse that can not be further specified to which mesodermal system were classified as mesoderm. For number of human cell types with matched mouse cell type, the threshold of specificity score is 0.05.

##### 5. Identification of DEGs

The traditional way of identifying DEGs by comparing one cell type versus all other cell types may not be appropriate here considering large number of cell types and most DEGs are not specific to only one cell type. We considered a gene as a DEG if it is not expressed in many cell types and is highly expressed in at least one cell type. First, genes with mean of normalized UMIs < 0.3 in more than 20% of cell types were selected. Second, the DEGs for a given cell type were defined as genes with high expression in the given cell type (detected in >= 25% of cells, mean of normalized UMIs >= 0.5, and z-score of mean of normalized UMIs >= 7 with non-expressing types as background). A *post hoc* test by Mann-Whitney U test showed that adjusted p values of identified DEGs are less than 0.005.

To exclude genes associated with lysis contamination from DEGs, for each cell type of hepatocyte, cardiomyocyte, and erythroid cells, we considered a gene could cause lysis contamination if it is a DEG of this cell type, has highest expression in this type, and mean of normalized UMIs > 2 in this type, which resulted 132 genes in total. The 132 genes were removed from DEGs of all cell types except the source cell type.

#### Analysis of ligands in signaling centers

Ligands were collected from 3,252 pairs of curated ligand-receptor interactions^21^. Ligands that are DEGs of at least one signaling center were selected and divided into two groups (Extended Data Fig. 6D). To compare ligand expression between human and mouse, gene ortholog relationship was downloaded from Ensembl (Ensembl Genes 100). Expression data of 9 signaling centers (ACP, MHB, ZLI, FP.b, RP.b^27^; FP.s, RP.s^25^; AER, ZPA^26^) in mouse at the corresponding stage were collected from three scRNA-seq datasets. For the seven signaling centers in nervous system, cells were obtained according to the annotation from original paper. Reclustering were performed to obtain cells of ZPA and AER (see Methods “spatial domain in limb” and Supplementary Table 2). All expression in mouse data were normalized to UMIs per 10,000. The ligands are considered expressing in mouse cell type at threshold of normalized UMIs >= 0.5.

#### Process of spatial transcriptome (ST)

##### 1. Preprocessing and quality control

Raw data were mapped to human genome (GRCh38) and filtered by Space Ranger 1.2.0 with default settings. Gene expression in each spot were normalized to UMIs per 10,000 UMIs. To assess the quality of spatial transcriptome, we compared our data to Stereo-seq in mouse embryos at E10.5^28^ using unsupervised clustering. Raw data of two sections (E1S2 and E2S1) of mouse embryos were downloaded and merged into transcriptome per 50 μm (diameter) spot (75×75 nanoballs), which is comparable to the resolution of our human data. Two human sections and two mouse sections were integrated (reciprocal PCA with top 1000 HVGs, only 1-on-1 orthologs were considered) and clustered (Louvain algorithm with 30 PCs) by Seurat at the level of spot. To examine whether human and mouse spots from the same cluster locate at the same anatomical region, we calculated the correlation of the spatial neighborhood for each cluster between human and mouse. The spatial neighborhood for a cluster on a section was defined as the proportions of clusters in its neighbors. The neighbors of a cluster were defined as the union of nearest 6 spots on the section of each spot in this cluster, excluding spots from the same cluster. We noticed that mouse section E2S1 is mostly mesodermal tissue in trunk and E1S2 is mostly neural tissue in trunk, thus section E2S1 was compared to human section 1 and section E1S2 is compared to human section 2.

##### 2. Deconvolution

Spatial transcriptome was deconvolved by 239 cell types from scRNA-seq with RCTD^29^ under multiple mode, which are all cell types excluding these from limb (not on any section), epithelium, fibroblast, and miscellaneous. For each spot, the percentages of cell types in the confident list reported by RCTD were considered as the proportion of cell types in this spot. Other cell types were considered as not available in the spot. The deconvolution process is effective as confident calls were reported by RCTD in 96% of spots. The deconvolution results of all cell types on two sections are in Supplementary Note 2 and 3. The proportion of developmental system in each spot was defined as the sum of proportion of cell types from each developmental system. For head mesoderm, undefined cell types having more than 20 spots with proportion > 0.05 on section 1 were shown in Fig. 2E.

##### 3. Signaling interaction

We designed a pipeline to infer signaling interaction using scRNA-seq and spatial transcriptome (Extended Data Fig. 8E). First, we filtered cell type pairs that have colocalization on any section (>= 5 spots with proportions of both cell types > 0.2). Cell types of blood and endothelium were not considered. For colocalized cell type pairs, we search for ligand-receptor pairs^21^ in which ligand and receptor are expressed in two cell types in scRNA-seq, respectively (mean of normalized UMIs > 0.5). Second, the enrichment of co-expression of ligand and receptor (normalized UMIs > 0.5) in the spots with colocalization of cell type pair was examined by hypergeometric test, as compared to total number of spots with co-expression of ligand and receptor on the section. A significant interaction was defined as Benjamini & Hochberg adjusted p value < 0.01 in at least one section. Two sections were processed separately and significant interactions were pooled in Supplementary Table 1G by taking the minimal adjusted p value from two sections for a ligand-receptor pair in a cell type pair.

#### Spatial domains in limb development

*In situ* data on limb staged from E9.5 to E11.5 mouse were collected from literatures and the EMBRYS database^37–46^ (Extended Data Fig. 10-11). The spatial location of cell types of forelimb was determined by comparing expression of DEGs between scRNA-seq and *in situ* data. Limb domains at three stages were summarized as schematic diagrams. Please see Supplementary Note 1 “annotation_vignette.pdf” for the detail of annotation process.

Single-cell RNA-seq data in mouse limb were downloaded from ENCODE project^26^. To compare with spatial domains between human and mouse, mouse mesenchymal cells (cell types annotated with “mesenchymal 1”, “mesenchymal 2”, “perichondrial”, “chondrocyte”, “foxp1+ perichondrial”) at E10.5 and E12 were reclustered and annotated in the same way with human data (Extended Data Fig. 9E and Supplementary Table 2). To obtain AER of mouse limb, mouse cells annotated with “epithelial 1” were reclustered and a cluster with specific expression of *Fgf8* was identified as AER (Supplementary Table 2).

The Umap co-embedding of limb mesenchyme of CS15-16 in human and E12 in mouse was performed by reciprocal PCA in Seurat (Extended Data Fig. 9H, 1-on-1 orthologs in HVGs, 20 PCs). Anchor pairs between chondrocyte and non-chondrocyte were removed in reciprocal PCA. To examine the corresponding mouse cells for human cells, we calculate the percentage of mouse cells in k-nearest neighbors (k=6) of each human cell in co-embedding space. To control the size factor, for each human domain, we randomly sampled 50 cells that are within mean + 1 standard deviation (SD) to the centroid of the given domain to examine neighboring mouse cells. The percentage of mouse cells in total cells of human and mouse was defined as the background value (43%). Mann-Whitney U test followed by Benjamini & Hochberg correction was applied to compare the percentage of mouse cells in nearest neighbors of each human domain to the background.

#### Neural tube patterning

Neural progenitors from the hindbrain and the spinal cord in scRNA-seq were used for pseudo-spatial analysis. All expressed HOX genes and 5 housekeeping genes (*RPL5*, *ELAVL1*, *ATP2C1*, *ARPC2*, *ARPC1A*, to prevent omitting cells with zero HOX gene expression) were used to reconstruct the pseudo-axis using Monocle^102^. We then defined a group of AP-related genes by calculating the correlation between gene expression and cell ordering on the pseudo AP axis. Genes with notably positive (4×SD) or negative (-4×SD) correlation were considered as AP-related genes.

To define AP axis on section 2 of ST, a midline was manually drawn in the spots on the dorsal trunk with proportion of neural cell types > 0.8 in deconvolution. An AP value was given in a decreasing order for each spot on the midline. For spots in the dorsal trunk that are not on the midline, the AP value was given to the same value with its nearest spot on the midline (Fig. 4C). The expression of a gene along AP axis in ST was calculated in three steps: 1) average spots with same AP value in ST; 2) smooth with a window size of 30 along AP axis; 3) normalize to 0-1 range. In Fig. 4C, to calculated Pearson’s correlation of lncRNA between scRNA-seq and ST, cells from CS13-14 in scRNA-seq were filtered and aligned to AP axis of ST by their ordering on pseudo axis. In Extended Data Fig. 12A, Pearson’s correlation between lncRNAs and HOX genes along AP axis of ST were calculated. HOX genes were grouped by number, e.g., HOX2 is from the average of *HOXA2* and *HOXB2*.

For patterning along the dorsal-ventral axis, cells were visualized by UMAP with HVGs excluding HOX genes. The same method was applied on neuron cells from the hindbrain and the spinal cord. To compare gene expression in neural types between human and mouse, we collected single-cell RNA-seq data in neural tube in mouse^62^ for comparison and data in neural tube in human^63^ for cross-validation. Raw UMI matrix were normalized into UMIs per 10,000 UMIs. The canonical markers of neuronal cell types of these two species were collected from Delile *et al.* ^62^.

#### LIN28A and embryogenesis in vertebrates

##### 1. RNA-seq data and preprocessing

RNA-seq in other vertebrates were downloaded from published data (Supplementary Table 3). Bulk RNA-seq were normalized into transcripts per million reads (TPM). Single-cell RNA-seq were normalized into UMIs per 10,000 UMIs.

##### 2. Defining embryonic stages upon timepoints

For each species of zebrafish, frog, and mouse, the pairwise Pearson’s correlation between samples were calculated based on the expression of TFs (Gene Ontology term GO:0003700). Clustering of time points was based on pairwise correlation between samples (R package dendextend, k=4). Considering the time series of mouse data are from three sources using different techniques (E2.5-E5.5 using bulk RNA-seq, E6.5-E8.5 using scRNA-seq, and E9.5-E13.5 using sci-RNA-seq for nuclear-RNA sequencing), correlation calculation and clustering were performed within each experiment. Single-cell RNA-seq at E6.5-E8.5 and E9.5-E13.5 were treated as pseudo bulk-seq for correlation analysis by averaging all cells. Similarity between zebrafish and frog, zebrafish and mouse, frog and mouse samples were measured by pairwise correlation based on the expression of homologous TFs. Considering the batch effect among mouse datasets, inter-species correlations at each stage were converted to z-score by row. The ortholog relationship was downloaded from Ensembl (Ensembl Genes 100). Dynamic time warping was performed to align inter-species stages based on their correlations (R package dtw).

##### 3. Detection of systemically changing genes

For systemically changing genes from stage 1 to stage 2, we compared transcriptome of zebrafish (6-8 hpf vs. 3-4.3 hpf), frog (stages 13-14 vs. stages 6-8), and mouse (E4.5-5.5 vs. E2.5-3.5). No human transcriptome data is available at this stage. From stage 3 to stage 4, we compared transcriptome of zebrafish (3-4 d vs. 10.3-16 hpf), frog (stages 31-34 vs. stages 15-18), mouse (E11.5 vs. E9.5-10.5) and human (CS15-16 vs. CS12-14). For zebrafish and frog, as well as mouse from stage 1 to stage 2, where bulk-seq is available, systemically changing genes were defined as those differentially expressed between stages (> 2 fold) in bulk RNA-seq and are systemically expressed (> 50% cell types) in scRNA-seq at the corresponding stages. For human from stage 3 to stage 4, systemically changing genes were defined as differentially expressed between stages across cell types (> 2 fold in at least 1 cell type and > 1.5 fold in at least 50% cell types). The fold change of cell type was set to 1 if no robust expression was detected in any stage in scRNA-seq (mean < 0.3). Genes with inconsistent change across cell types (> 1.5 fold in at least 1 cell type and < -1.5 fold in at least 1 cell type) were removed. The same definition was applied to systemically changing genes in mouse from stage 3 to stage 4 except the threshold of robust expression is set to 0.01 and fold change is set to 1.3, since the mouse scRNA-seq data in this time window has a tenfold decrease in total UMIs compared to human data. Cell types used in scRNA-seq data were defined as that in the original publication. For human, cell types with clear identity were used and cell types in limb and neural tube were merged (Supplementary Table 1C). To avoid distortion by cell types with low number of cells, we only considered cell types with more than 40 cells at both early and late stages in each species.

##### 4. Pathway enrichment analysis

Pathway enrichment analysis for systemically changing genes was performed against gene sets of Reactome on MSigDB^103^. P values were calculated by hypergeometric test and corrected by Benjamini & Hochberg method. The enriched pathways were defined as pathways with adjusted p value < 10^-4^ in at least 2 species. Pathways that are both enriched from stage 1 to stage 2 and from stage 3 to stage 4 were highlighted in Supplementary Table 5.

#### Data integration

Single-cell RNA-seq data in human embryos between 72 days and 129 days in estimated postconceptual age were downloaded from Cao *et al.* ^7^ (raw UMI matrix of 377,456 cells from https://descartes.brotmanbaty.org/). To compensate the low detected UMIs per cell (median 863) in sci-RNA-seq3, we pooled 5 cells per type per embryo into 1 meta-cell, which resulted median UMIs 2871 per meta-cell (Extended Data Fig. 14A). Two Seurat objects were created with raw UMI matrix of our data and Cao’s data, respectively. Reciprocal PCA in Seurat package was applied to integrate two datasets by top 2,000 HVGs. The joint PCA (30 PCs) is used as the input of dimension reduction by UMAP.

To calculate the similarity between our cell types and cell types at the later stages (Cao’s dataset), we first cleaned cell types in both datasets. 73 cell types in Cao’s dataset were included except 4 cell types from trophoblasts. To balance cell types between two datasets, we regrouped cell types in our data into 72 cell types (*e.g.*, all types of frontonasal mesenchyme were merged as one cell type because it was not sampled in Cao’s dataset), and exclude cell types from epithelium, fibroblast, and miscellaneous (Supplementary Table 1). Slingshot^82^ was used to calculate the pairwise distance between cell types from two datasets in the joint PCA space from Seurat. We considered three types of linkages based on Slingshot distance, mutual best match, best match, and second-best match. First, mutual best matches of cell types between two datasets were taken. Second, for each cell type in Cao’s dataset, the best matched cell type in our dataset was defined as the cell type with minimal distance. For 6 terminal cell types in our dataset (Supplementary Table 1), only 1 best match from Cao’s dataset was allowed. Other cell types in Cao’s dataset best matched to terminal cell types in our dataset were relinked to next best match. Third, for each cell type in Cao’s dataset, the second-best matched cell type in our dataset was defined if the z-score of distance of the cell type with second minimal distance is less than -2. For a cell type in Cao’s dataset with mutual best match, any cell type in our dataset with distance great than 3 fold of the minimal distance was excluded from second-best match. The three types of linkages between two datasets were visualized by R package igraph (layout ‘layout_nicely’). Final, known lineage relationships within our dataset (Supplementary Table 1C) were added onto the graph.

Single-cell RNA-seq data in eye of human embryos (Hu’s dataset)^9^ were collected and cell types of eye in three datasets (our data, Hu’s data, and Cao’s data) were integrated. Three Seurat objects were created with raw UMI matrix of cell types of eye in three datasets, respectively (meta-cells defined above were used for Cao’s dataset). Reciprocal PCA in Seurat package was applied to integrate three datasets by top 2,000 HVGs. The joint PCA (30 PCs) is used as the input of dimension reduction by UMAP. In an ideal trajectory, cells proceed from earlier to later time point. To incorporate temporal information as a constraint, we modeled trajectory reconstruction as minimum span tree problem in a directed graph. First, pairwise distance between cell types were calculated by Slingshot as the weight of edge. Second, cell types were assigned to 4 groups according to their stages (Supplementary Table 6). Third, Chu-Liu/Edmonds’ algorithm^84^ was performed to find minimum spanning tree only allow links from early group to late group (R package optrees, ‘msArborEdmonds’). Pseudotime of cells in RPE and RGC lineage were analyzed in Slingshot (‘slingPseudotime’), respectively. To identify key genes potentially involved in the differentiation of each cell type, we normalized data matrix into UMIs per 10,000 in each dataset. The key genes of a cell type were defined as genes that expressed in this cell type (mean of normalized UMIs >= 0.5 & fraction >= 0.4), upregulated relative to its pseudo-mother in trajectory (> 2 fold), and not upregulated in any of its pseudo-sister (Supplementary Table 6).

#### Data and code availability

The raw and processed data generated in this study can be downloaded from the NCBI Gene Expression Omnibus (GSE157329). An online depository for cell types and gene expression (developed with R package VisCello^104^) is available at https://heoa.shinyapps.io/base/. R Scripts for analysis and figures are available at https://heoa.shinyapps.io/code/.

---

**Extended Data Fig. 1.**
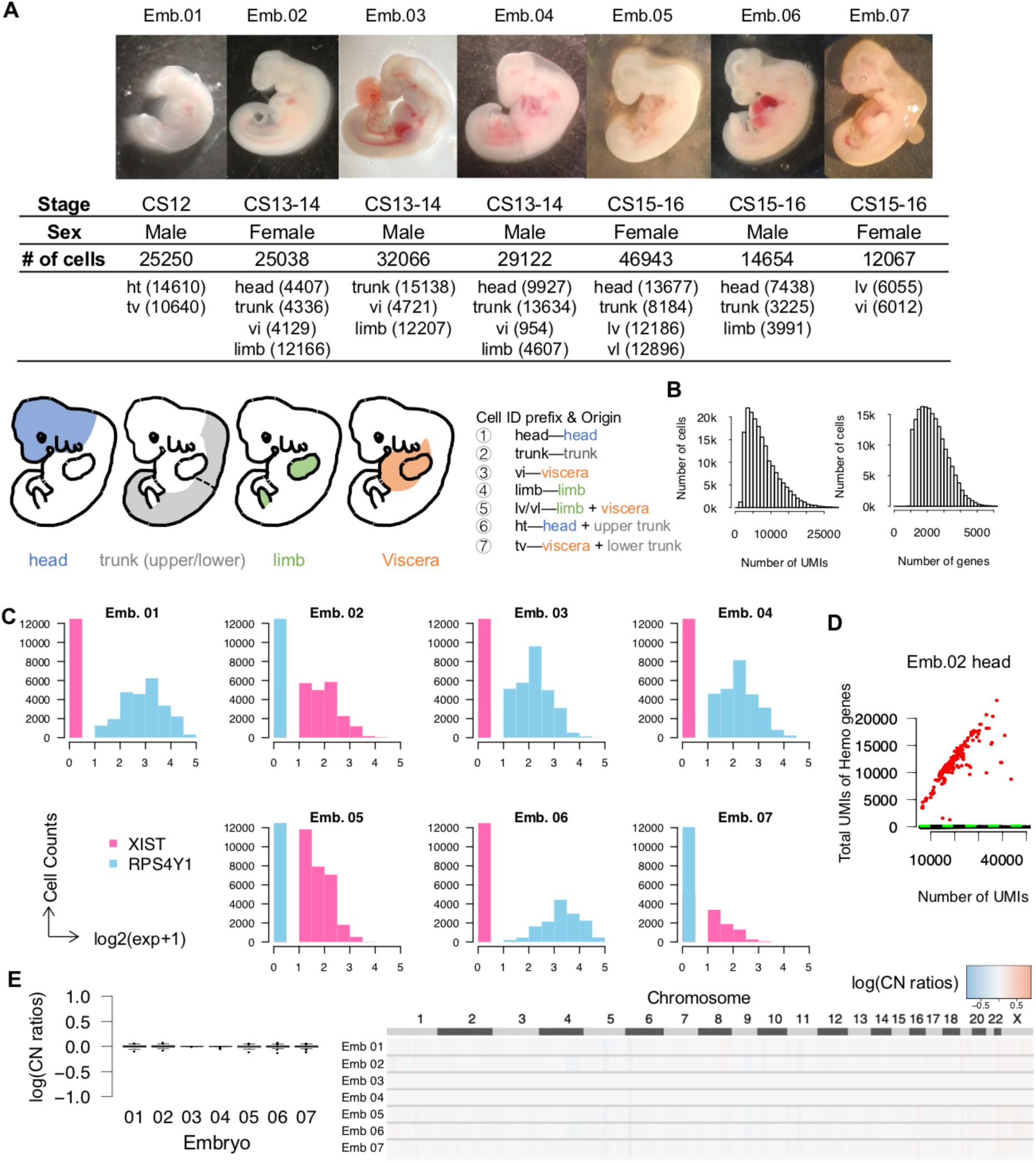
Embryos and data quality. (**A**), Images of embryos and the number of high-quality cells in each sample. Schematic diagrams (lower panel) showing the territory of embryo dissection parts. (**B**), Histograms showing the distribution of total UMIs and gene numbers per cell. (**C**), Distribution of *XIST* (female) versus *RSP4Y1* (male) expression from randomly sampled 1000 cells per embryo. (**D**), UMIs of hemoglobin genes (*HBA1*, *HBA2*, *HBE1*, *HBG1*, *HBG2*, *HBZ*) against total UMIs. Each dot denotes a cell. Cells with high linear slope (above the green line) was identified as erythroid cells (red) (see Methods). Black dots denote cells that were not identified as erythroid cells. (**E**), CopyKAT^18^, designed for 3’ or 5’ scRNA-seq with sparse coverage, was used to estimate copy number aberration in 7 embryos with default setting. 500 cells were randomly sampled in each embryo. Log2 ratios of copy number of segmentations (220 kb of size) were shown by embryo (left) and by genomic position (right).

**Extended Data Fig. 2.**
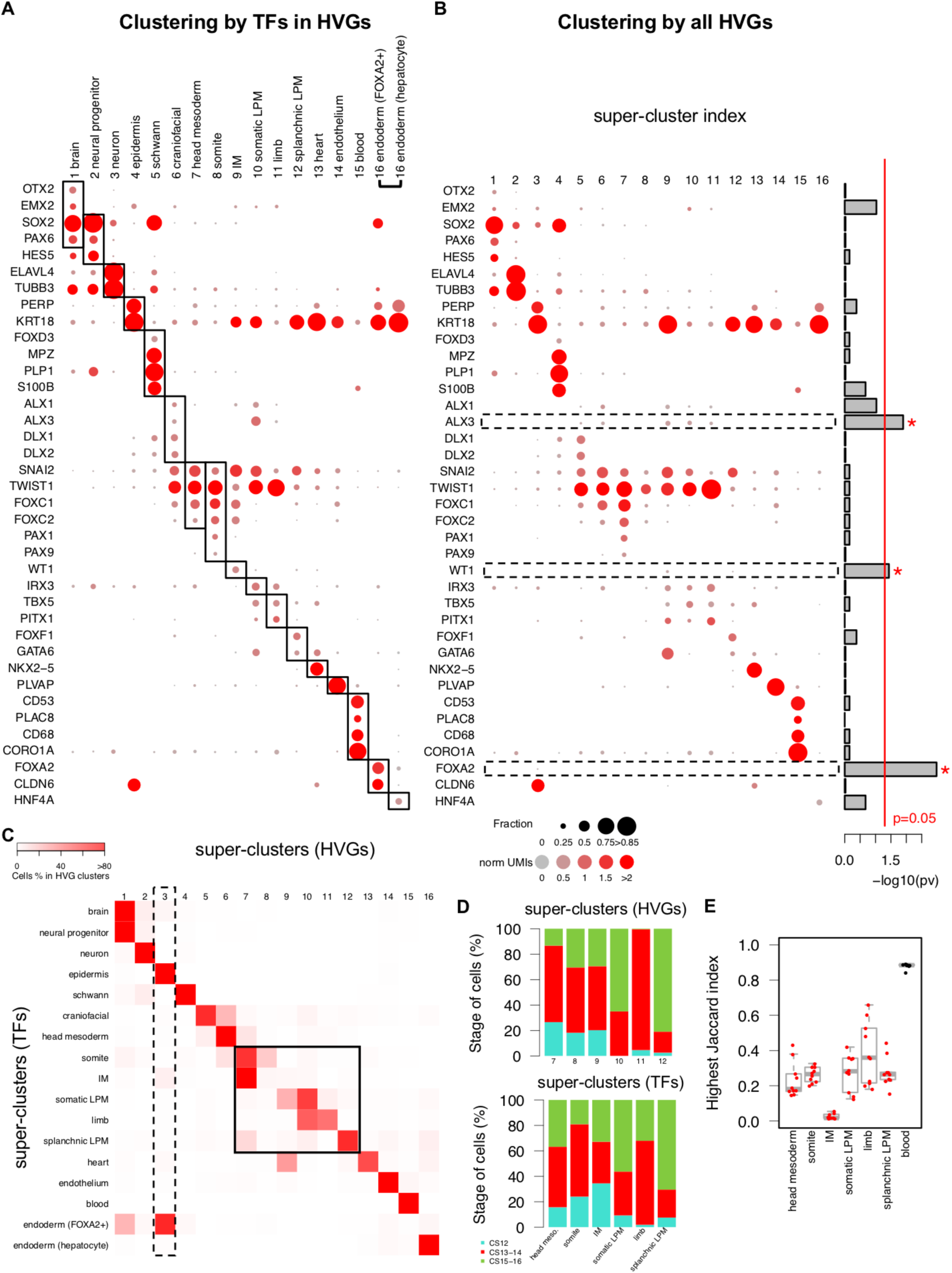
The top level of clustering. (**A**), The expression pattern of marker genes in super-clusters of clustering with TFs in HVGs. The annotations were showed on the column. Because the most representative marker of hepatocyte and other endoderm are different, to better show the specificity of marker genes, endoderm super-cluster was separated into two columns: endoderm (hepatocyte) and other endoderm (*FOXA2*+). (**B**), The expression pattern of marker genes in super-clusters of clustering with all HVGs. Right panel, p values of Kolmogorov-Smirnov test of mean UMIs of marker genes across super-clusters between two sets of clustering. Red line, p = 0.05; boxes, genes with p < 0.05. (**C**), The percentages of cells from super-clusters by TFs (column) for each super-cluster by HVGs (row). Boxes denote super-clusters 3 and 7-12 by HVGs that do not have clear match to super-clusters by TFs. (**D**), The stage distribution of cells in mismatched mesodermal super-clusters. (**E**), The match of mesodermal super-clusters from panel D in the downsampling of HVGs to the same number of TFs in HVGs. In each downsampling, the highest Jaccard index between a mesodermal super-cluster by TFs and all super-clusters by downsampled HVGs were showed as a red dot. The downsampling was performed for 10 times. The match of blood super-cluster by TFs was showed as control (black dot).

**Extended Data Fig. 3.**
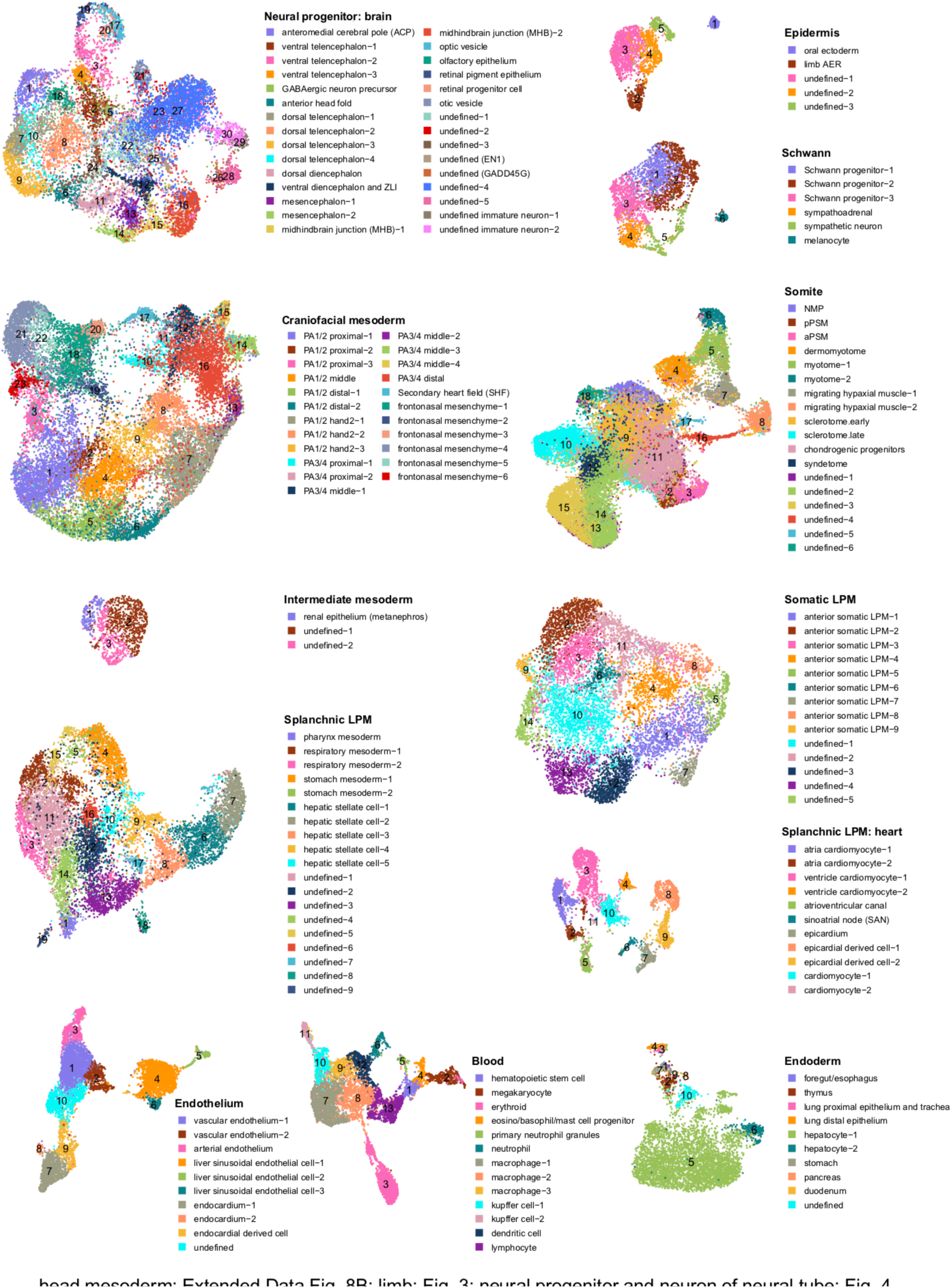
UMAP in each developmental system. To avoid redundancy, 7 developmental systems are not included here, which are head mesoderm (in Extended Data Fig. 8B), limb (in Fig. 3B and Extended Data Fig. 9A), neural progenitor and neuron of neural tube (in Fig. 4D), sensory neuron (in HEOA depository), epithelium (in HEOA depository), fibroblast (in HEOA depository), and PGC (containing one cell type searched by marker genes so that Umap is not available, Methods).

**Extended Data Fig. 4.**
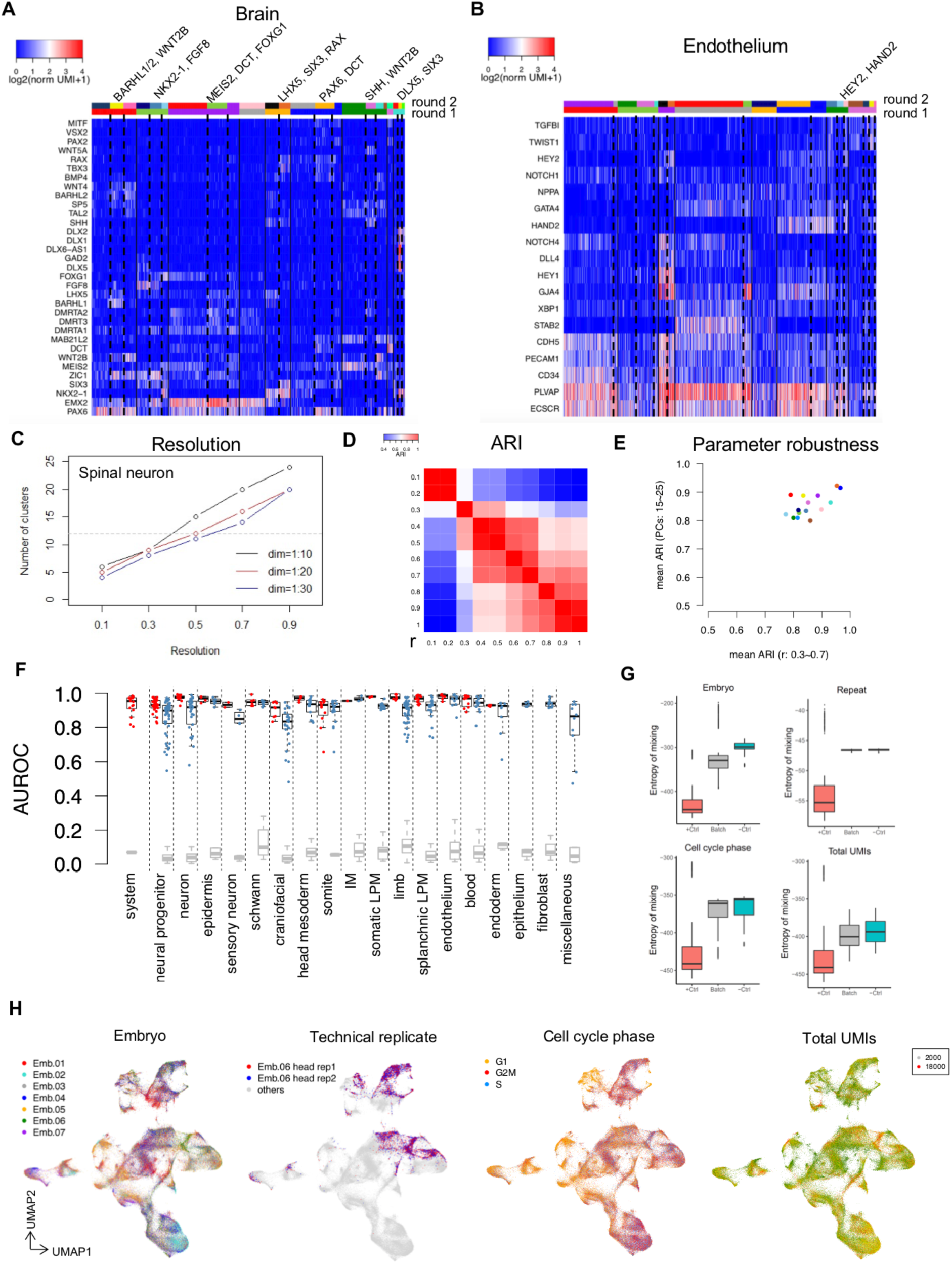
Iterative clustering and quality control. (**A**), The expression of markers relative to boundaries of first round of clustering (solid lines) and second round of clustering (dash lines) in brain. Markers that support the division of second round of clustering were indicated on the top. Two color bars on top denote clustering results of first round and second round, respectively. (**B**), The expression of markers relative to boundaries of one round of clustering and second round of clustering in endothelium. Convention follows panel a. (**C**), The number of clusters resulted from a series of resolution ‘r’ and PCs in the clustering of spinal neuron. (**D**), The pairwise ARI between clusters resulted from different resolution ‘r’ in spinal neuron. (**E**), X-axis, the mean ARI of clusters between chosen ‘r’ (0.5) and a series of ‘r’ (0.3-0.7, increment 0.1, excluding 0.5) when PCs was fixed to 20. Y-axis, the mean ARI of clusters between chosen PCs (20) and a series of PCs (15-25, increment 1, excluding 20) when r was fixed to 0.5. Each dot denotes a super-cluster. (**F**), Cross-validation on clustering by scPred^20^. The first column shows the AUROC (area under receiver operating curve) of testing the identity of developmental systems (each red dot is a system). Other columns show the AUROC of testing “Celltype_annotation” and “Final_annotation” (Supplementary Table 1C) within each system (expect PGC) in red and blue dots, respective (Methods). For systems (epithelium, fibroblast, miscellaneous) that have the same “Celltype_annotation” and “Final_annotation”, only the result of “Final_annotation” is showed. Testing of randomly shuffled identity was served as control in each column (grey). (**G**), Batch effects of embryos, technical replicates, cell cycle phase, and total UMIs estimated by the entropy of mixing^101^ (Methods). Batch effect is anti-correlated to entropy of mixing. Boxplots showing the entropy of mixing using cluster identities as batch (‘+Ctrl’, clustering totally driven by batch effect), entropy of mixing by this effect (‘Batch’), and entropy of mixing using randomly assigned batch (‘-Ctrl’, no batch effect). The center line denotes the median, while the box contains the 25th to 75th percentiles. The whiskers mark 1.5x interquartile range. (**H**), UMAP of all cells colored by embryo, technical replicates, cell cycle phase, and total UMIs. We made 2 technical replicates for one library (head sample of Emb. 06), thus the 2 replicates are indicated and other cells are in grey in the plot of technical replicate.

**Extended Data Fig. 5.**
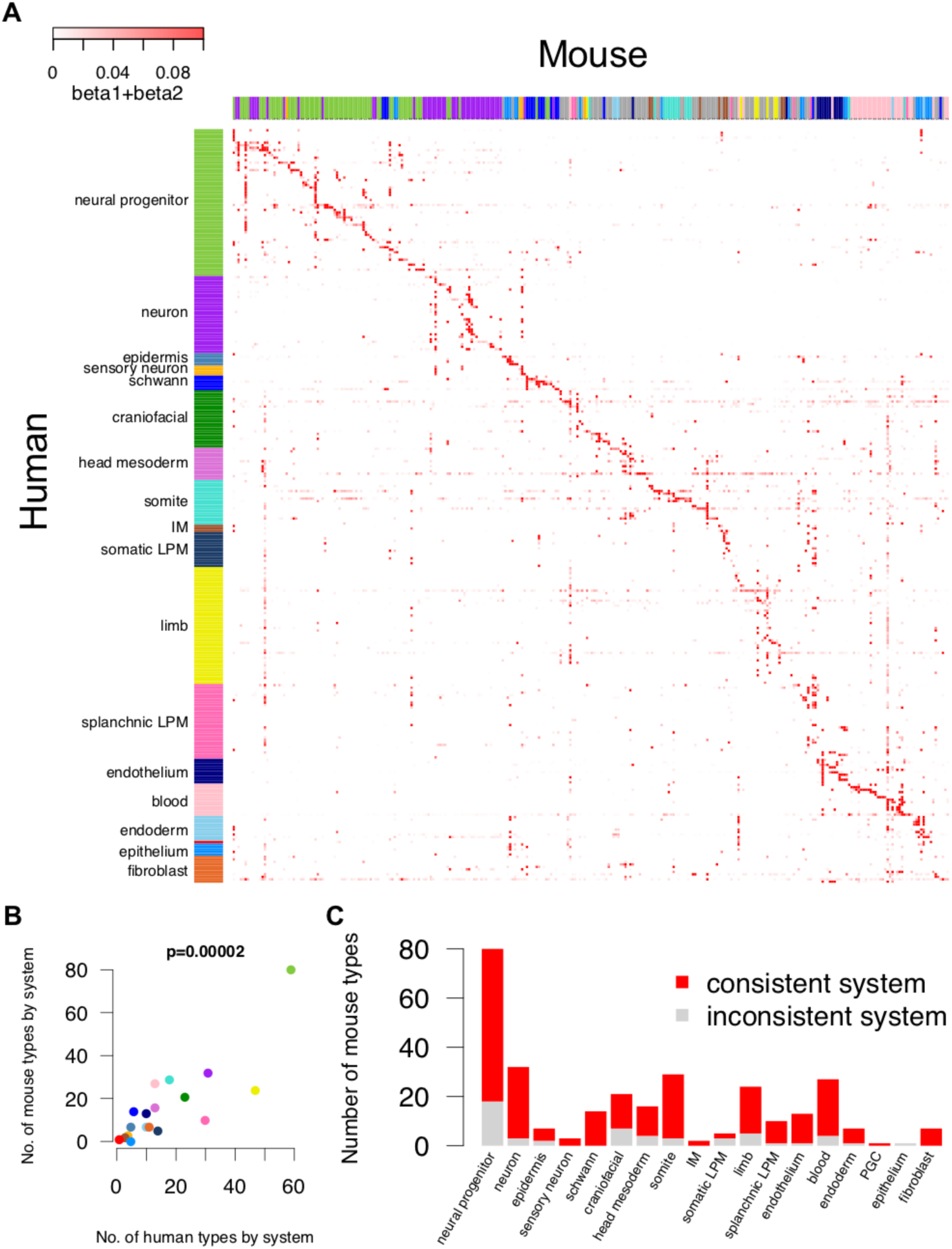
The comparison with mouse snRNA-seq data. (**A**), The specificity score of hit mouse cell types at the corresponding stage^2^ (Methods, Supplementary Table 2). Each row is a human cell type and each column is a mouse cell type. Mouse cell types were ordered by its best matched human type. The color bar on mouse side shows the developmental system of mouse cell type according to annotation form the original publication (Supplementary Table 2). The grey color on columns denotes mesodermal cell types in mouse that can not be further specified to which human mesodermal system. (**B**), The comparison of number of cell types in each human developmental system and number of hit mouse cell types in each human developmental system. The p value was calculated based on Pearson’s product moment correlation coefficient. (**C**), The consistency on the developmental system of hit mouse cell types in each human developmental system.

**Extended Data Fig. 6.**
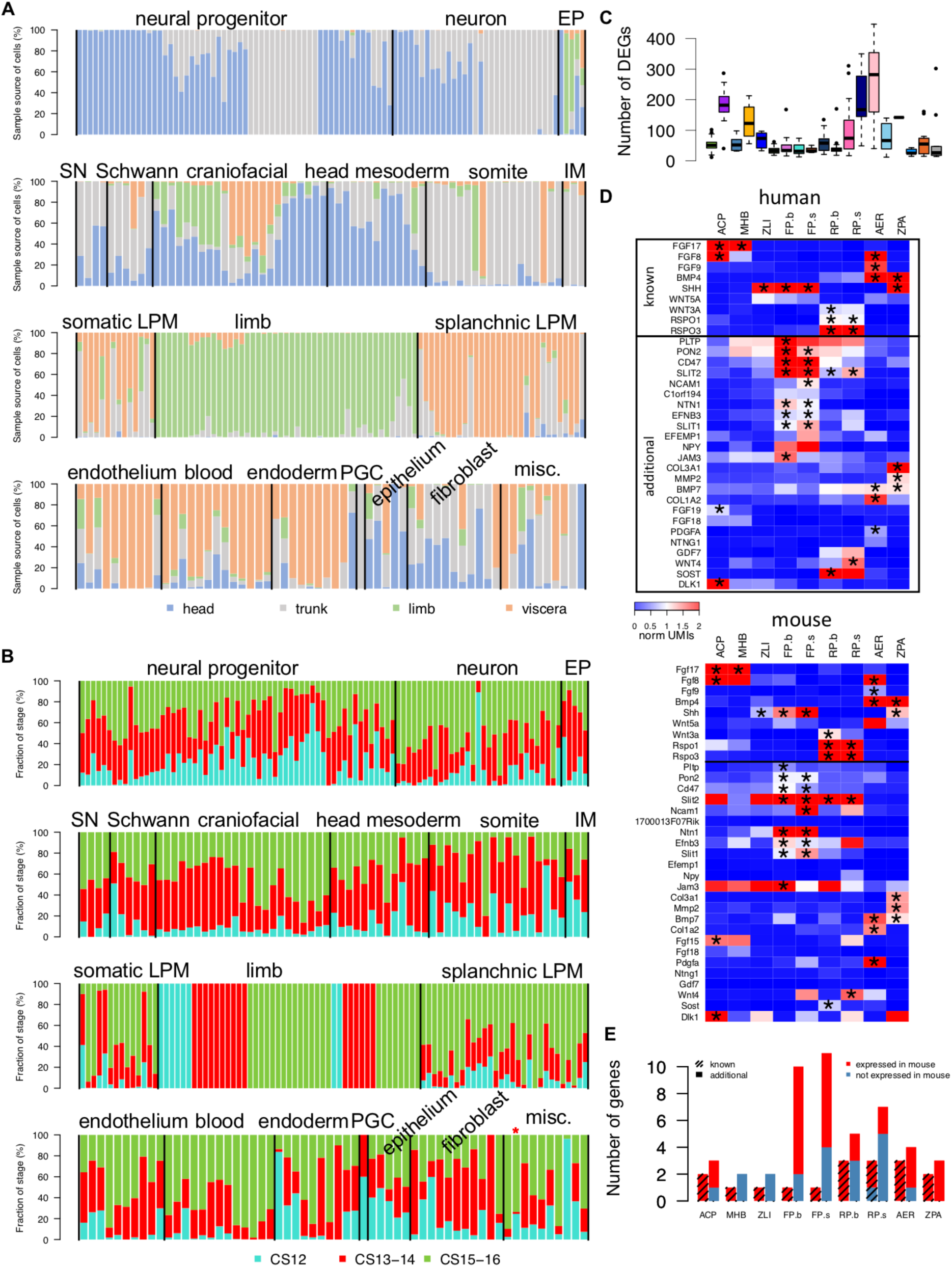
Dissection parts, embryonic stages, and DEGs of cell types. (**A**), The origin of dissection parts of cells in each cell type ordered by system. EP, epidermis; SN, sensory neuron; IM, intermediate mesoderm; LPM, lateral plate mesoderm; PGC, primordial germ cell; misc., miscellaneous. (**B**), Stage distribution of cells in each cell type ordered by system. * denotes cell types missing cells from CS13-14, defined by total number of cells > 50, ratio of cells from CS12 embryo > 0.05, number of cells from CS12 embryo > 5, ratio of cells from CS15-16 embryo > 0.05, number of cells from CS15-16 embryo > 5, ratio of cells from CS13-14 embryo < 0.05, and number of cells from CS13-14 embryo < 5. (**C**), Number of DEGs per cell type for each developmental system (see Fig. 1 for convention). The center line denotes the median, while the box contains the 25th to 75th percentiles. The whiskers mark 1.5x interquartile range. (**D**), The expression of ligands in DEGs of 9 signaling centers in human (top) and their expression in mouse cell types from published data^25–27^ (bottom) (Methods). Asterisks denote ligands that are also expressed in the corresponding cell types in mouse. ANR, anterior neural ridge; MHB, mid-hindbrain boundary; ZLI, zona limitans intrathalamica; FP, floor plate (brain, spinal); RP, roof plate (brain, spinal); AER, apical ectodermal ridge; ZPA, zone of polarizing activity. (**E**), The number of ligands that are expressed or not in mouse by signaling center. For each signaling center, left bar shows known ligands and right bar shows additional ligands.

**Extended Data Fig. 7.**
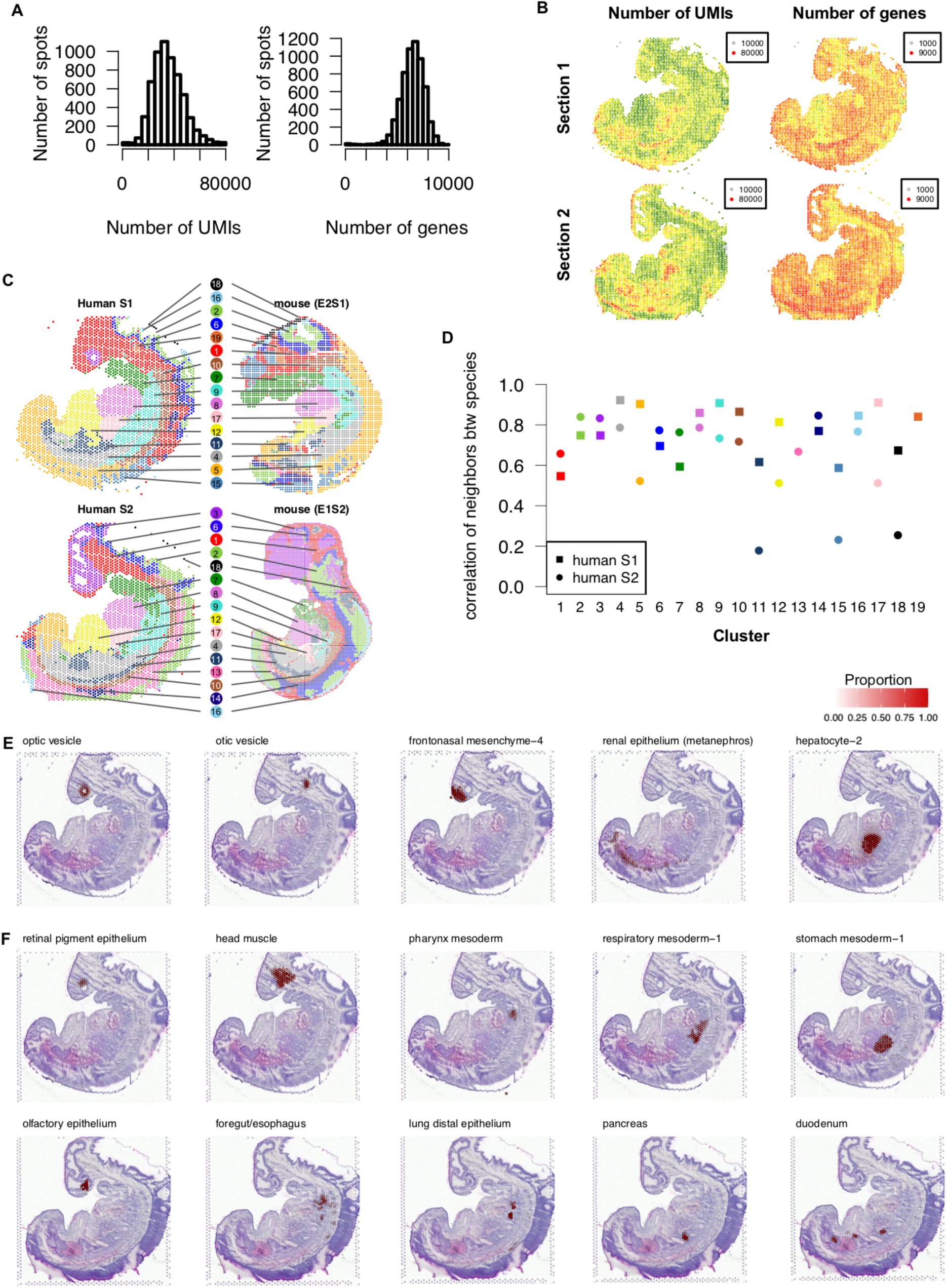
Quality control of spatial transcriptome. (**A**), The distribution of total UMIs and gene numbers per spot. (**B**), The distribution of total UMIs and gene numbers on each section. (**C**), The co-clustering of 2 human sections and 2 published mouse sections^28^ on the spot level (Methods). Cluster identities of spots are indicated by color and number on each tissue section. (**D**), The correlation of spatial neighborhood of each cluster between human and mouse. The spatial neighborhood for a cluster on a section was defined as the proportions of clusters in its neighbors. The neighbors of a cluster were defined as the union of nearest 6 spots on the section of each spot in this cluster, excluding spots from the same cluster. Human section 1 was compared to mouse E2S1 and human section 2 was compared to mouse E1S2. The color and number of each cluster correspond to panel C. (**E**), The proportion of 5 recognizable structures on H&E staining of section 1 in deconvolution. (**F**), The proportion of structures that are difficult to be recognized only by H&E staining on section 1 (first row) and section 2 (second row) in deconvolution. Five structures are showed in each section as examples.

**Extended Data Fig. 8.**
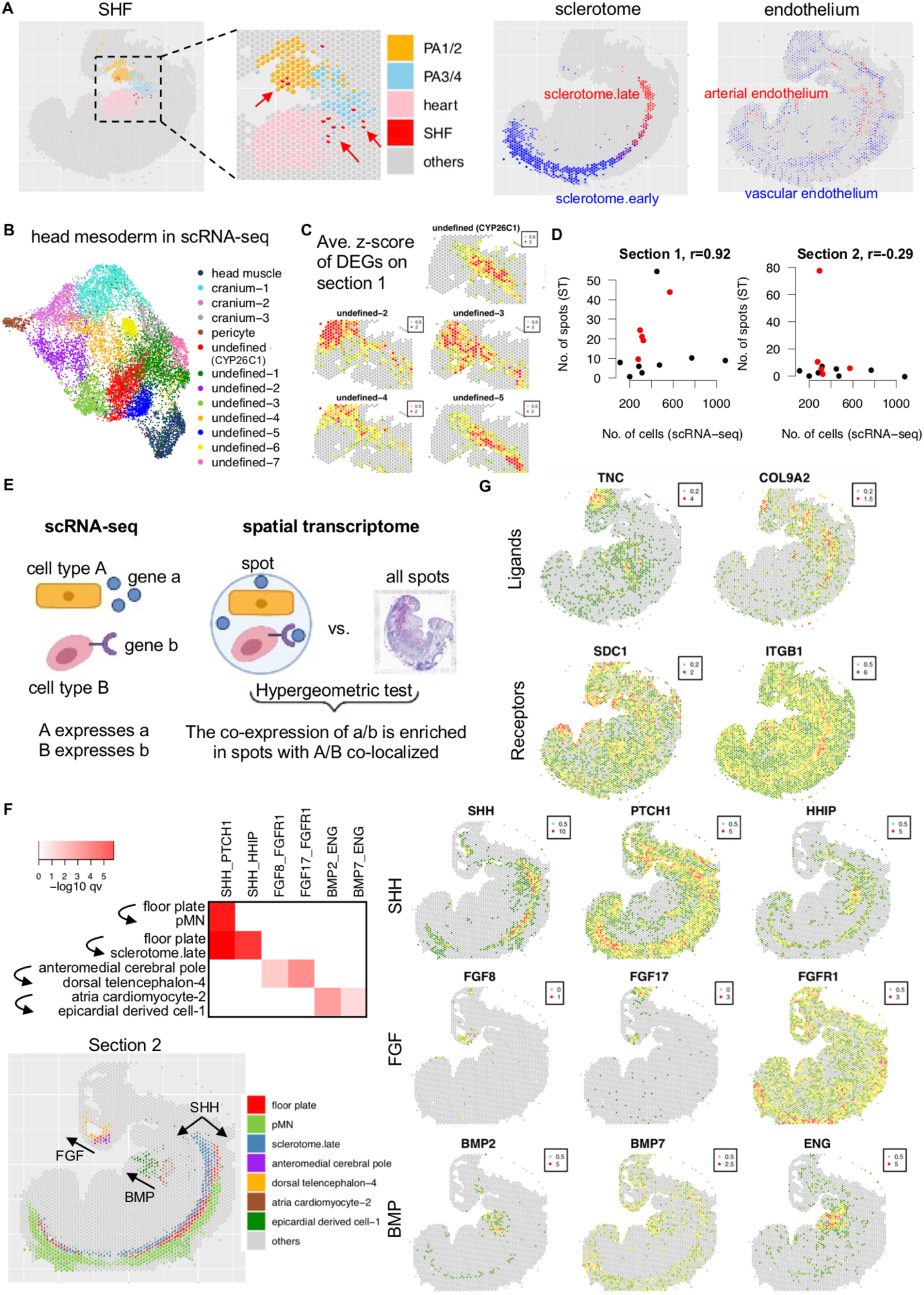
Deconvolution, head mesoderm, and signaling interaction in spatial transcriptome. (**A**), The deconvolution result of selected cell types on section 1. From left to right, second heart field (SHF) along with pharynx arches (PA) and heart, sclerotome, arterial and vascular endothelium. Arrows in the panel of SHF denote the detection of SHF in PA 1/2, PA 3/4, and heart. (**B**), The UMAP of cells in head mesoderm in scRNA-seq colored by cell type. (**C**), The average z-score of DEGs of the 5 undefined cell types in each spot of section 1. The window of ST is the same with that in Fig. 2E. Only top DEGs from scRNA-seq showed in Fig. 2E were used. (**D**), The comparison of detection of cell types between scRNA-seq and ST. Each dot is a cell type in head mesoderm and red dots are the 5 undefined cell types. Pearson’s correlation was calculated on the detection of 5 undefined cell types between scRNA-seq and section 1 (mainly mesodermal tissues), between scRNA-seq and section 2 (mainly ectodermal tissues). To be consistent on developmental stage between scRNA-seq and ST, only cells from CS13-CS14 were counted in scRNA-seq. (**E**), The pipeline of inferring signaling interaction by scRNA-seq and spatial transcriptome (Methods). (**F**), Examples of known signaling interactions identified by integrating scRNA-seq and ST. For all significant interactions, see Supplementary Table 1G. Top left, the -log10 adjusted p value of selected signaling interactions on section 2. Each row is a pair of cell types and each column is a pair of ligand-receptor. Bottom left, the proportion of cell types involved in these interactions in ST. Right panel, the expression of ligands and receptors in these interaction by pathway. (**G**), The expression of genes in ST involved in signaling interaction showed in Fig. 2G. First row is ligand and second row is receptor.

**Extended Data Fig. 9.**
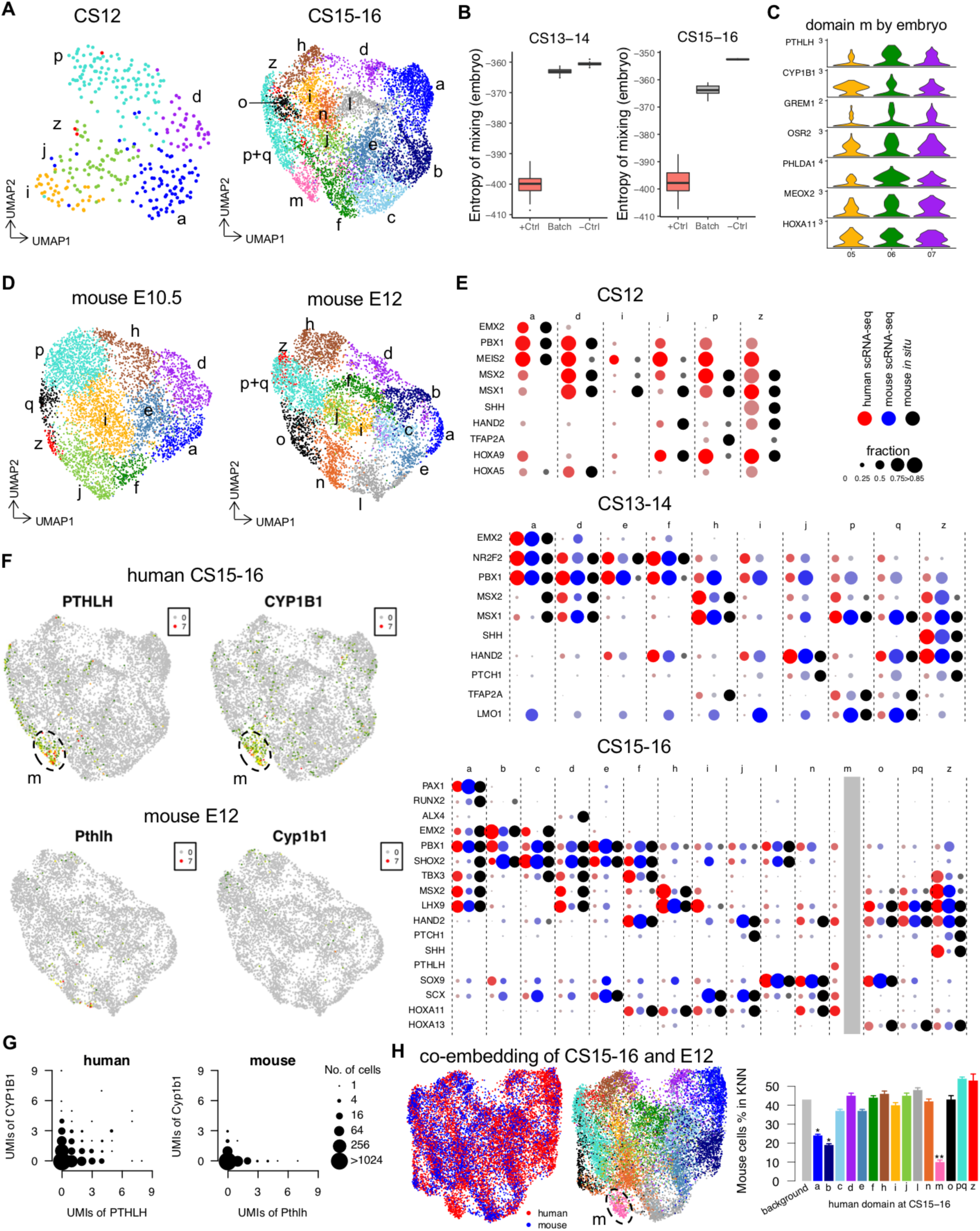
Spatial domains in limb mesenchymal cells. (**A**), UMAP visualization of human forelimb cell types at CS12 and CS15-16. See Fig. 3B for CS13-14. Cells are colored by domains as labeled in Fig. 3. (**B**), Batch effect of embryo estimated by entropy of mixing at CS13-14 and CS15-16. See Extended Data Fig. 4G for conventions. Mann-Whitney U test was applied to compare entropy between batch by embryo and positive control (+Ctrl). (**C**), The expression of top signature genes of domain *m* in domain *m* by embryo. Y-axis denotes normalized UMIs. (**D**), UMAP visualization of mouse forelimb cell types at E10.5 and E12 by reclustering a published dataset^26^. Cells are colored by domains (Supplementary Table 2). (**E**), The expression of marker genes in each domain from human scRNA-seq (red), mouse scRNA-seq (blue), and mouse *in situ* data (green) at three stages. No mouse scRNA-seq data is available at E9.5. Mouse *in situ* data in each domain were manually classified to strong expression (large dot), weak expression (small dot) and no expression (no dot) based on *in situ* images. Domain *m* at E12 was not identified on reclustering of mouse data. (**F**), The expression of specific markers of domain *m* on UMAP in human and mouse. Dash circles denote domain *m* in UMAP of human data. (**G**), The UMIs of *PTHLH* and *CYP1B1* in each cell at CS15-16 of human and E12 of mouse. The size of dot denotes number of cells with corresponding values. (**H**), Left panel, Umap co-embedding of CS15-16 of human and E12 of human colored by species and domains (see panel A for colors of domains). Dash circles denote domain *m* in co-embedding. Right panel, the average percentage of mouse cells in k-nearest neighbors (KNN) of human cells in each domain in co-embedding space (Methods). The first bar (background) denotes the percentage of mouse cells in total cells of human and mouse. Error bars denote standard error. Mann-Whitney U test was applied to compare each domain and background. Significant levels of adjusted p values are denoted with asterisks (domain *a*, 10^-7^; domain *b*, 10^-7^; domain *m*, 10^-9^).

**Extended Data Fig. 10.**
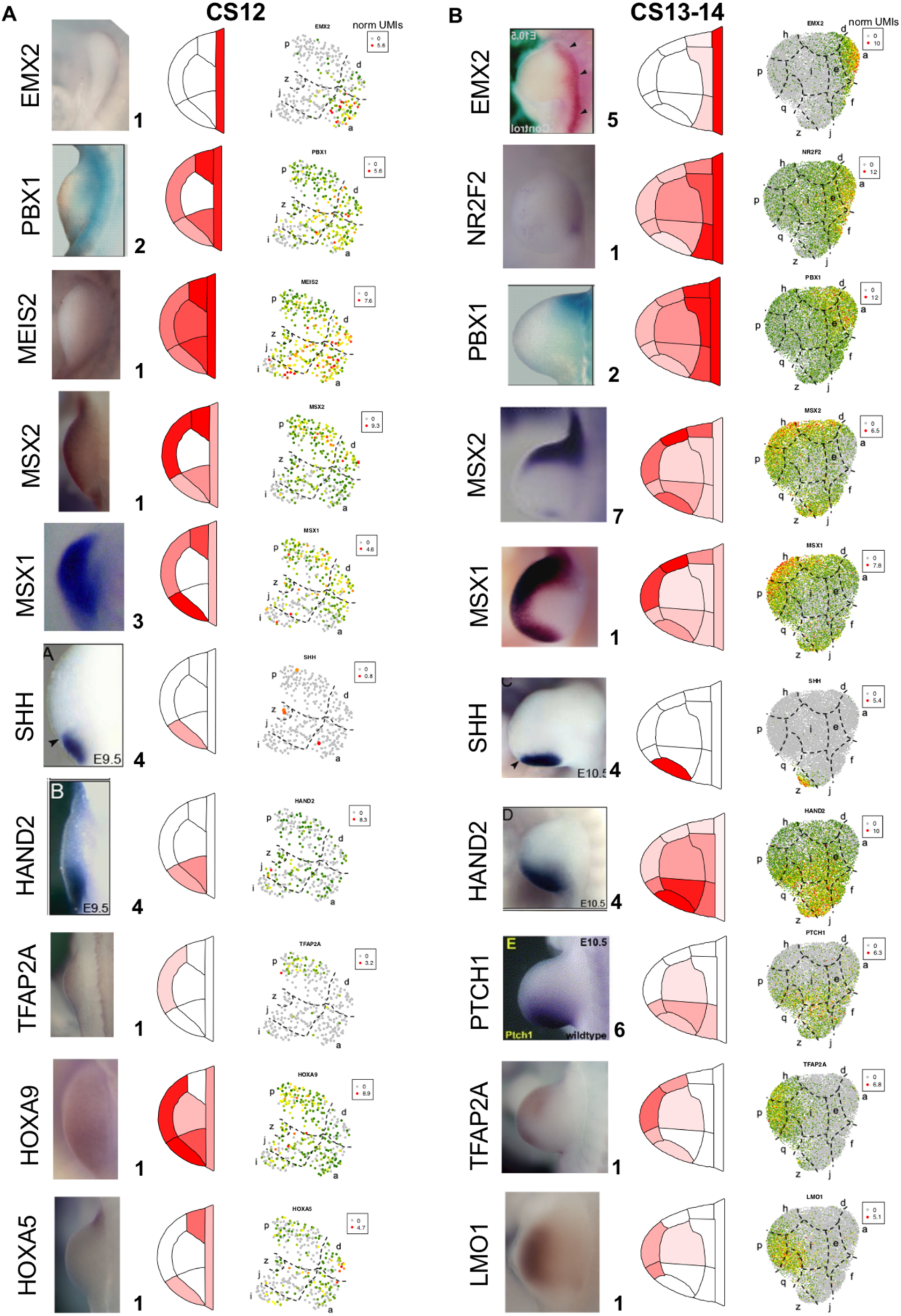
*In situ* evidence of spatial domains in limb mesenchymal cells. The comparison of marker genes in *in situ* and scRNA-seq at CS12 (**A**) and CS13-14 (**B**). *In situ* results were collected from mouse studies at the corresponding stage (E9.5 for CS12 and E10.5 for CS13-14). Numbers following each *in situ* picture denote references (see Extended Data Fig. 11). The gene expression in spatial domains in scRNA-seq were showed in schematic diagram colored by mean of normalized UMIs in the domain (2nd column). The gene expression in each cell in scRNA-seq were showed on UMAP (3rd column).

**Extended Data Fig. 11.**
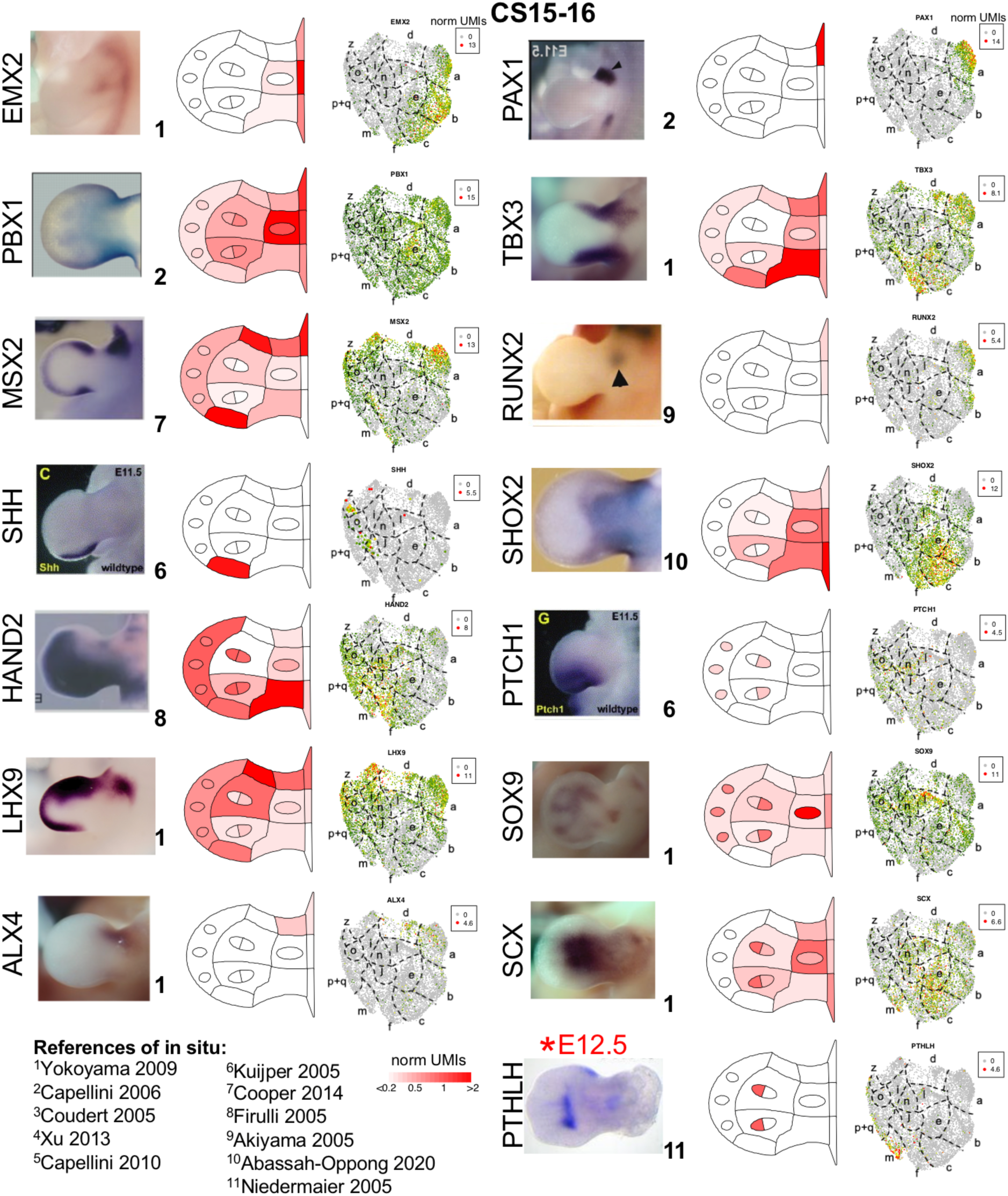
*In situ* evidence of spatial domains in limb mesenchymal cells. The comparison of marker genes in *in situ* at E11.5 (except *Pthlh*) and scRNA-seq at CS15-16. See Extended Data Fig. 10 for convention. The *in situ* data of *Pthlh* is from E12.5 of mouse because no *in situ* data is available at E11.5. For the detail of references, please see reference section.

**Extended Data Fig. 12.**
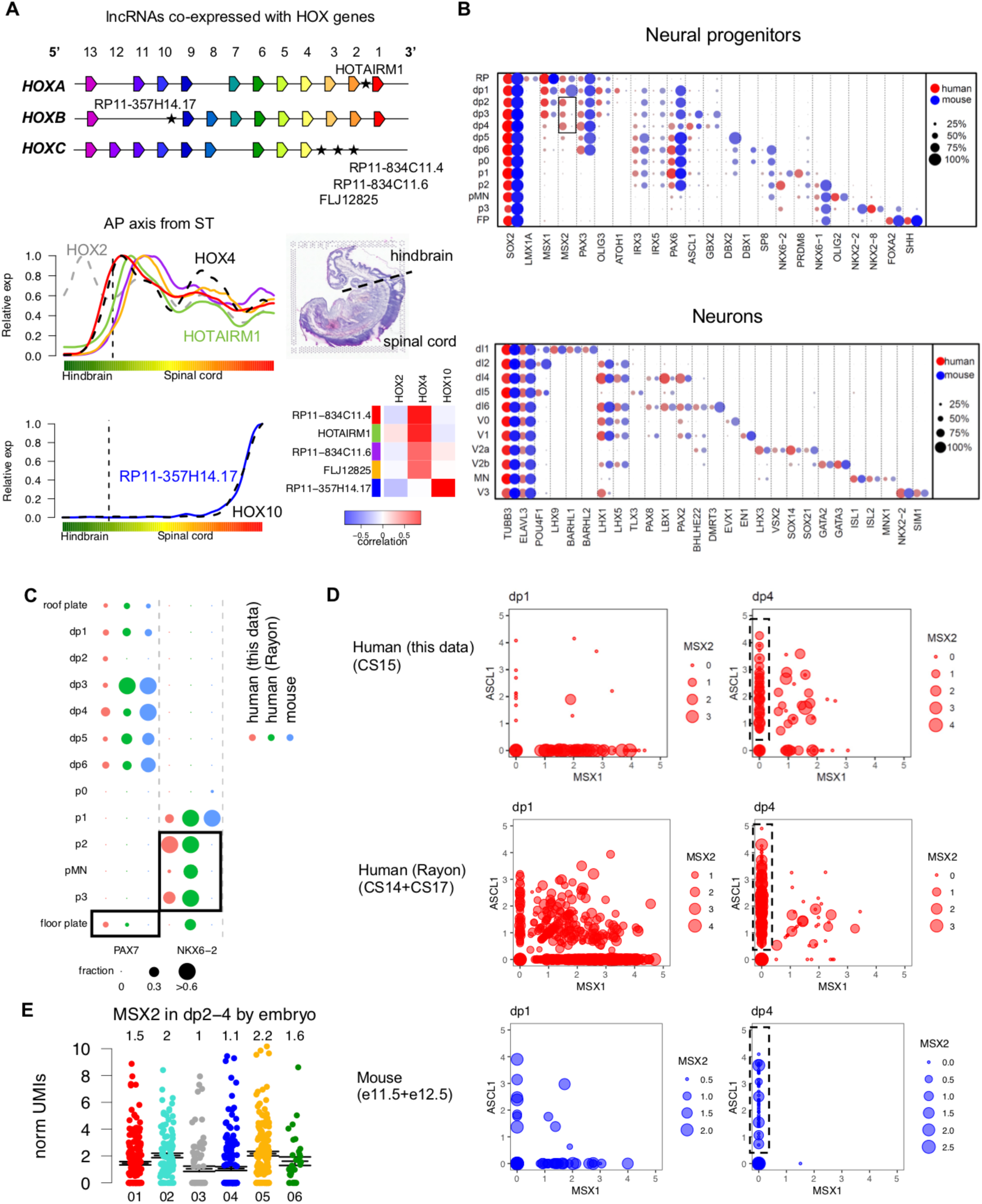
Neural tube patterning. (**A**), Upper panel, schematic diagram of the genomic location of the 5 lncRNAs identified as AP related genes in neural tube. Lower-left panel, the expression pattern of 5 lncRNAs (solid lines, see left bar of the heatmap for color legend) and nearby HOX genes (dash lines) along the AP axis on section 2 of ST. Lower-right panel, the division between hindbrain and spinal cord on H&E staining of section 2, and the correlation on gene expression between lncRNAs and HOX genes along AP axis of section 2 (Methods). (**B**), The comparison of expression of canonical markers for cell types in neural progenitors (upper panel) and neurons (lower panel) between human and mouse. Box denotes the difference of *MSX2* between human and mouse in neural progenitors. (**C**), The comparison of expression of *PAX7* and *NKX6-2* between human and mouse in neural progenitors. Boxes denote human specific expression that are consistent between two datasets of human. (**D**), The expression of markers (dp1 marker *MSX1* and dp4 marker *ASCL1*, log2 scaled) that distinguish dp1 and dp4 cells in individual cells in human (our dataset and Rayon’s dataset) and mouse datasets. Dot size shows *MSX2* expression. Dash boxes denote most confident dp4 cells (*ASCL1*>0 and *MSX1*=0), which are showed in Fig. 4F. (**E**), The expression of *MSX2* in dp2-4 in each human embryo in our data. Each dot denotes a cell. Black lines denote mean and standard error. The number on top is mean of normalized UMIs in each embryo. Note the trunk sample of Emb.07 did not pass quality control so that no cell is from Emb.07.

**Extended Data Fig. 13.**
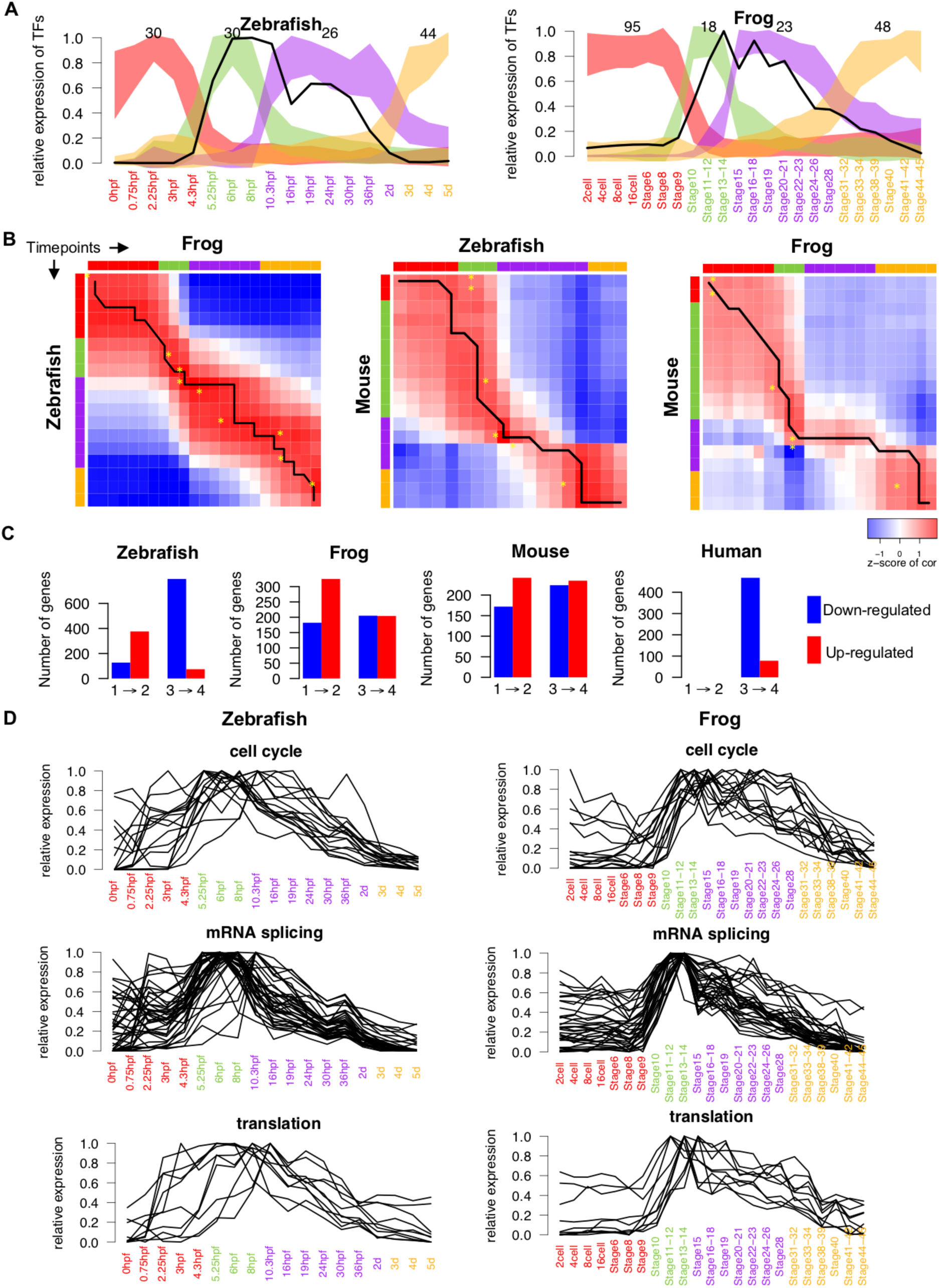
LIN28A in vertebrate embryogenesis. (**A**), Expression pattern of 4 groups of TFs in zebrafish and frog. Each group of TFs were identified as TFs that are highly expressed in the corresponding stage. Numbers above curves indicate number of TFs in each group. The width of each curve represents 2 standard deviations within each group of TFs. The black lines show the expression of *LIN28A* in each species. (**B**), Pairwise correlation of timepoints between species by homologous TFs. Correlation is scaled by row (Methods). Black line denotes the alignment of timepoints between species by dynamic time warping. The yellow asterisks denote the match of timepoints from a previous study^105^. (**C**), Numbers of systemically up- (red) and down-regulated (blue) genes from stage 1 to stage 2 (labeled as 1➜2) and from stage 3 to stage 4 (labeled as 3➜4). No transcriptome data is available for stages 1 and 2 in human. (**D**), Expression dynamics of genes in cell cycle, mRNA splicing and translation pathways that are positively correlated to *LIN28A* in zebrafish and frog.

**Extended Data Fig. 14.**
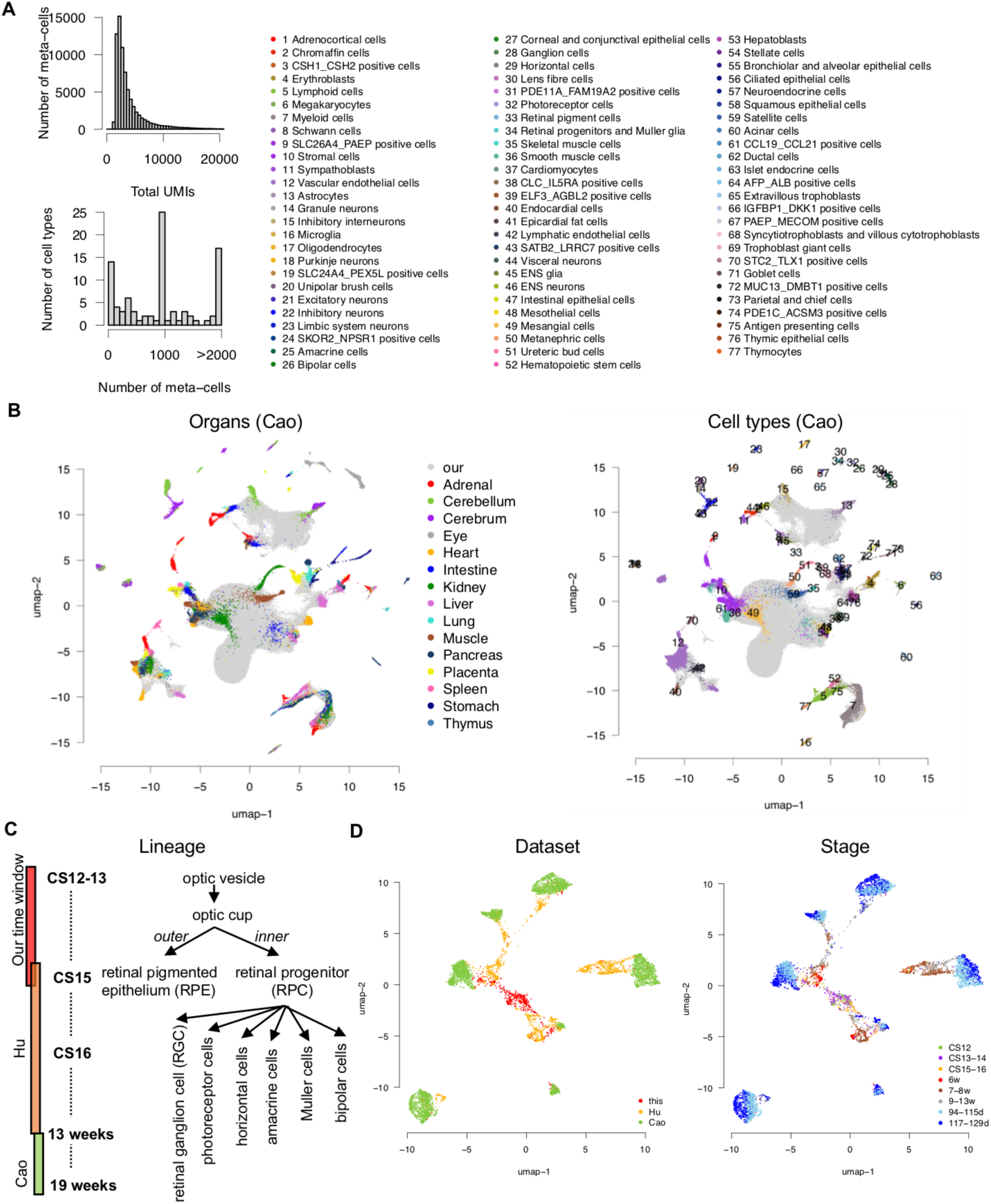
Data integration in human embryos. (**A**), Left panel, the distribution of total UMIs in meta-cells in Cao’s data and the distribution of numbers of meta-cells in each cell type in Cao’s data. See Methods for the definition of meta-cell. Right panel, legend of cell types in Cao’s data for panel B. (**B**), The joint UMAP of our dataset (grey) and Cao’s dataset colored by organ source and by cell type in Cao’s dataset. (**C**), Lineage development of human eye and time windows of three studies in integration. (**D**), Joint UMAP of cell types of eye in three studies colored by dataset and stage.

## Supplementary tables and information

**Supplementary Table 1. (separate file)**

Cell types and DEGs.

**Supplementary Table 2. (separate file)**

The comparison between human and mouse sn/scRNA-seq.

**Supplementary Table 3. (separate file)**

LIN28A in vertebrate embryogenesis.

**Supplementary Table 4. (separate file)**

Systemically changing genes in vertebrate embryogenesis.

**Supplementary Table 5. (separate file)**

Enriched pathways in systemically changing genes.

**Supplementary Table 6. (separate file)**

Data integration in human embryos.

**Supplementary Note 1. (separate file)**

Annotation vignette.

**Supplementary Note 2. (separate file)**

The proportion of each cell type in the deconvolution of section 1 of spatial transcriptome. Blank plot denotes no detection in deconvolution for a given cell type.

**Supplementary Note 3. (separate file)**

The proportion of each cell type in the deconvolution of section 2 of spatial transcriptome. Blank plot denotes no detection in deconvolution for a given cell type.

## References

1 Schoenwolf, G. C. & Larsen, W. J. (William J. Larsen’s human embryology. (Churchill Livingstone/Elsevier, 2009).

2 Cao, J. et al. The single-cell transcriptional landscape of mammalian organogenesis. Nature 1 (2019) doi:10.1038/s41586-019-0969-x.

3 Farrell, J. A. et al. Single-cell reconstruction of developmental trajectories during zebrafish embryogenesis. Science (80-.). 360, eaar3131 (2018).

4 Wagner, D. E. et al. Single-cell mapping of gene expression landscapes and lineage in the zebrafish embryo. Science (80-.). 360, 981–987 (2018).

5 Pijuan-Sala, B. et al. A single-cell molecular map of mouse gastrulation and early organogenesis. Nature 1 (2019) doi:10.1038/s41586-019-0933-9.

6 Han, X. et al. Construction of a human cell landscape at single-cell level. Nature 581, 303–309 (2020).

7 Cao, J. et al. A human cell atlas of fetal gene expression. Science (80-.). 370, eaba7721 (2020).

8 Polioudakis, D. et al. A Single-Cell Transcriptomic Atlas of Human Neocortical Development during Mid-gestation. Neuron 0, (2019).

9 Hu, Y. et al. Dissecting the transcriptome landscape of the human fetal neural retina and retinal pigment epithelium by single-cell RNA-seq analysis. PLOS Biol. 17, e3000365 (2019).

10 Li, L. et al. Single-Cell RNA-Seq Analysis Maps Development of Human Germline Cells and Gonadal Niche Interactions. Cell Stem Cell 20, 891–892 (2017).

11 Gao, S. et al. Tracing the temporal-spatial transcriptome landscapes of the human fetal digestive tract using single-cell RNA-sequencing. Nat. Cell Biol. 20, 721–734 (2018).

12 Popescu, D.-M. et al. Decoding human fetal liver haematopoiesis. Nature 1–7 (2019) doi:10.1038/s41586-019-1652-y.

13 Asp, M. et al. A Spatiotemporal Organ-Wide Gene Expression and Cell Atlas of the Developing Human Heart. Cell 179, 1647–1660.e19 (2019).

14 Nowakowski, T. J. et al. Spatiotemporal gene expression trajectories reveal developmental hierarchies of the human cortex. Science (80-.). 358, 1318–1323 (2017).

15 Zhong, S. et al. A single-cell RNA-seq survey of the developmental landscape of the human prefrontal cortex. Nature 1. (2018) doi:10.1038/nature25980.

16 Young, M. D. et al. Single-cell transcriptomes from human kidneys reveal the cellular identity of renal tumors. Science (80-.). 361, 594–599 (2018).

17 Li, M. et al. Integrative functional genomic analysis of human brain development and neuropsychiatric risks. Science (80-.). 362, eaat7615 (2018).

18 Gao, R. et al. Delineating copy number and clonal substructure in human tumors from single-cell transcriptomes. Nat. Biotechnol. 2021 395 39, 599–608 (2021).

19 Dyer, L. A. & Kirby, M. L. The role of secondary heart field in cardiac development. Dev Biol 336, 137–144 (2009).

20 Alquicira-Hernandez, J., Sathe, A., Ji, H. P., Nguyen, Q. & Powell, J. E. ScPred: Accurate supervised method for cell-type classification from single-cell RNA-seq data. Genome Biol. 20, 1–17 (2019).

21 Cabello-Aguilar, S. et al. SingleCellSignalR: inference of intercellular networks from single-cell transcriptomics. Nucleic Acids Res. 48, e55 (2020).

22 Capdevila, J. & Izpisua Belmonte, J. C. Patterning mechanisms controlling vertebrate limb development. Annu Rev Cell Dev Biol 17, 87–132 (2001).

23 Echevarria, D., Vieira, C., Gimeno, L. & Martinez, S. Neuroepithelial secondary organizers and cell fate specification in the developing brain. Brain Res Brain Res Rev 43, 179–191 (2003).

24 Ramilowski, J. A. et al. A draft network of ligand–receptor-mediated multicellular signalling in human. Nat. Commun. 6, 7866 (2015).

25 Delile, J. et al. Single cell transcriptomics reveals spatial and temporal dynamics of gene expression in the developing mouse spinal cord. Development 146, dev.173807 (2019).

26 He, P. et al. The changing mouse embryo transcriptome at whole tissue and single-cell resolution. Nature 583, 760– 767 (2020).

27 La Manno, G. et al. Molecular architecture of the developing mouse brain. Nature 1–5 (2021) doi:10.1038/s41586-021-03775-x.

28 Chen, A. et al. Spatiotemporal transcriptomic atlas of mouse organogenesis using DNA nanoball-patterned arrays. Cell 185, 1777–1792.e21 (2022).

29 Cable, D. M. et al. Robust decomposition of cell type mixtures in spatial transcriptomics. Nat. Biotechnol. 2021 1–10 (2021) doi:10.1038/s41587-021-00830-w.

30 Trainor, P. A., Tan, S. S. & Tam, P. P. L. Cranial paraxial mesoderm: Regionalisation of cell fate and impact on craniofacial development in mouse embryos. Development 120, 2397–2408 (1994).

31 Bumcrot, D. A. & McMahon, A. P. Somite Differentiation: Sonic signals somites. Curr. Biol. 5, 612–614 (1995).

32 Chen, B., Kwan, K. Y., Rubenstein, J. L. R. & Rakic, P. Patterning and Cell Type Specification in the Developing CNS and PNS: comprehensive developmental neuroscience. Amsterdam: Academic Press (2020). doi:10.1016/c2017-0-00860-1.

33 van Wijk, B., Moorman, A. F. M. & van den Hoff, M. J. B. Role of bone morphogenetic proteins in cardiac differentiation. Cardiovasc. Res. 74, 244–255 (2007).

34 Kim, S. H., Turnbull, J. & Guimond, S. Extracellular matrix and cell signalling: the dynamic cooperation of integrin, proteoglycan and growth factor receptor. J. Endocrinol. 209, 139–151 (2011).

35 Chiovaro, F., Chiquet-Ehrismann, R. & Chiquet, M. Transcriptional regulation of tenascin genes. Cell Adh. Migr. 9, 34 (2015).

36 Benazet, J. D. & Zeller, R. Vertebrate limb development: moving from classical morphogen gradients to an integrated 4-dimensional patterning system. Cold Spring Harb Perspect Biol 1, a001339 (2009).

37 Yokoyama, S. et al. A Systems Approach Reveals that the Myogenesis Genome Network Is Regulated by the Transcriptional Repressor RP58. Dev. Cell 17, 836–848 (2009).

38 Capellini, T. D. et al. Pbx1/Pbx2 requirement for distal limb patterning is mediated by the hierarchical control of Hox gene spatial distribution and Shh expression. Development 133, 2263–2273 (2006).

39 Coudert, A. E. et al. Expression and regulation of the Msx1 natural antisense transcript during development. Nucleic Acids Res. 33, 5208–18 (2005).

40 B,X. et al. Hox5 interacts with Plzf to restrict Shh expression in the developing forelimb. Proc. Natl. Acad. Sci. U. S. A. 110, 19438–19443 (2013).

41 Capellini, T. D. et al. Scapula development is governed by genetic interactions of Pbx1 with its family members and with Emx2 via their cooperative control of Alx1. Development 137, 2559–2569 (2010).

42 Kuijper, S. et al. Function and regulation of Alx4 in limb development: complex genetic interactions with Gli3 and Shh. Dev Biol 285, 533–544 (2005).

43 Cooper, K. L. et al. Patterning and post-patterning modes of evolutionary digit loss in mammals. Nature 511, 41–45 (2014).

44 Firulli, B. A. et al. Altered Twist1 and Hand2 dimerization is associated with Saethre-Chotzen syndrome and limb abnormalities. Nat. Genet. 37, 373–381 (2005).

45 Akiyama, H. et al. Osteo-chondroprogenitor cells are derived from Sox9 expressing precursors. Proc. Natl. Acad. Sci. U. S. A. 102, 14665–70 (2005).

46 Abassah-Oppong, S. et al. A gene desert required for regulatory control of pleiotropic Shox2 expression and embryonic survival. bioRxiv 2020.11.22.393173 (2020) doi:10.1101/2020.11.22.393173.

47 Bergen, V., Lange, M., Peidli, S., Wolf, F. A. & Theis, F. J. Generalizing RNA velocity to transient cell states through dynamical modeling. Nat. Biotechnol. (2020) doi:10.1038/s41587-020-0591-3.

48 Vargesson, N. et al. Cell fate in the chick limb bud and relationship to gene expression. Development 124, (1997).

49 Zakany, J. & Duboule, D. The role of Hox genes during vertebrate limb development. Curr. Opin. Genet. Dev. 17, 359–366 (2007).

50 Zhang, B. et al. A human embryonic limb cell atlas resolved in space and time. bioRxiv 2022.04.27.489800 (2022) doi:10.1101/2022.04.27.489800.

51 Lefebvre, V. & Smits, P. Transcriptional control of chondrocyte fate and differentiation. Birth Defects Res. Part C Embryo Today Rev. 75, 200–212 (2005).

52 Kobayashi, T. et al. Indian hedgehog stimulates periarticular chondrocyte differentiation to regulate growth plate length independently of PTHrP. J. Clin. Invest. 115, 1734–42 (2005).

53 Niedermaier, M. et al. An inversion involving the mouse Shh locus results in brachydactyly through dysregulation of Shh expression. J. Clin. Invest. 115, 900–909 (2005).

54 Duboule, D. The (unusual) heuristic value of Hox gene clusters; a matter of time? Dev. Biol. 484, 75–87 (2022).

55 Philippidou, P. & Dasen, J. S. Hox genes: choreographers in neural development, architects of circuit organization. Neuron 80, 12–34 (2013).

56 Le Dreau, G. & Marti, E. Dorsal-ventral patterning of the neural tube: a tale of three signals. Dev Neurobiol 72, 1471– 1481 (2012).

57 Uehara, M. et al. CYP26A1 and CYP26C1 cooperatively regulate anterior-posterior patterning of the developing brain and the production of migratory cranial neural crest cells in the mouse. Dev Biol 302, 399–411 (2007).

58 Sarropoulos, I., Marin, R., Cardoso-Moreira, M. & Kaessmann, H. Developmental dynamics of lncRNAs across mammalian organs and species. Nature 571, 510–514 (2019).

59 De Kumar, B. & Krumlauf, R. HOXs and lincRNAs: Two sides of the same coin. Sci Adv 2, e1501402 (2016).

60 Zhang, X. et al. A myelopoiesis-associated regulatory intergenic noncoding RNA transcript within the human HOXA cluster. Blood 113, 2526–2534 (2009).

61 Wilson, L. & Maden, M. The mechanisms of dorsoventral patterning in the vertebrate neural tube. Dev Biol 282, 1– 13 (2005).

62 Delile, J. et al. Single cell transcriptomics reveals spatial and temporal dynamics of gene expression in the developing mouse spinal cord. Development 146, (2019).

63 Rayon, T., Maizels, R., Barrington, C. & Briscoe, J. Single-cell transcriptome profiling of the human developing spinal cord reveals a conserved genetic programme with human-specific features. Development 148, (2021).

64 Moss, E. G. Heterochronic Genes and the Nature of Developmental Time. Curr. Biol. 17, R425–R434 (2007).

65 Ambros, V. & Horvitz, H. Heterochronic mutants of the nematode Caenorhabditis elegans. Science (80-.). 226, 409–416 (1984).

66 Boroviak, T. et al. Lineage-Specific Profiling Delineates the Emergence and Progression of Naive Pluripotency in Mammalian Embryogenesis. Dev. Cell 35, 366–382 (2015).

67 Tan, M. H. et al. RNA sequencing reveals a diverse and dynamic repertoire of the Xenopus tropicalis transcriptome over development. Genome Res. 23, 201–16 (2013).

68 Briggs, J. A. et al. The dynamics of gene expression in vertebrate embryogenesis at single-cell resolution. Science (80-.). 360, eaar5780 (2018).

69 White, R. J. et al. A high-resolution mRNA expression time course of embryonic development in zebrafish. Elife 6, (2017).

70 Yang, D.-H. & Moss, E. G. Temporally regulated expression of Lin-28 in diverse tissues of the developing mouse. Gene Expr. Patterns 3, 719–726 (2003).

71 Faas, L. et al. Lin28 proteins are required for germ layer specification in Xenopus. Development 140, 976–86 (2013).

72 Nieuwkoop, P. D. & Faber, J. Normal Table of Xenopus laevis (Daudin). (Oxford, UK: Taylor and Francis, 1994).

73 Carroll, E. J. & Hedrick, J. L. Hatching in the toad Xenopus laevis: Morphological events and evidence for a hatching enzyme. Dev. Biol. 38, 1–13 (1974).

74 Irie, N. & Kuratani, S. Comparative transcriptome analysis reveals vertebrate phylotypic period during organogenesis. Nat. Commun. 2, 248 (2011).

75 Cardoso-Moreira, M. et al. Gene expression across mammalian organ development. Nature 1 (2019) doi:10.1038/s41586-019-1338-5.

76 Shyh-Chang, N. & Daley, G. Q. Lin28: Primal Regulator of Growth and Metabolism in Stem Cells. Cell Stem Cell 12, 395–406 (2013).

77 Wilbert, M. L. et al. LIN28 Binds Messenger RNAs at GGAGA Motifs and Regulates Splicing Factor Abundance. Mol. Cell 48, 195–206 (2012).

78 Agarwal, V., Bell, G. W., Nam, J.-W. & Bartel, D. P. Predicting effective microRNA target sites in mammalian mRNAs. Elife 4, (2015).

79 Shinoda, G. et al. Fetal deficiency of lin28 programs life-long aberrations in growth and glucose metabolism. Stem Cells 31, 1563–73 (2013).

80 Robinton, D. A. et al. The Lin28/let-7 Pathway Regulates the Mammalian Caudal Body Axis Elongation Program. Dev. Cell 48, 396–405.e3 (2019).

81 Komarovsky Gulman, N., Armon, L., Shalit, T. & Urbach, A. Heterochronic regulation of lung development via the Lin28-Let-7 pathway. FASEB J. 33, 12008–12018 (2019).

82 Street, K. et al. Slingshot: cell lineage and pseudotime inference for single-cell transcriptomics. BMC Genomics 19, 477 (2018).

83 Marquardt, T. & Gruss, P. Generating neuronal diversity in the retina: one for nearly all. Trends Neurosci. 25, 32–38 (2002).

84 Chu Y. & Liu, T. On the shortest arborescence of a directed graph. Sci. Sin. 14, 1396–1400 (1965).

85 Fuhrmann, S. Eye morphogenesis and patterning of the optic vesicle. Curr. Top. Dev. Biol. 93, 61–84 (2010).

86 Theos, A. C., Truschel, S. T., Raposo, G. & Marks, M. S. The Silver locus product Pmel17/gp100/Silv/ME20: controversial in name and in function. Pigment Cell Res 18, 322–336 (2005).

87 Ma, X. et al. The transcription factor MITF in RPE function and dysfunction. Prog Retin Eye Res 73, 100766 (2019).

88 Nowotschin, S. et al. The emergent landscape of the mouse gut endoderm at single-cell resolution. Nature (2019) doi:10.1038/s41586-019-1127-1.

89 Regev, A. et al. The Human Cell Atlas. Elife 6, (2017).

90 Regev, A., et al. The Human Cell Atlas White Paper. (2018).

91 Tyser, R. C. V. et al. Single-cell transcriptomic characterization of a gastrulating human embryo. Nature 1–5 (2021) doi:10.1038/s41586-021-04158-y.

92 Moris, N. et al. An in vitro model of early anteroposterior organization during human development. Nature 1–6 (2020) doi:10.1038/s41586-020-2383-9.

93 O’Rahilly, R. & Müller, F. Developmental stages in human embryos: revised and new measurements. Cells. Tissues. Organs 192, 73–84 (2010).

94 Kaufman & Matthew, H. Atlas Of Mouse Development. (Springer Berlin Heidelberg, 2007).

95 SL, W., R, L. & AM, K. Scrublet: Computational Identification of Cell Doublets in Single-Cell Transcriptomic Data. Cell Syst. 8, 281–291.e9 (2019).

96 Zhang, T. et al. A single-cell analysis of the molecular lineage of chordate embryogenesis. Sci. Adv. 6, eabc4773 (2020).

97 Butler, A., Hoffman, P., Smibert, P., Papalexi, E. & Satija, R. Integrating single-cell transcriptomic data across different conditions, technologies, and species. Nat. Biotechnol. 36, 411–420 (2018).

98 Santos, A., Wernersson, R. & Jensen, L. J. Cyclebase 3.0: A multi-organism database on cell-cycle regulation and phenotypes. Nucleic Acids Res. 43, D1140–D1144 (2015).

99 Giotti, B. et al. Assembly of a parts list of the human mitotic cell cycle machinery. J. Mol. Cell Biol. 11, 703–718 (2019).

100 Zeisel, A. et al. Molecular Architecture of the Mouse Nervous System. Cell 174, 999–1014.e22 (2018).

101 Haghverdi, L., Lun, A. T. L., Morgan, M. D. & Marioni, J. C. Batch effects in single-cell RNA-sequencing data are corrected by matching mutual nearest neighbors. Nat. Biotechnol. 36, 421–427 (2018).

102 Qiu, X. et al. Reversed graph embedding resolves complex single-cell trajectories. Nat. Methods 14, 979–982 (2017).

103 Subramanian, A. et al. Gene set enrichment analysis: a knowledge-based approach for interpreting genome-wide expression profiles. Proc. Natl. Acad. Sci. U. S. A. 102, 15545–50 (2005).

104 Packer, J. S. et al. A lineage-resolved molecular atlas of C. elegans embryogenesis at single cell resolution. doi:10.1101/565549.

105 Irie, N. & Kuratani, S. Comparative transcriptome analysis reveals vertebrate phylotypic period during organogenesis. Nat. Commun. 2, 248 (2011).

