## Supplementary Note 1 for "A single-cell transcriptome atlas of human early embryogenesis"

### Annotation Vignette

#### 1 Overview

This note demonstrates how we annotated cell types. In this note, the developmental system ‘blood’ is used as a general example to show the annotation process and ‘limb’ as a special case. For all other systems, the detail can be found in “annotation_all” on https://heoa.shinyapps.io/code/.

#### 2 The clustering of cell types

Before annotation, this section gives a brief summary of how the clustering was performed in each system using ‘blood’ as example. For other systems, please visit our online depository https://heoa.shinyapps.io/code/.

##### 2.1 Identification of highly variable genes (HVGs)

We begin with the normalized matrix of gene by cell, which contains cells identified as blood system in the global clustering. HVGs in each developmental system were identified by Poisson distribution in cells from this system (Zhang et al. 2020, Pubmed ID: 33148647). To avoid genes that are systemically different between samples were identified as HVGs, HVGs were detected by sample and then merged (Figure 1). Then hemoglobin genes, mitochondrial genes, cell cycle genes, sex-specific genes, and batch-effect genes collected from literatures were excluded from HVGs (see Methods).

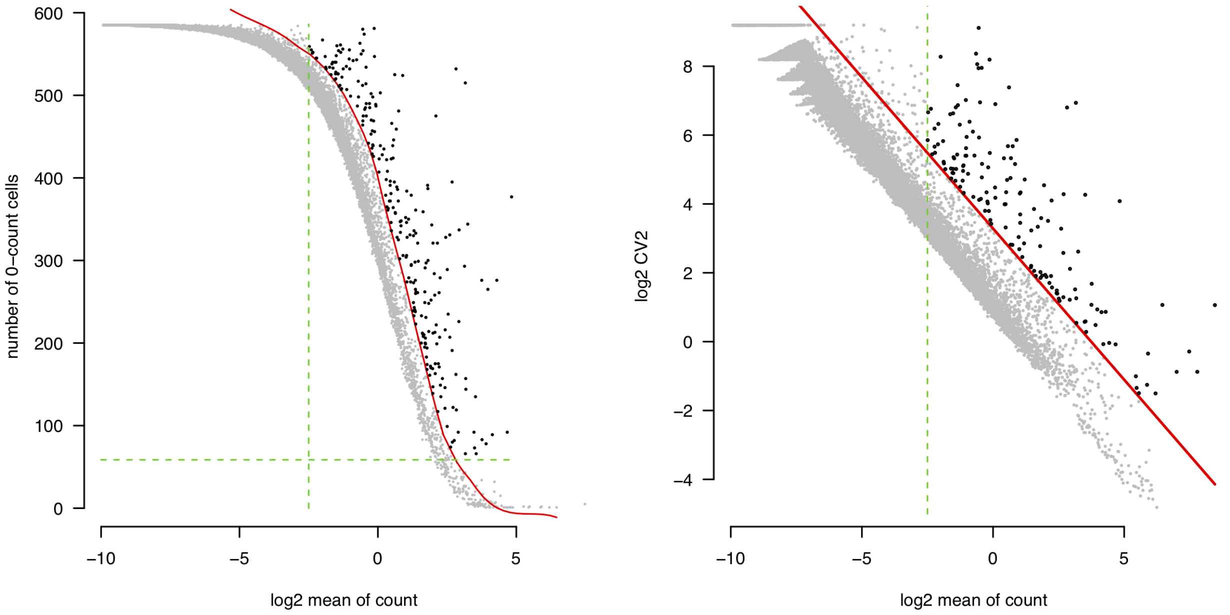


Figure 1: The detection of HVGs in Emb. 05 in blood system. black dots: HVGs, grey dots: all other genes, red line: threshold.

#### 2.2 Iterative clustering

For each developmental system, we performed 1 or 2 rounds of clustering, which is dependent on the complexity of developmental system (see Methods 'parameter optimalization'). A wrap function for iterative clustering was executed for each system.

```
library(Seurat)
blood_res <- r2_wrap(cell = blood_cell, mx = norm_mx, raw_count = raw_mx,
    remove_gene = vis_rmg, path = "../result/blood/", phe_r = 0.5, which_phe = 1)
# r2_wrap: a wrap function for iterative clustering (code avaiable
# online); norm_mx: normalized matrix; raw_mx: raw matrix; vis_rmg:
# genes that are involved in batch effect to be removed; phe_r:
# resolution of 'FindClusters' in Seurat
```

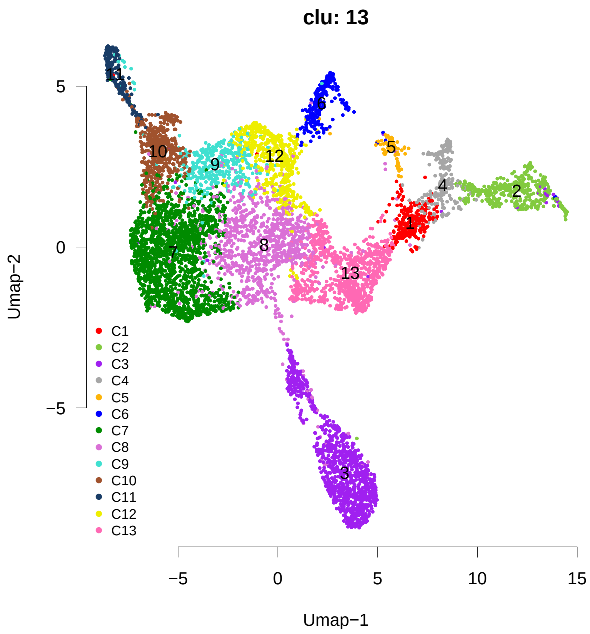


Figure 2: Clusters identified in blood system.

### 3 Annotation

#### 3.1 Marker expression

To annotate each cluster resulted from iterative clustering, two ways of visualization of marker expression were performed, i.e., heatmaps on Umap and dot plot. Heatmaps on Umap provide a global view of the expression of a marker but is hard to be scaled up for many markers. Dot plot provides a global view of all markers in a quantitative way but less intuitive than heatmap on Umap. The combine of two plots could largely increase the efficiency and reliability of annotation.

```
# Read markers of blood system
library(openxlsx)
s1b <- read.xlsx("../result/reclustering3/TJ_clu/Extended Data Table 1 12-19.xlsx",
    sheet = 2, startRow = 2)  # Supplementary table 1B
mk_data <- s1b[s1b[, 2] == "blood", c(1, 4)]
mk <- lapply(mk_data[, 1], function(x) {
```

```r
    genes <- sapply(as.character(mk_data[mk_data[, 1] == x, 2]), function(x) {
        strsplit(x, split = "+", fixed = T)[[1]]
    })
    return(get_id(genes))
})
names(mk) <- mk_data[, 1]
```

```r
# Heatmaps of markers on Umap.  'plot_genes_on_tsne_md': a function for
# plotting a group of genes on Umap (code avaiable online)
for (i in 1:length(mk)) {
    plot_genes_on_tsne_md(tsne = blood_res[[1]][[1]], mx = raw_mx, genes = mk[[i]],
        plot_cex = 0.5, is_order = F, file_name = names(mk)[i])
}
```

```
## [1] "dendritic cell"
```

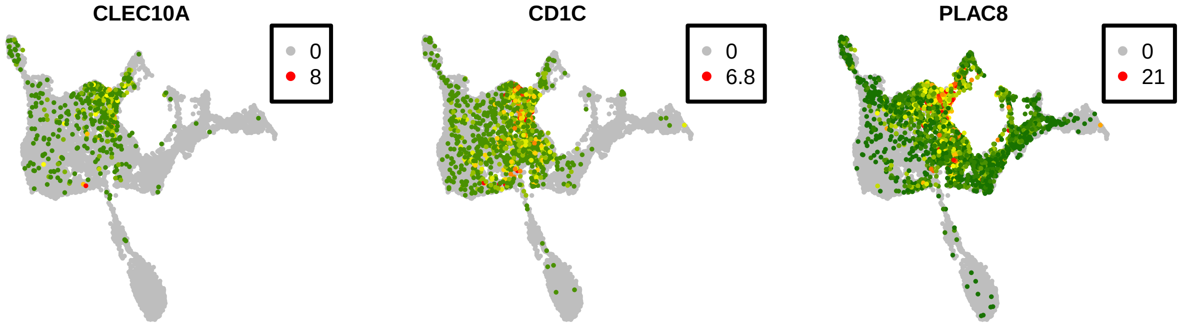


```
## [1] "eosino/basophil/mast cell progenitor"
```

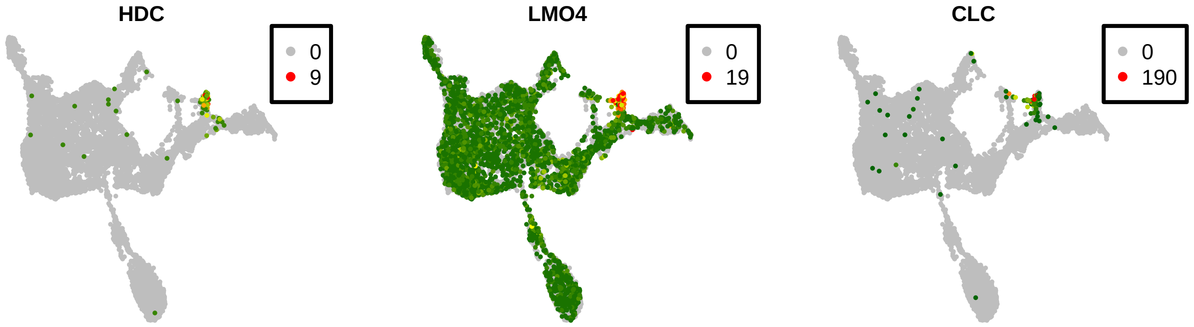


```
## [1] "hematopoietic stem cell"
```

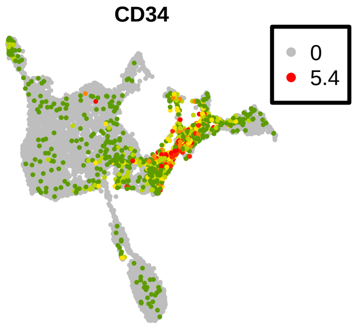


```
## [1] "lymphocyte"
```

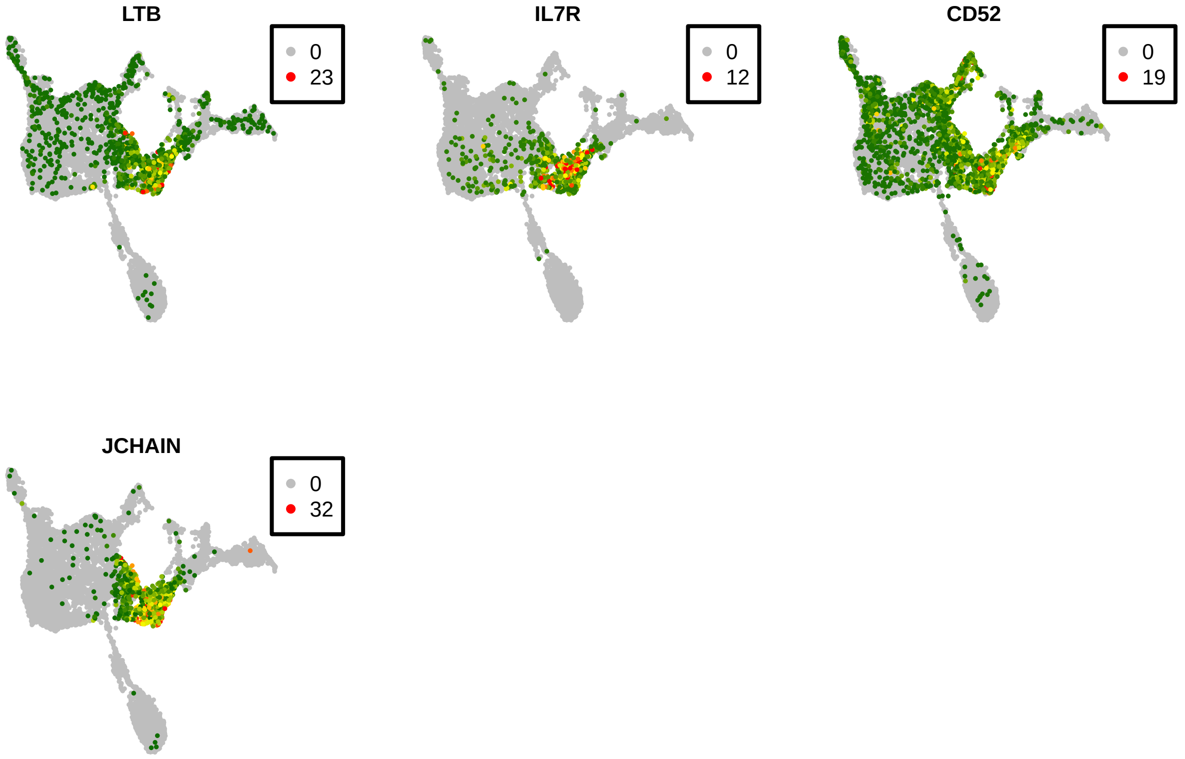


```
## [1] "macrophage"
```

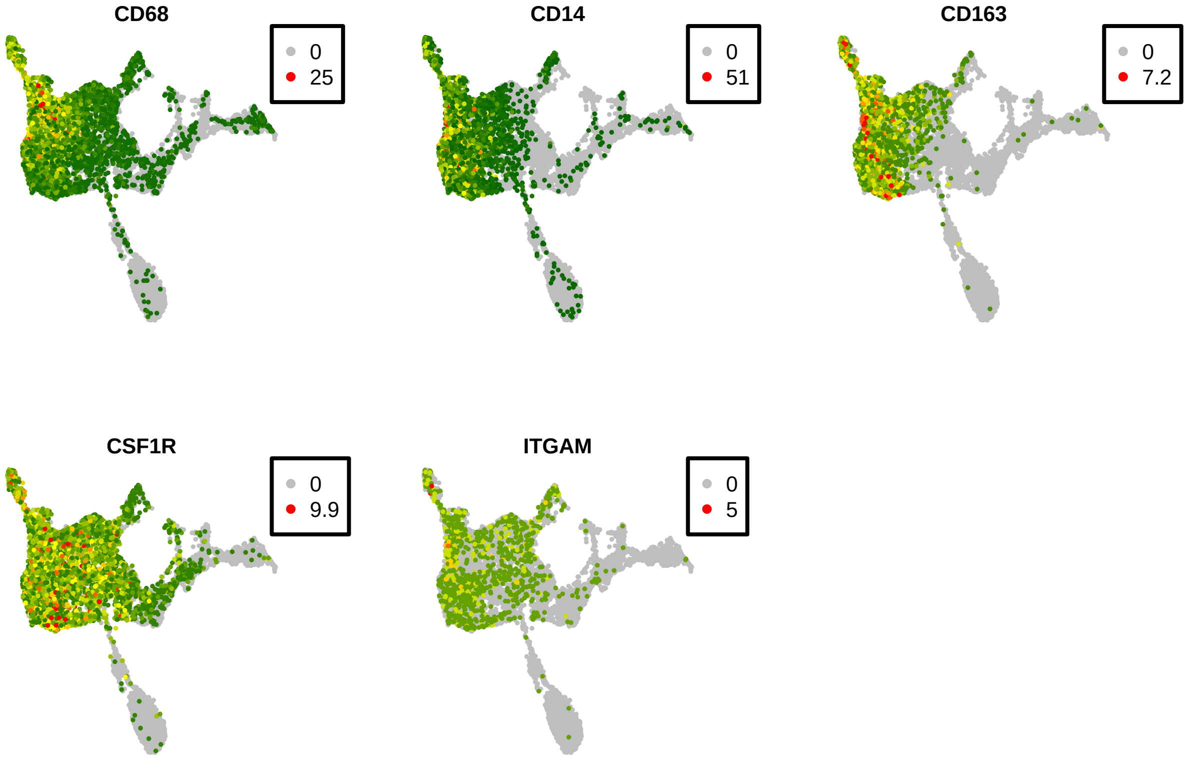


```
## [1] "kupffer cell"
```

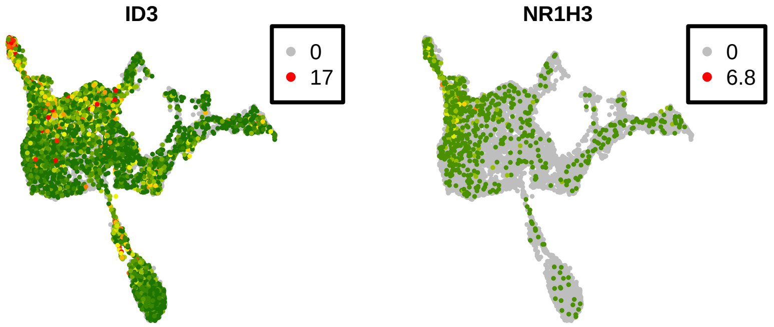


#### [1] "megakaryocyte"

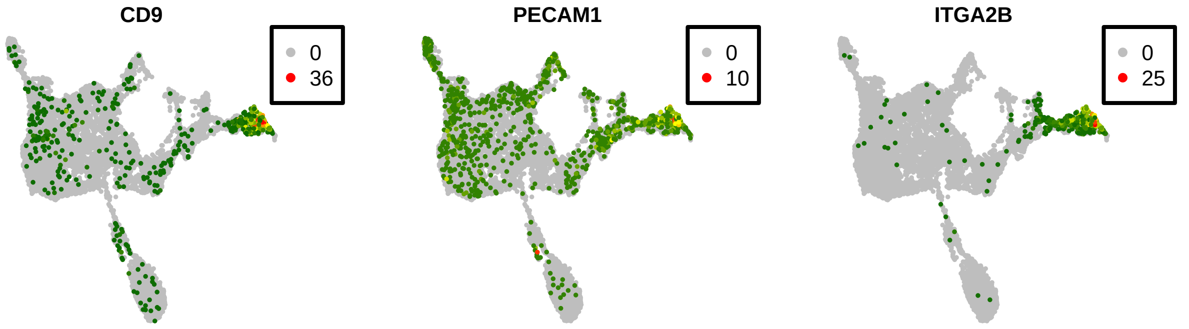


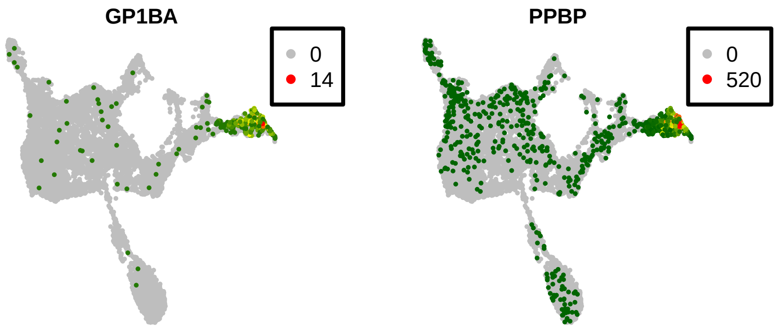


#### [1] "neutrophil"

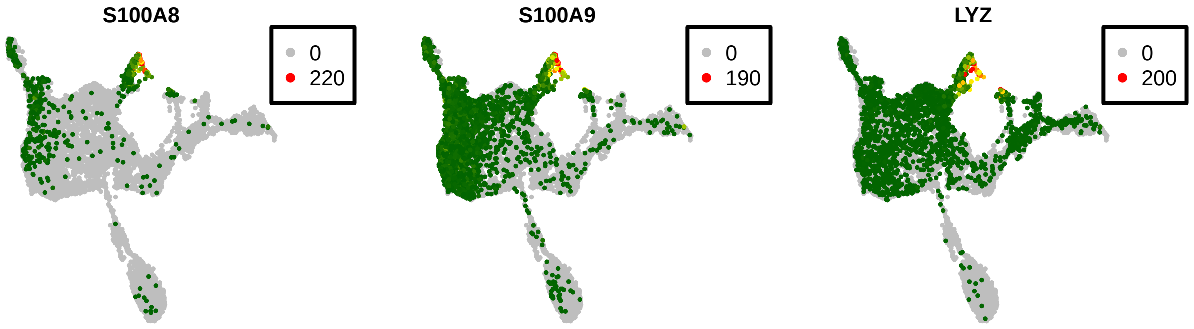


#### [1] "primary neutrophil granules"

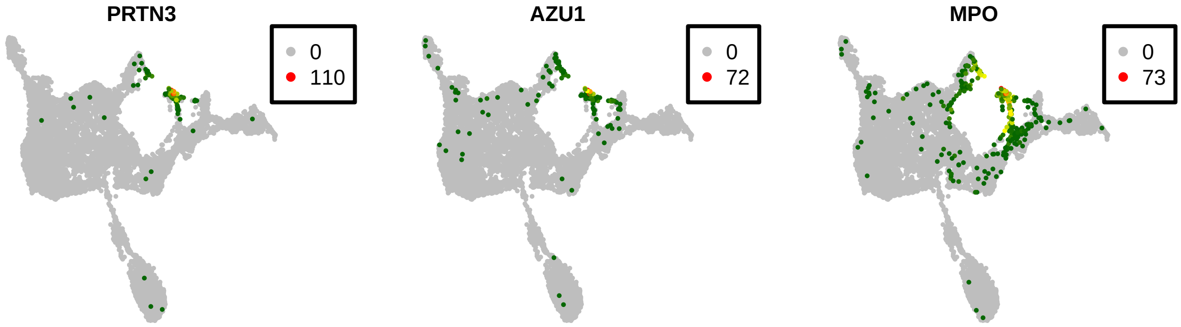


```
## [1] "erythroid"
```

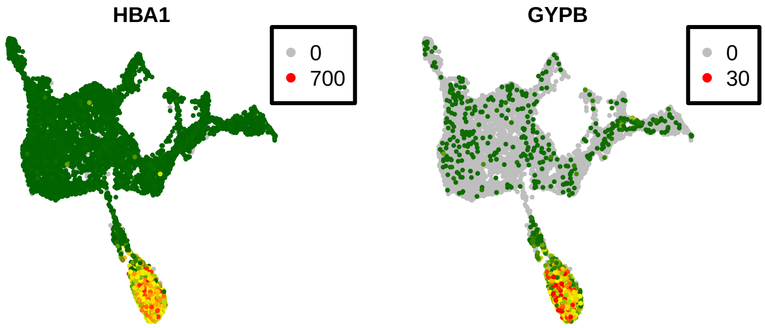


```
# Dot plot for all markers in each cluster.  'DotPlot': a function in
# package Seurat for dot plot
library(ggplot2)
plot_mx <- raw_mx[, rownames(blood_res[[1]][[1]])]
rownames(plot_mx) <- ge2an[rownames(plot_mx)]  # 'ge2an': convert gene IDs to gene symbols
seu <- CreateSeuratObject(plot_mx, min.cells = 0, min.features = 0)
seu <- NormalizeData(seu)
seu@active.ident <- as.factor(blood_res[[1]][[3]][, 1])
DotPlot(seu, features = ge2an[unique(unlist(mk))], col.max = 1) + theme(axis.text.x = element_text(size
    angle = 90, vjust = 0.5, hjust = 1))
```

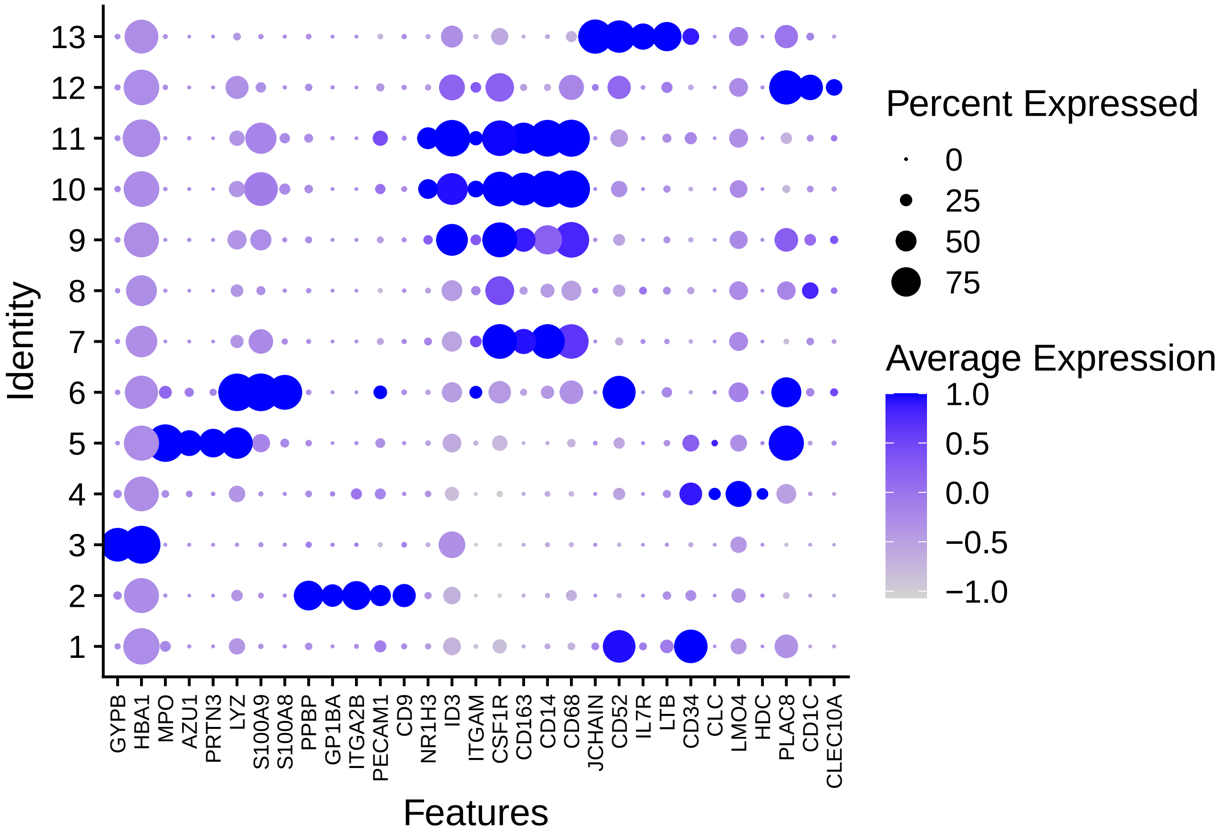


##### 3.2 Assign annotation

Using two types of visualization on marker expression, we can now annotate each cluster with 2-5 markers. For clusters that are annotated by the same cell type, consider them as subtypes named with "cell type - number".

```
# Assign annotation for each cluster according to the expression of
# markers.
anno <- c("hematopoietic stem cell", "megakaryocyte", "erythroid", "eosino/basophil/mast cell progenito
    "primary neutrophil granules", "neutrophil", "macrophage", "macrophage",
    "macrophage", "kupffer cell", "kupffer cell", "dendritic cell", "lymphocyte")
anno2 <- anno
dup_anno <- unique(anno[duplicated(anno)])
for (i in 1:length(dup_anno)) {
    ind <- anno2 == dup_anno[i]
    anno2[ind] <- paste(anno2[ind], 1:sum(ind), sep = "-")
}
print(anno2)
```

```
##  [1] "hematopoietic stem cell"            "megakaryocyte"
##  [3] "erythroid"                          "eosino/basophil/mast cell progenitor"
##  [5] "primary neutrophil granules"        "neutrophil"
##  [7] "macrophage-1"                       "macrophage-2"
##  [9] "macrophage-3"                       "kupffer cell-1"
## [11] "kupffer cell-2"                     "dendritic cell"
## [13] "lymphocyte"
```

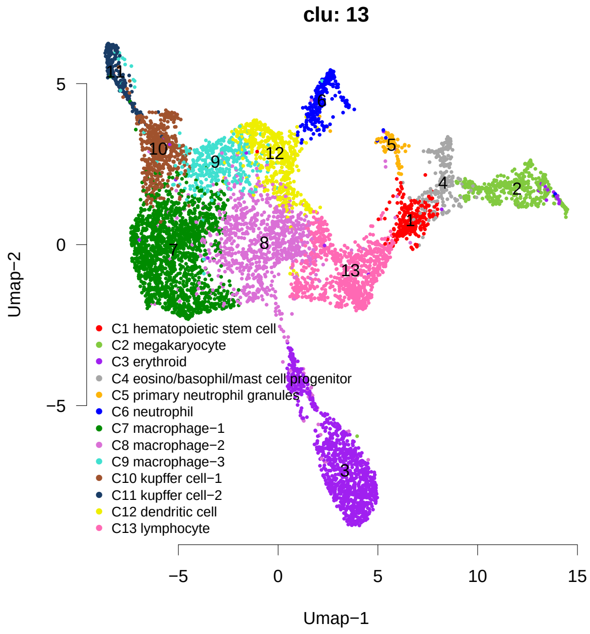


Figure 3: The annotations of clusters identified in blood system.

### 4 Special case of annotation

#### 4.1 Spatial domains of limb bud

The extensive *in situ* data available in limb bud of mouse embryos (https://www.embrys.jp/embrys/html/MainMenu.html) and pseudo-axis information revealed by Umap allows us to determine the spatial location of clusters in limb bud. This procedure is different from the traditional way of annotating cell types showed above. Limb bud at CS13-14 is showed here as a special case of annotation (Figure 4).

We chose genes reported to have functional impact on limb bud development as marker genes and only considered *in situ* data at corresponding stage (e.g., E10.5 for CS13-14). A minimal set of markers with diverse expression pattern were showed here in Figure 4 to explain the procedure of annotating spatial location at CS13-14 (for all markers we used, see Supplementary Figure 10). Like solving a **jigsaw puzzle**, we started with gene with simple expression pattern and continued with genes that are relevant to annotated clusters. The stepwise annotation is explained below.

a) The Umap of cells of forelimb mesenchyme at CS13-14 and the summary of spatial domains we resolved (Figure 4a).

b) *EMX2 in situ* indicates that cluster $a$ is at the most proximal part of limb bud (Figure 4b).

c) *PBX1 in situ* indicates that the spatial locations of cluster $d$ and $e$ are next to cluster $a$ (Figure 4c).

d) *MSX2 in situ* could distinguish cluster $d$ from cluster $e$ and indicates that the spatial location of cluster $h$ is next to cluster $d$ on the anterior side (Figure 4d).

e) *SHH* is well-known to be expressed in ZPA (zone of polarizing activity), which is at the posterior part of limb bud, indicating the location of cluster $z$ (Figure 4e).

f) *LMO1 in situ* indicates that cluster $p$ and $q$ is at the distal part of limb bud (Figure 4f).

g) *HAND2 in situ* indicates shows that it is expressed in the posterior, which separates cluster $q$ from $p$ and also suggests that $f$ and $j$ are proximal to $z$. Furthermore, the low expression indicates cluster $i$, to be in the middle (Figure 4g).

h) *NR2F2 in situ* distinguishes the location of cluster $f$ from cluster $j$ (Figure 4h).

Taking together, the spatial domains of limb bud at CS13 were summarized in Figure 4a (right). The spatial map is *post hoc* tested by the expression of HOX genes (see main text).

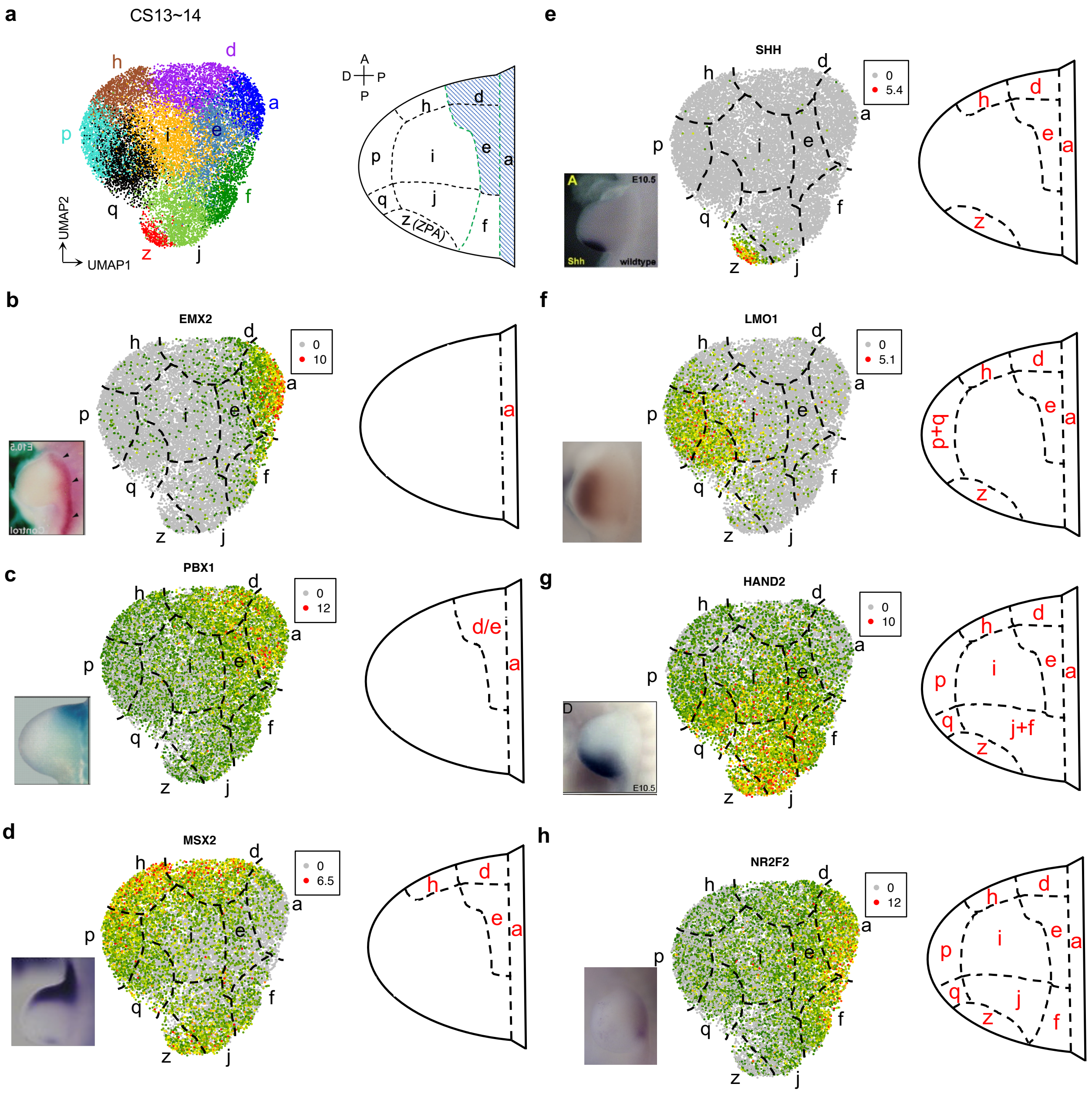


Figure 4: Stepwise annotate spatial location of clusters in limb bud at CS13-14. (a) Umap of limb bud at CS13-14 colored by 10 clusters (left) and spatial locations of clusters annotated by marker genes (right). (b-h) Stepwise annotate spatial location of each cluster. From left to right, *in situ* data, heatmap of gene expression, and annotated clusters (also see Supplementary Figure 10).

#### 4.2 The comparison of our reconstruction at CS15-16 and spatial transcriptome of human limb bud at 5.6 weeks

To further test our reconstruction of spatial domains in limb bud, we compared the expression of 20 patterning genes in our reconstruction to spatial transcriptome of human limb bud at similar stage (Zhang et al. 2022, bioRxiv). These genes were independently selected by Zhang and colleagues. 17 out of 20 genes (*e.g.*, *HAND1*, *ZIC3*, *IRX3*) show consistent expression pattern between two datasets, indicating the high fidelity of our spatial reconstruction (Figure 5).

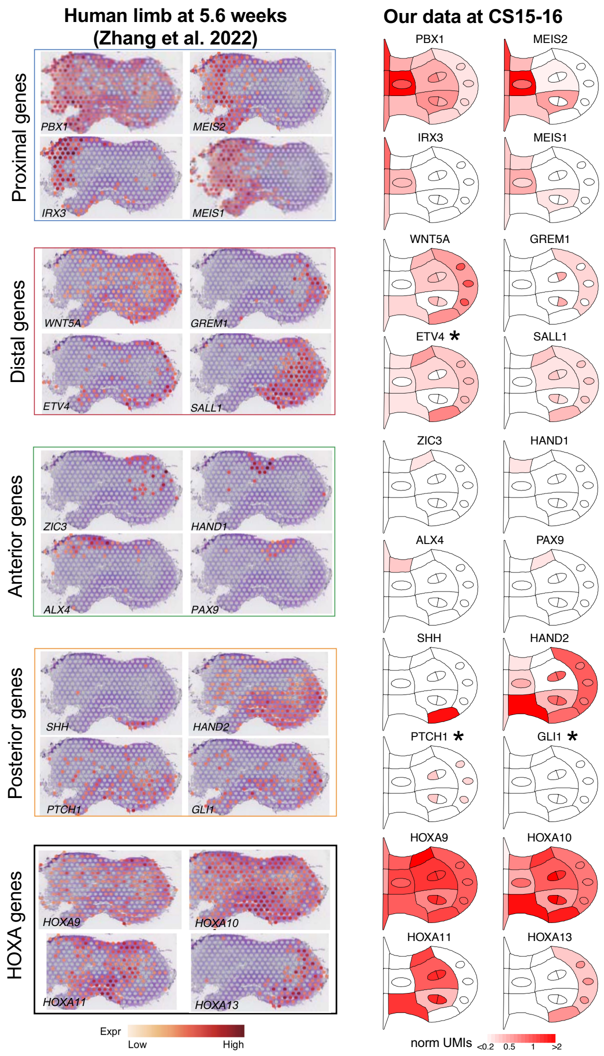


Figure 5: The comparison of our reconstruction at CS15-16 and spatial transcriptome of human limb bud at 5.6 weeks (Zhang et al. 2022, bioRxiv). The expression of 20 genes from 5 groups selected by Zhang and colleagues are showed, *i.e.*, proximal, distal, anterior, posterior, and HOXA genes. Asterisks denote genes that are inconsistent between two datasets.

```
# Version of packages
sessionInfo()
```

```
## R version 3.6.3 (2020-02-29)
## Platform: x86_64-apple-darwin15.6.0 (64-bit)
## Running under: macOS Catalina 10.15.7
```

```
##
## Matrix products: default
## BLAS:   /Library/Frameworks/R.framework/Versions/3.6/Resources/lib/libRblas.0.dylib
## LAPACK: /Library/Frameworks/R.framework/Versions/3.6/Resources/lib/libRlapack.dylib
##
## locale:
## [1] en_US.UTF-8/en_US.UTF-8/en_US.UTF-8/C/en_US.UTF-8/en_US.UTF-8
##
## attached base packages:
## [1] stats     graphics  grDevices utils     datasets  methods   base
##
## other attached packages:
## [1] ggplot2_3.3.5 rmarkdown_2.7 Seurat_3.1.5
##
## loaded via a namespace (and not attached):
##  [1] nlme_3.1-144       tsne_0.1-3         matrixStats_0.58.0 RcppAnnoy_0.0.18   RColorBrewer_1.1-2
##  [6] httr_1.4.2         sctransform_0.3.2  tools_3.6.3        utf8_1.2.1         R6_2.5.0
## [11] irlba_2.3.3        KernSmooth_2.23-16 uwot_0.1.10        DBI_1.1.1          lazyeval_0.2.2
## [16] colorspace_2.0-0   withr_2.4.1        tidyselect_1.1.0   gridExtra_2.3      compiler_3.6.3
## [21] cli_3.1.1          formatR_1.9        plotly_4.9.3       labeling_0.4.2     scales_1.1.1
## [26] lmtest_0.9-38      ggridges_0.5.3     pbapply_1.4-3      stringr_1.4.0      digest_0.6.27
## [31] pkgconfig_2.0.3    htmltools_0.5.1.1  parallelly_1.24.0  highr_0.8          htmlwidgets_1.5.3
## [36] rlang_1.0.1        generics_0.1.0     farver_2.1.0       zoo_1.8-9          jsonlite_1.7.2
## [41] ica_1.0-2          dplyr_1.0.5        magrittr_2.0.2     patchwork_1.1.1    Matrix_1.3-2
## [46] Rcpp_1.0.6         munsell_0.5.0      fansi_0.4.2        ape_5.4-1          reticulate_1.18
## [51] lifecycle_1.0.0    stringi_1.5.3      yaml_2.2.1         MASS_7.3-51.5      Rtsne_0.15
## [56] plyr_1.8.6         grid_3.6.3         parallel_3.6.3     listenv_0.8.0      ggrepel_0.9.1
## [61] crayon_1.4.1       lattice_0.20-38    cowplot_1.1.1      splines_3.6.3      knitr_1.37
## [66] pillar_1.7.0       igraph_1.2.6       future.apply_1.7.0 reshape2_1.4.4     codetools_0.2-16
## [71] leiden_0.3.7       glue_1.6.1         evaluate_0.14      data.table_1.13.6  png_0.1-7
## [76] vctrs_0.3.8        gtable_0.3.0       RANN_2.6.1         purrr_0.3.4        tidyr_1.1.3
## [81] future_1.21.0      assertthat_0.2.1   xfun_0.30          rsvd_1.0.3         survival_3.1-8
## [86] viridisLite_0.3.0  tibble_3.1.0       cluster_2.1.0      globals_0.14.0     fitdistrplus_1.1-3
## [91] ellipsis_0.3.2     ROCR_1.0-11
```
