## Supplementary Note 2 for "A single-cell transcriptome atlas of human early embryogenesis"

neural progenitor−1: anteromedial cerebral pole (ACP)

neural progenitor−2: ventral telencephalon−1

neural progenitor−3: ventral telencephalon−2

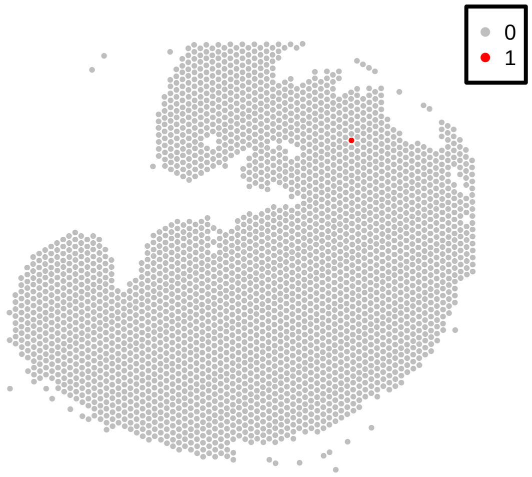


neural progenitor−4: ventral telencephalon−3

neural progenitor−5: GABAergic neuron precursor

neural progenitor−6: anterior head fold

neural progenitor−7: dorsal telencephalon−1

neural progenitor−8: dorsal telencephalon−2

neural progenitor−9: dorsal telencephalon−3

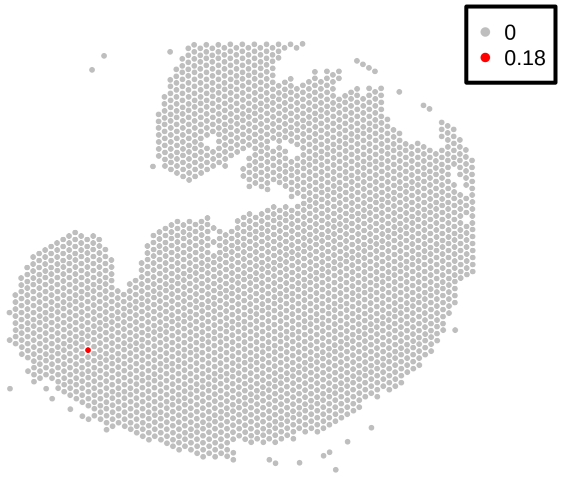


neural progenitor−10: dorsal telencephalon−4

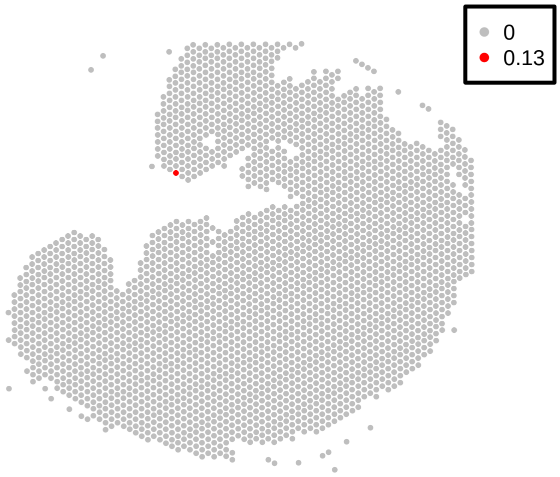


neural progenitor−11: dorsal diencephalon

neural progenitor−12: ventral diencephalon and ZLI

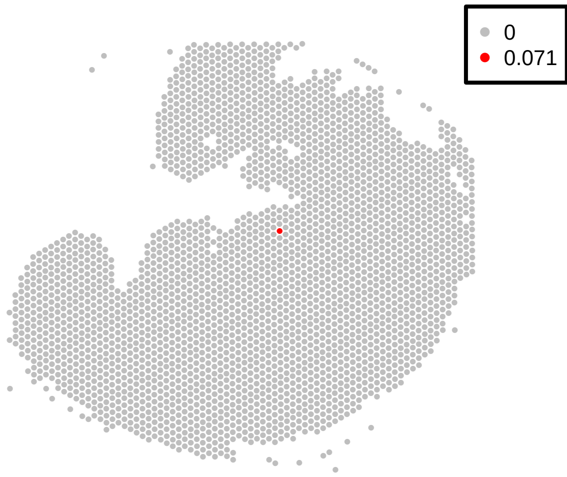

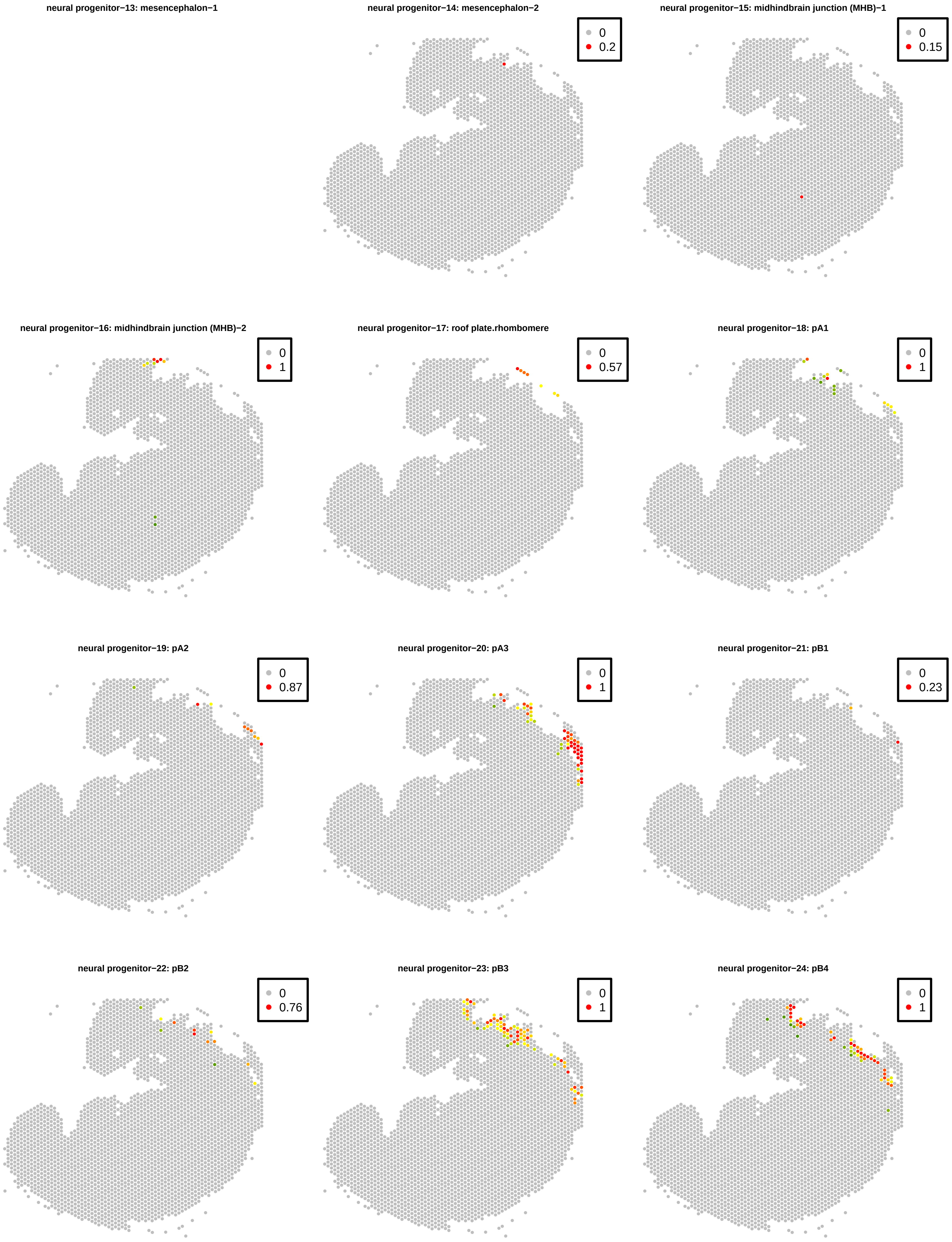

neural progenitor−13: mesencephalon−1
neural progenitor−14: mesencephalon−2
0
0.2
neural progenitor−15: midhindbrain junction (MHB)−1
0
0.15
neural progenitor−16: midhindbrain junction (MHB)−2
0
1
neural progenitor−17: roof plate.rhombomere
0
0.57
neural progenitor−18: pA1
0
1
neural progenitor−19: pA2
0
0.87
neural progenitor−20: pA3
0
1
neural progenitor−21: pB1
0
0.23
neural progenitor−22: pB2
0
0.76
neural progenitor−23: pB3
0
1
neural progenitor−24: pB4
0
1

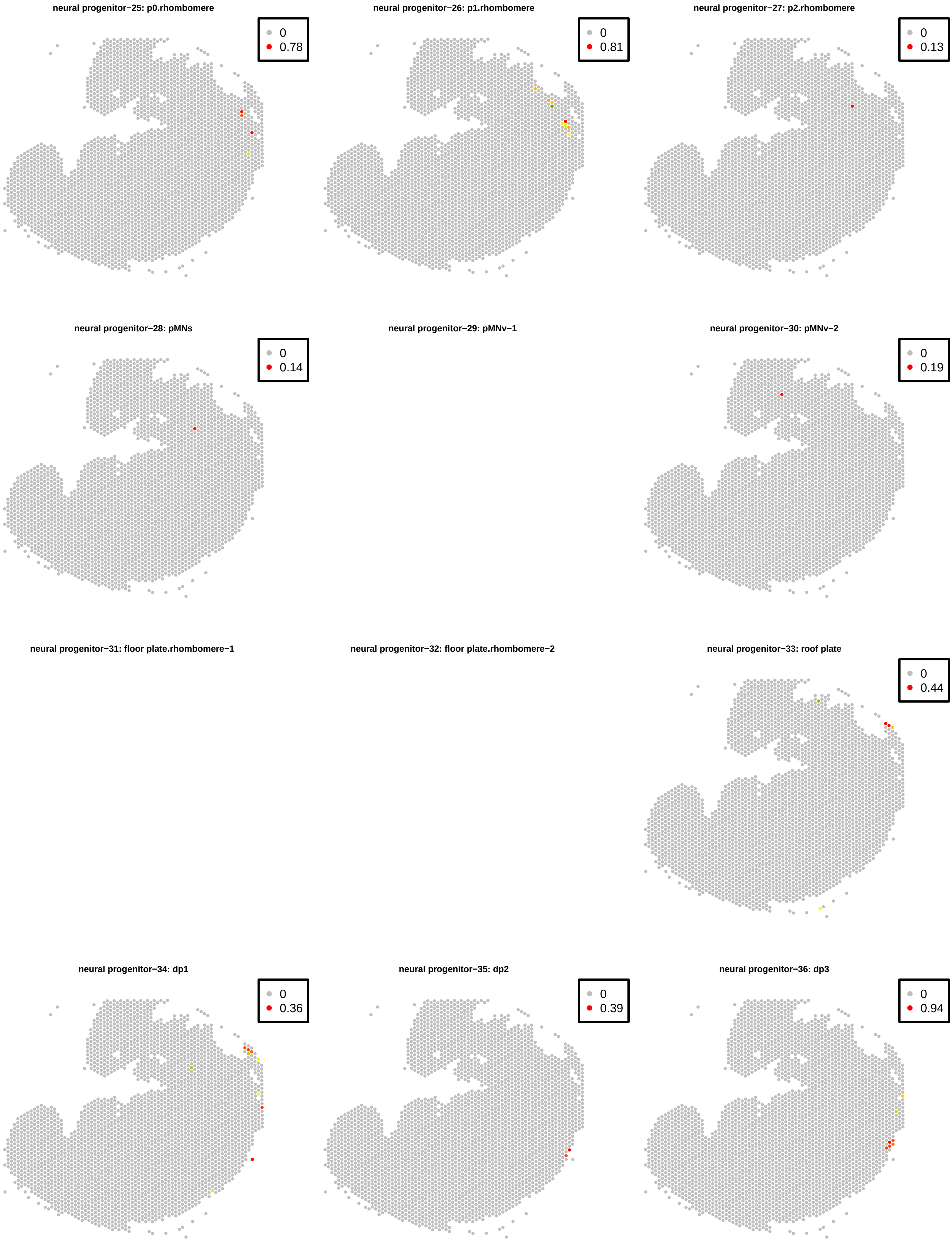
neural progenitor−25: p0.rhombomere
0
0.78
neural progenitor−26: p1.rhombomere
0
0.81
neural progenitor−27: p2.rhombomere
0
0.13
neural progenitor−28: pMNs
0
0.14
neural progenitor−29: pMNv−1
neural progenitor−30: pMNv−2
0
0.19
neural progenitor−31: floor plate.rhombomere−1
neural progenitor−32: floor plate.rhombomere−2
neural progenitor−33: roof plate
0
0.44
neural progenitor−34: dp1
0
0.36
neural progenitor−35: dp2
0
0.39
neural progenitor−36: dp3
0
0.94

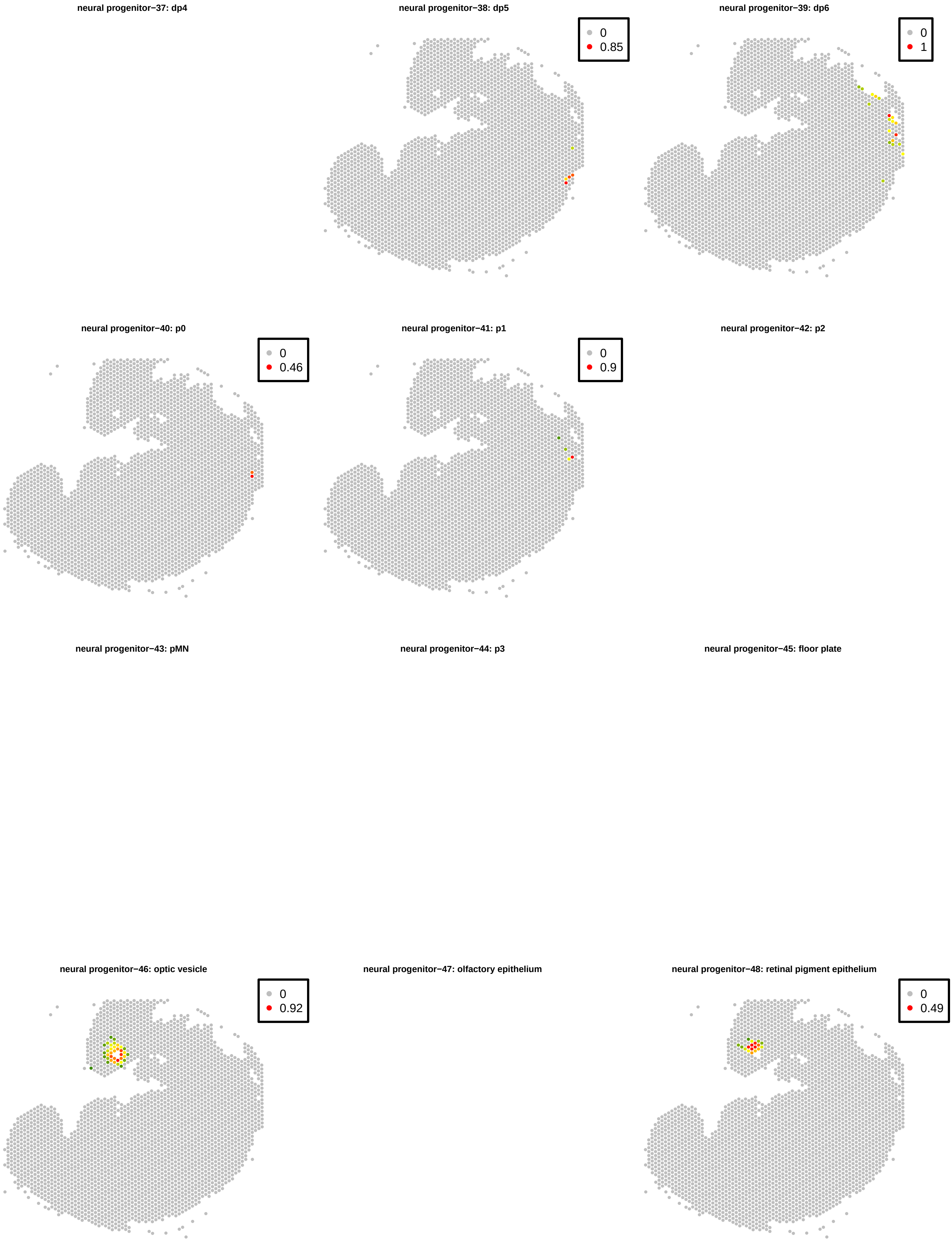

neural progenitor−37: dp4
neural progenitor−38: dp5
0
0.85
neural progenitor−39: dp6
0
1
neural progenitor−40: p0
0
0.46
neural progenitor−41: p1
0
0.9
neural progenitor−42: p2
neural progenitor−43: pMN
neural progenitor−44: p3
neural progenitor−45: floor plate
neural progenitor−46: optic vesicle
0
0.92
neural progenitor−47: olfactory epithelium
neural progenitor−48: retinal pigment epithelium
0
0.49

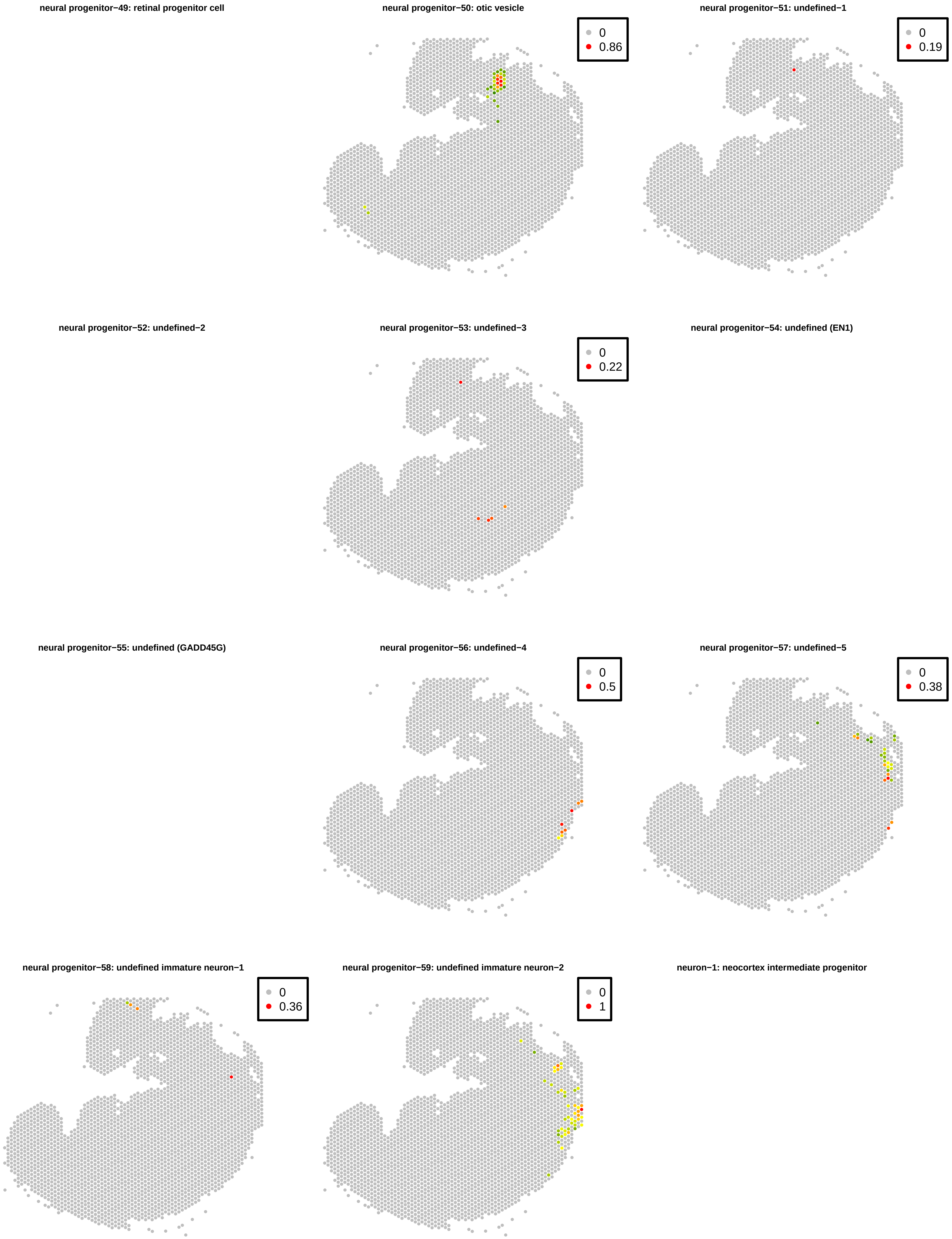
neural progenitor−49: retinal progenitor cell
neural progenitor−50: otic vesicle
0
0.86
neural progenitor−51: undefined−1
0
0.19
neural progenitor−52: undefined−2
neural progenitor−53: undefined−3
0
0.22
neural progenitor−54: undefined (EN1)
neural progenitor−55: undefined (GADD45G)
neural progenitor−56: undefined−4
0
0.5
neural progenitor−57: undefined−5
0
0.38
neural progenitor−58: undefined immature neuron−1
0
0.36
neural progenitor−59: undefined immature neuron−2
0
1
neuron−1: neocortex intermediate progenitor

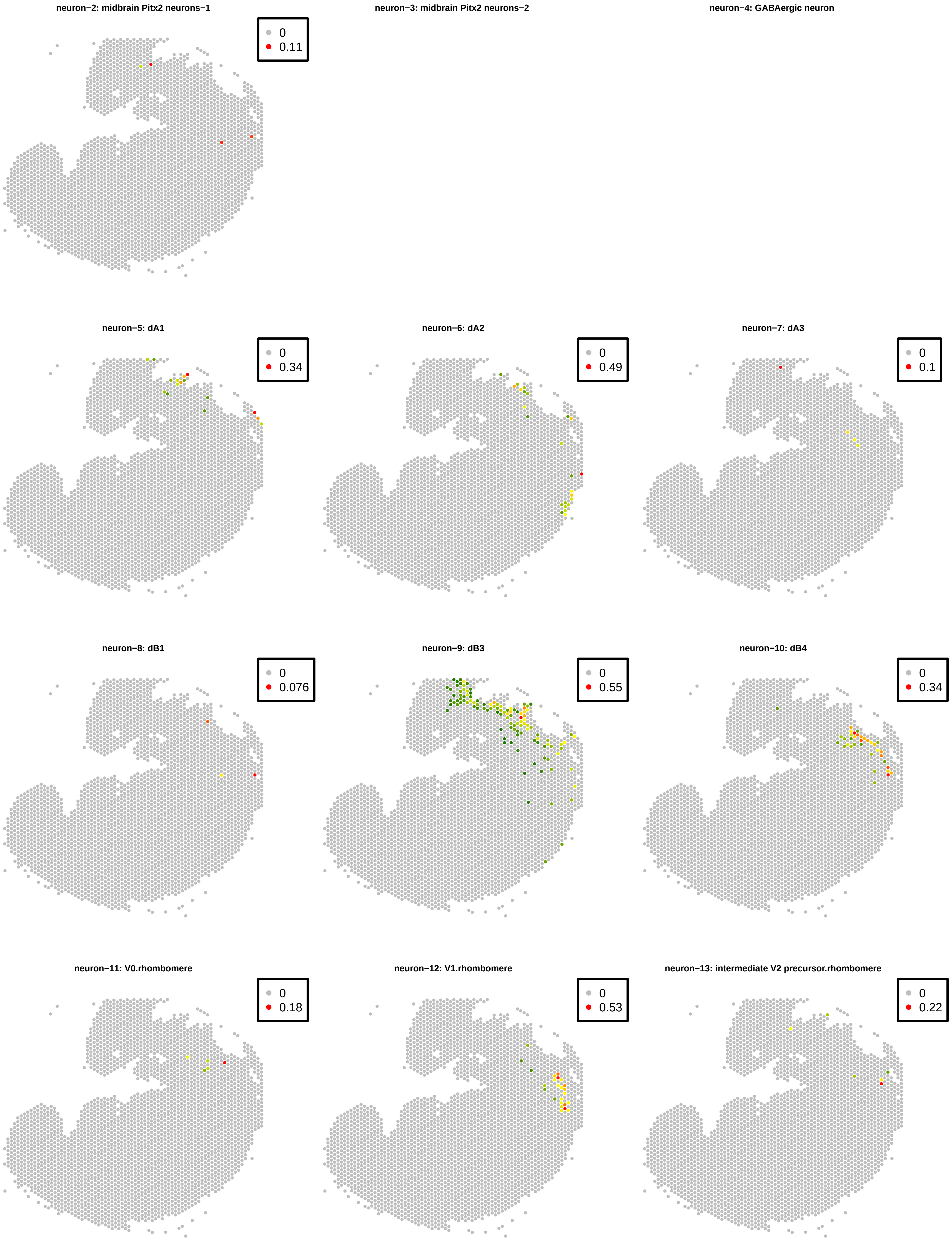
neuron−2: midbrain Pitx2 neurons−1
0
0.11
neuron−3: midbrain Pitx2 neurons−2
neuron−4: GABAergic neuron
neuron−5: dA1
0
0.34
neuron−6: dA2
0
0.49
neuron−7: dA3
0
0.1
neuron−8: dB1
0
0.076
neuron−9: dB3
0
0.55
neuron−10: dB4
0
0.34
neuron−11: V0.rhombomere
0
0.18
neuron−12: V1.rhombomere
0
0.53
neuron−13: intermediate V2 precursor.rhombomere
0
0.22

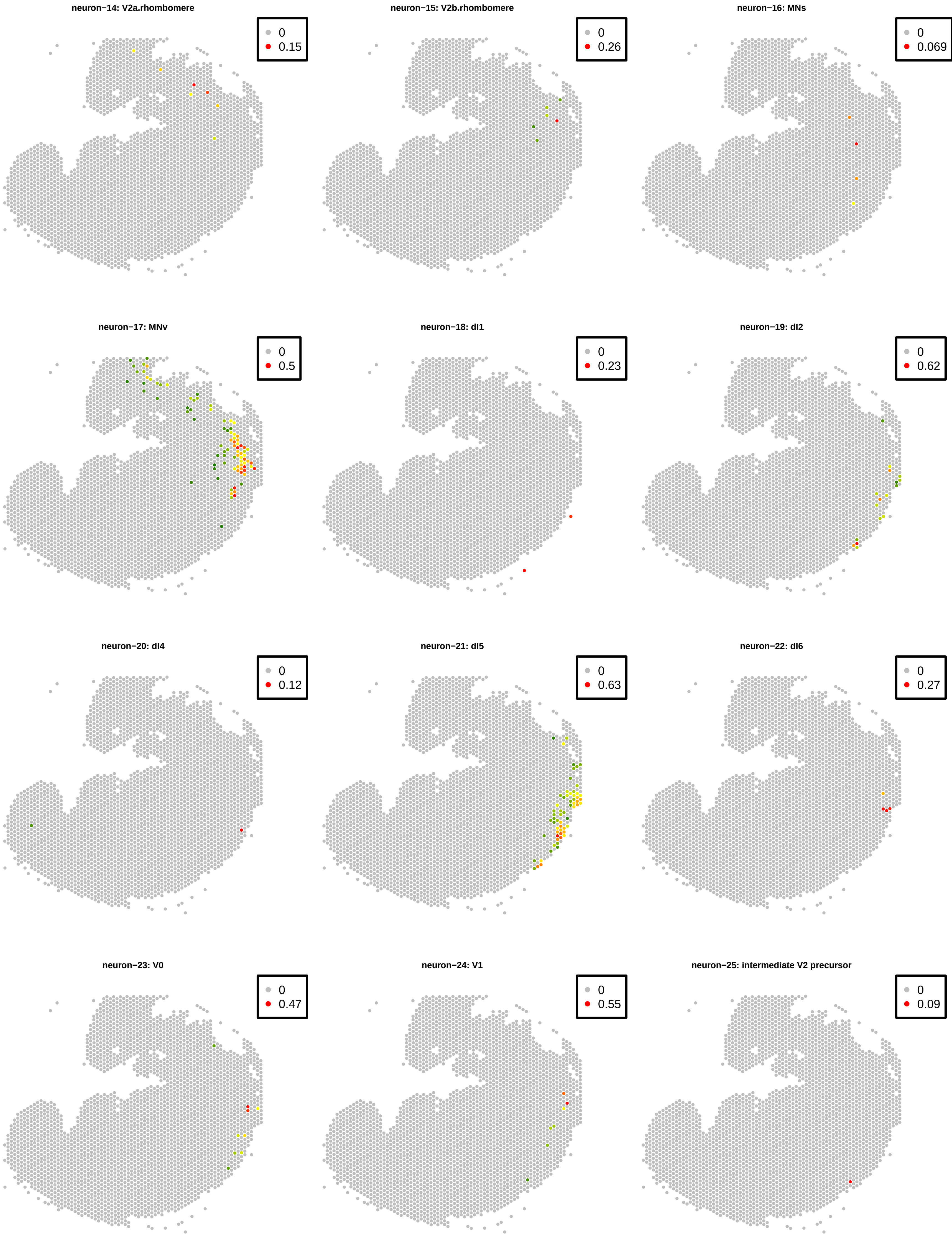
neuron−14: V2a.rhombomere
0
0.15
neuron−15: V2b.rhombomere
0
0.26
neuron−16: MNs
0
0.069
neuron−17: MNv
0
0.5
neuron−18: dI1
0
0.23
neuron−19: dI2
0
0.62
neuron−20: dI4
0
0.12
neuron−21: dI5
0
0.63
neuron−22: dI6
0
0.27
neuron−23: V0
0
0.47
neuron−24: V1
0
0.55
neuron−25: intermediate V2 precursor
0
0.09

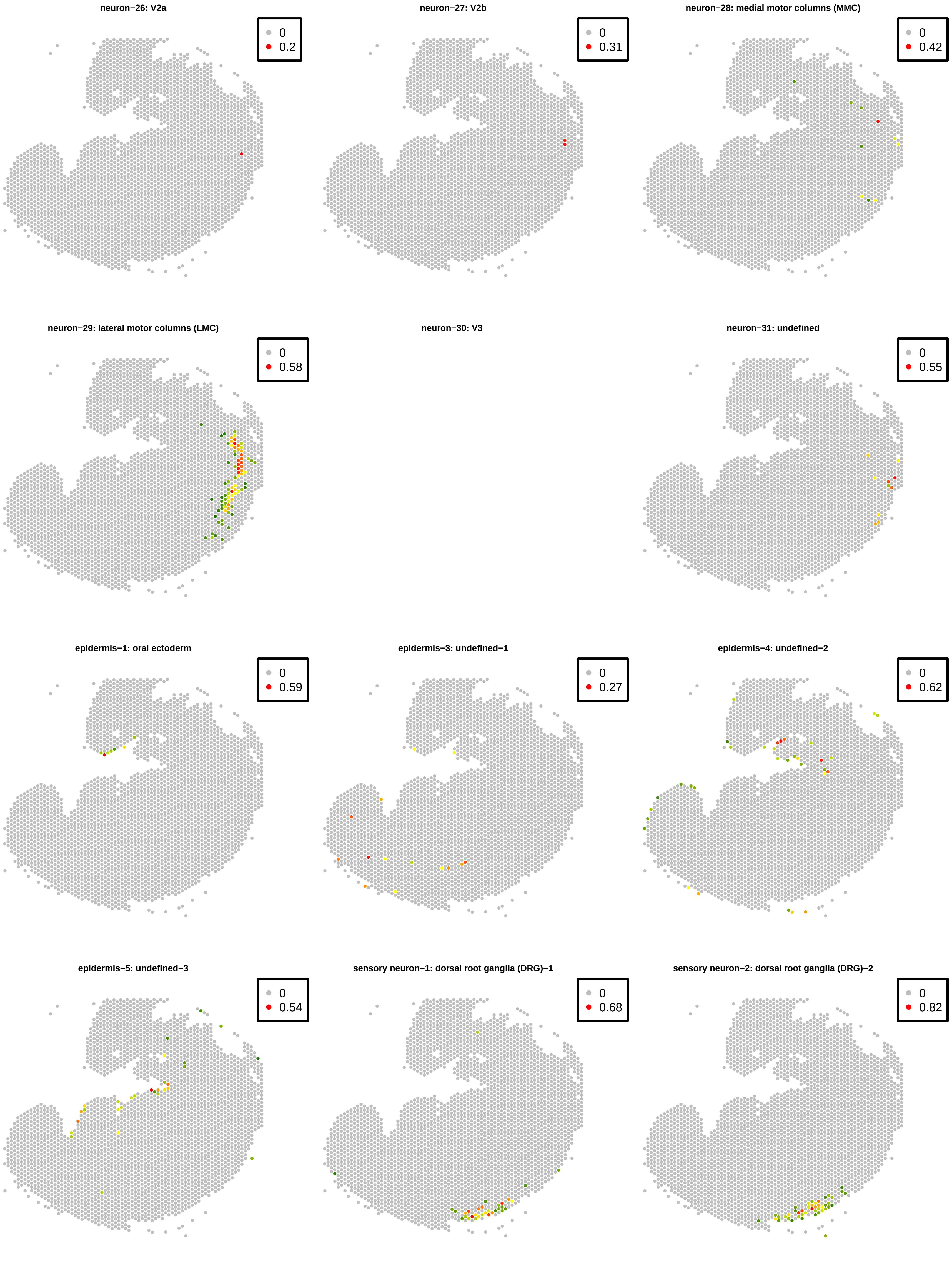
neuron−26: V2a
0
0.2
neuron−27: V2b
0
0.31
neuron−28: medial motor columns (MMC)
0
0.42
neuron−29: lateral motor columns (LMC)
0
0.58
neuron−30: V3
neuron−31: undefined
0
0.55
epidermis−1: oral ectoderm
0
0.59
epidermis−3: undefined−1
0
0.27
epidermis−4: undefined−2
0
0.62
epidermis−5: undefined−3
0
0.54
sensory neuron−1: dorsal root ganglia (DRG)−1
0
0.68
sensory neuron−2: dorsal root ganglia (DRG)−2
0
0.82

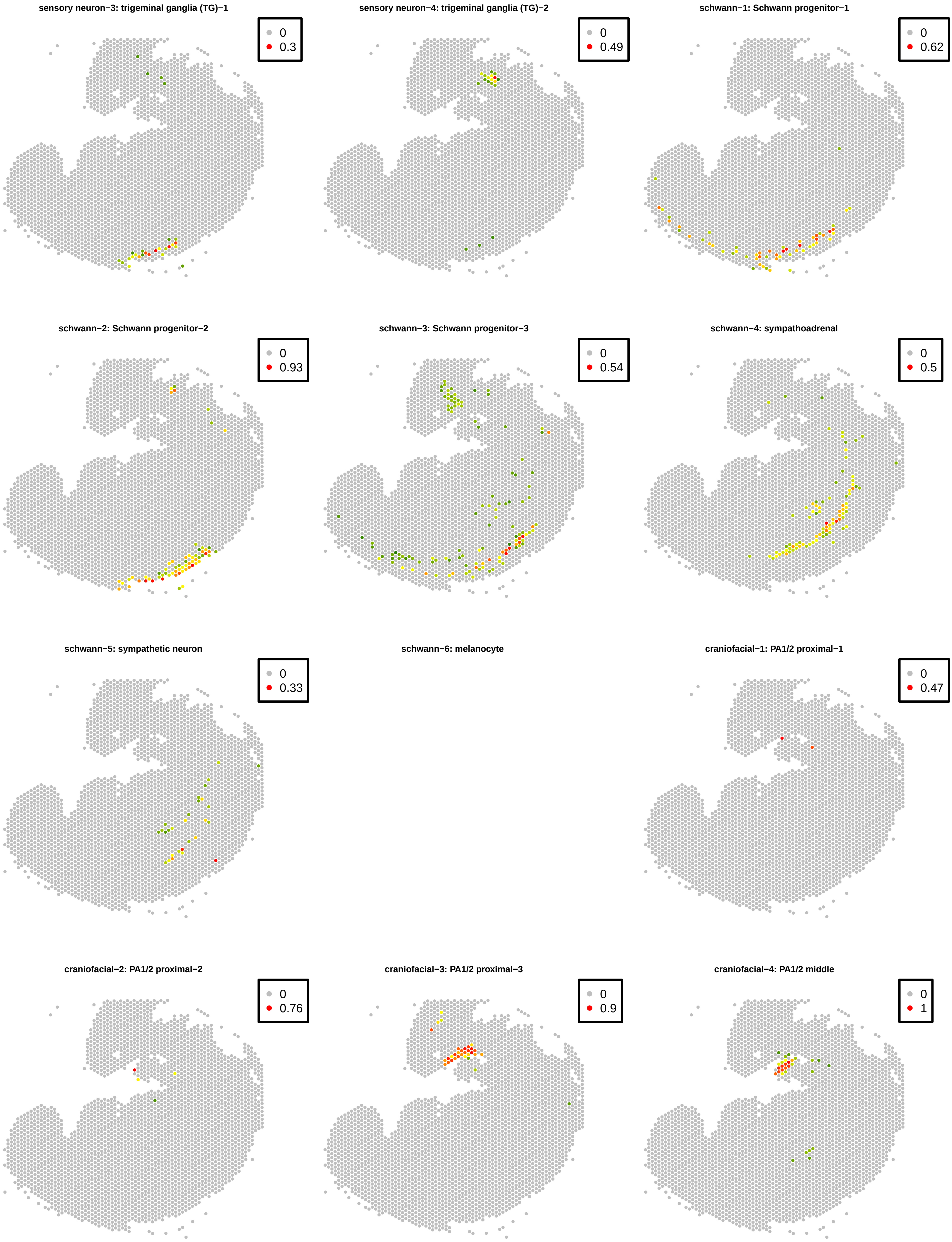
sensory neuron−3: trigeminal ganglia (TG)−1
0
0.3
sensory neuron−4: trigeminal ganglia (TG)−2
0
0.49
schwann−1: Schwann progenitor−1
0
0.62
schwann−2: Schwann progenitor−2
0
0.93
schwann−3: Schwann progenitor−3
0
0.54
schwann−4: sympathoadrenal
0
0.5
schwann−5: sympathetic neuron
0
0.33
schwann−6: melanocyte
craniofacial−1: PA1/2 proximal−1
0
0.47
craniofacial−2: PA1/2 proximal−2
0
0.76
craniofacial−3: PA1/2 proximal−3
0
0.9
craniofacial−4: PA1/2 middle
0
1

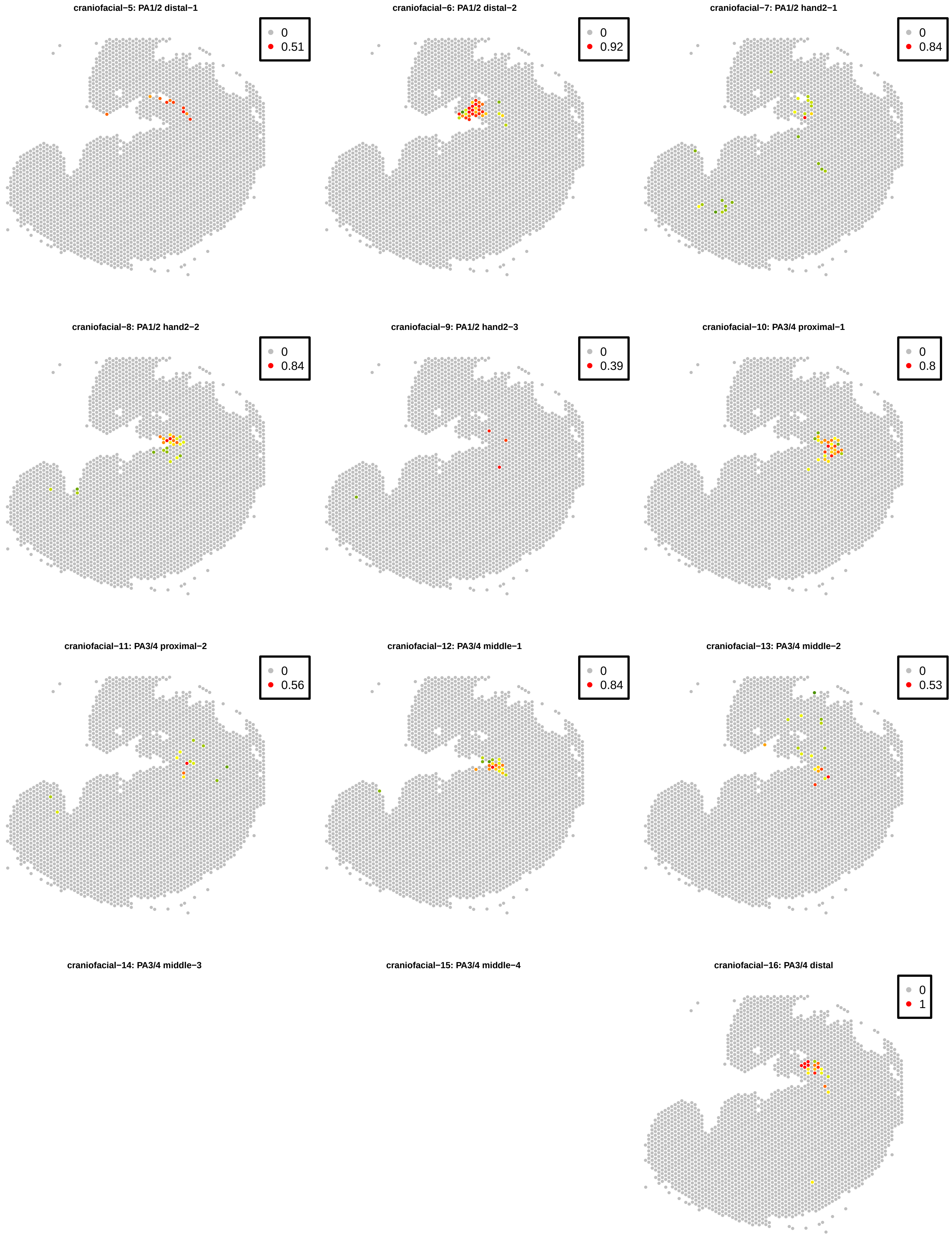
craniofacial−5: PA1/2 distal−1
0
0.51
craniofacial−6: PA1/2 distal−2
0
0.92
craniofacial−7: PA1/2 hand2−1
0
0.84
craniofacial−8: PA1/2 hand2−2
0
0.84
craniofacial−9: PA1/2 hand2−3
0
0.39
craniofacial−10: PA3/4 proximal−1
0
0.8
craniofacial−11: PA3/4 proximal−2
0
0.56
craniofacial−12: PA3/4 middle−1
0
0.84
craniofacial−13: PA3/4 middle−2
0
0.53
craniofacial−14: PA3/4 middle−3
craniofacial−15: PA3/4 middle−4
craniofacial−16: PA3/4 distal
0
1

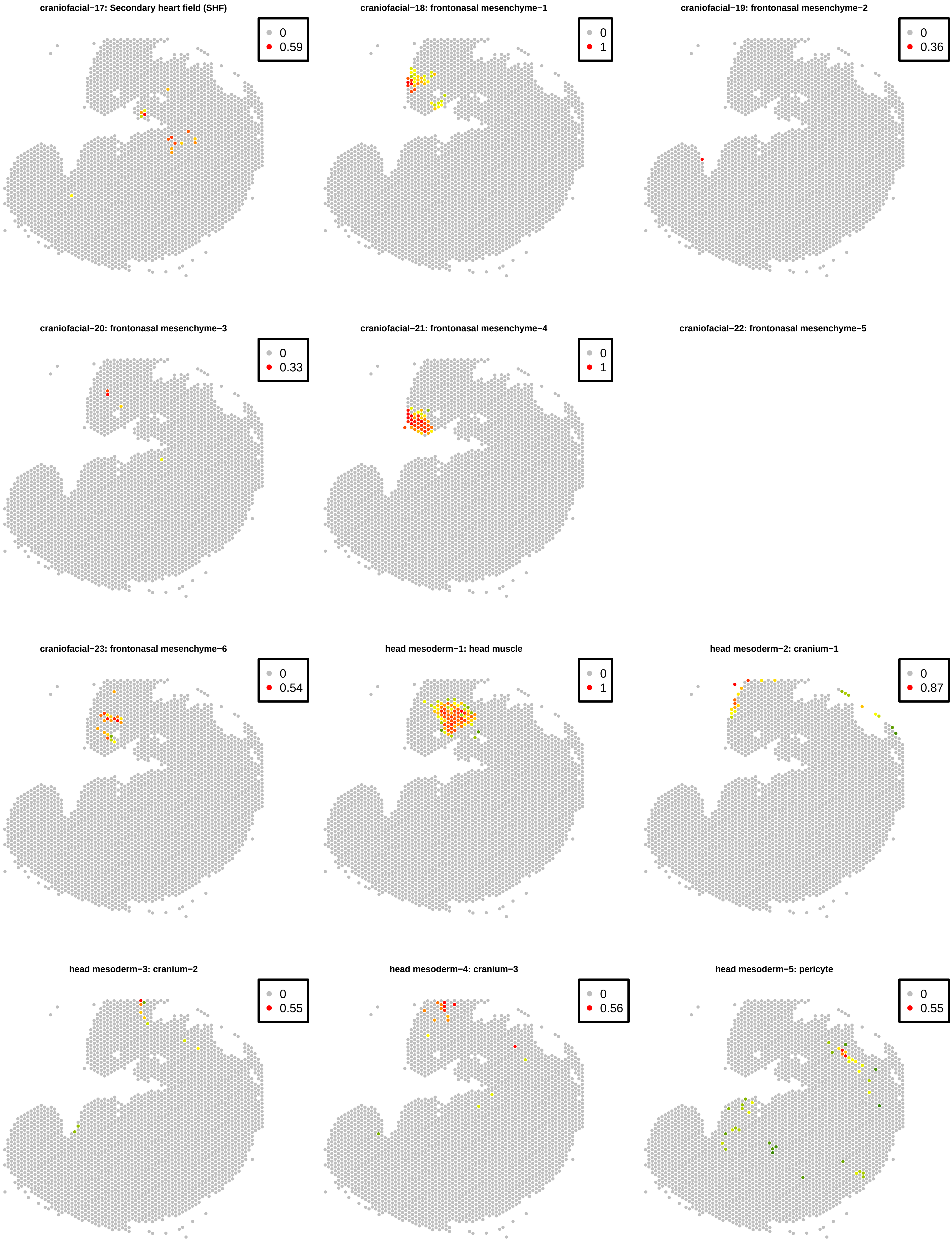
craniofacial−17: Secondary heart field (SHF)
0
0.59
craniofacial−18: frontonasal mesenchyme−1
0
1
craniofacial−19: frontonasal mesenchyme−2
0
0.36
craniofacial−20: frontonasal mesenchyme−3
0
0.33
craniofacial−21: frontonasal mesenchyme−4
0
1
craniofacial−22: frontonasal mesenchyme−5
craniofacial−23: frontonasal mesenchyme−6
0
0.54
head mesoderm−1: head muscle
0
1
head mesoderm−2: cranium−1
0
0.87
head mesoderm−3: cranium−2
0
0.55
head mesoderm−4: cranium−3
0
0.56
head mesoderm−5: pericyte
0
0.55

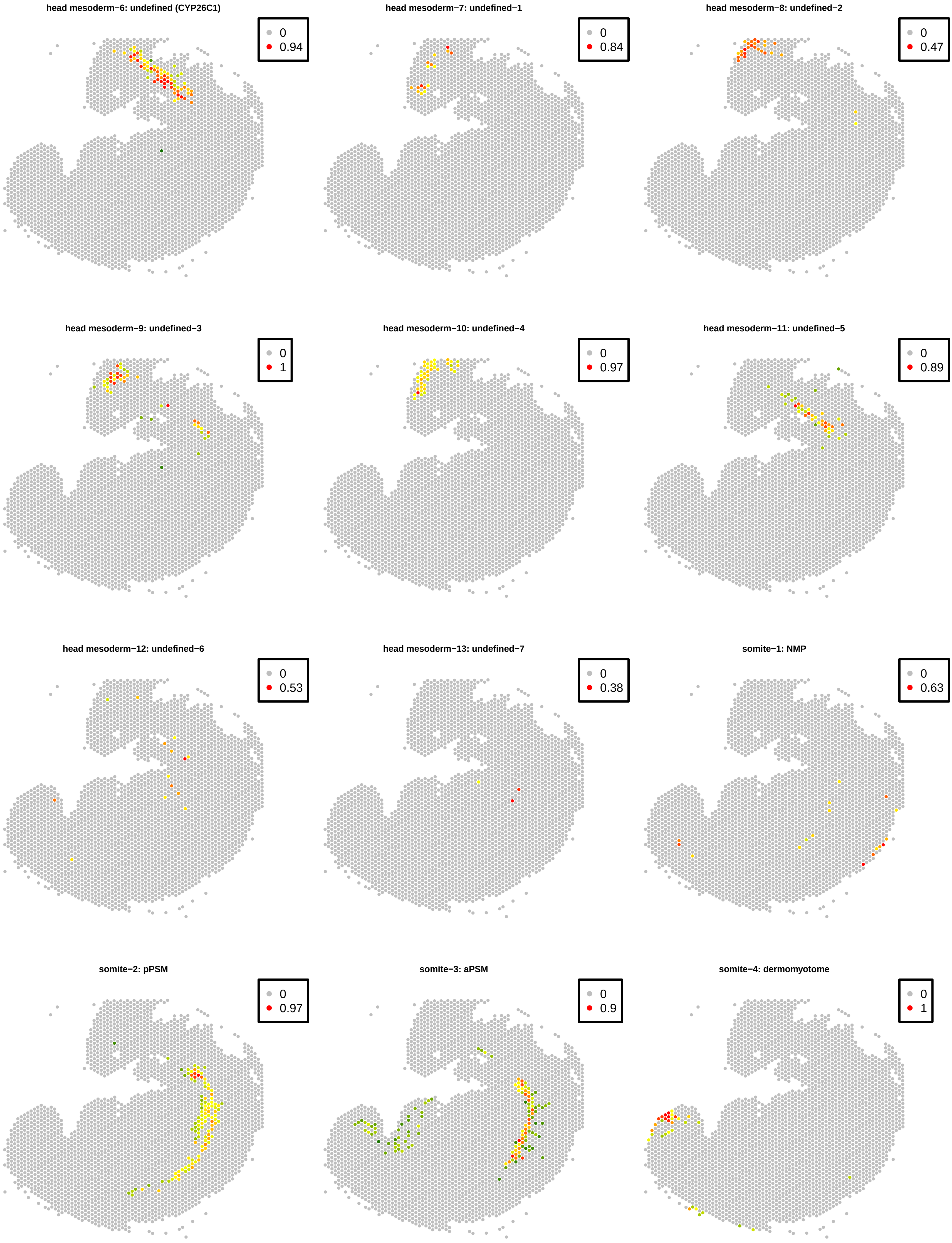
head mesoderm−6: undefined (CYP26C1)
0
0.94
head mesoderm−7: undefined−1
0
0.84
head mesoderm−8: undefined−2
0
0.47
head mesoderm−9: undefined−3
0
1
head mesoderm−10: undefined−4
0
0.97
head mesoderm−11: undefined−5
0
0.89
head mesoderm−12: undefined−6
0
0.53
head mesoderm−13: undefined−7
0
0.38
somite−1: NMP
0
0.63
somite−2: pPSM
0
0.97
somite−3: aPSM
0
0.9
somite−4: dermomyotome
0
1

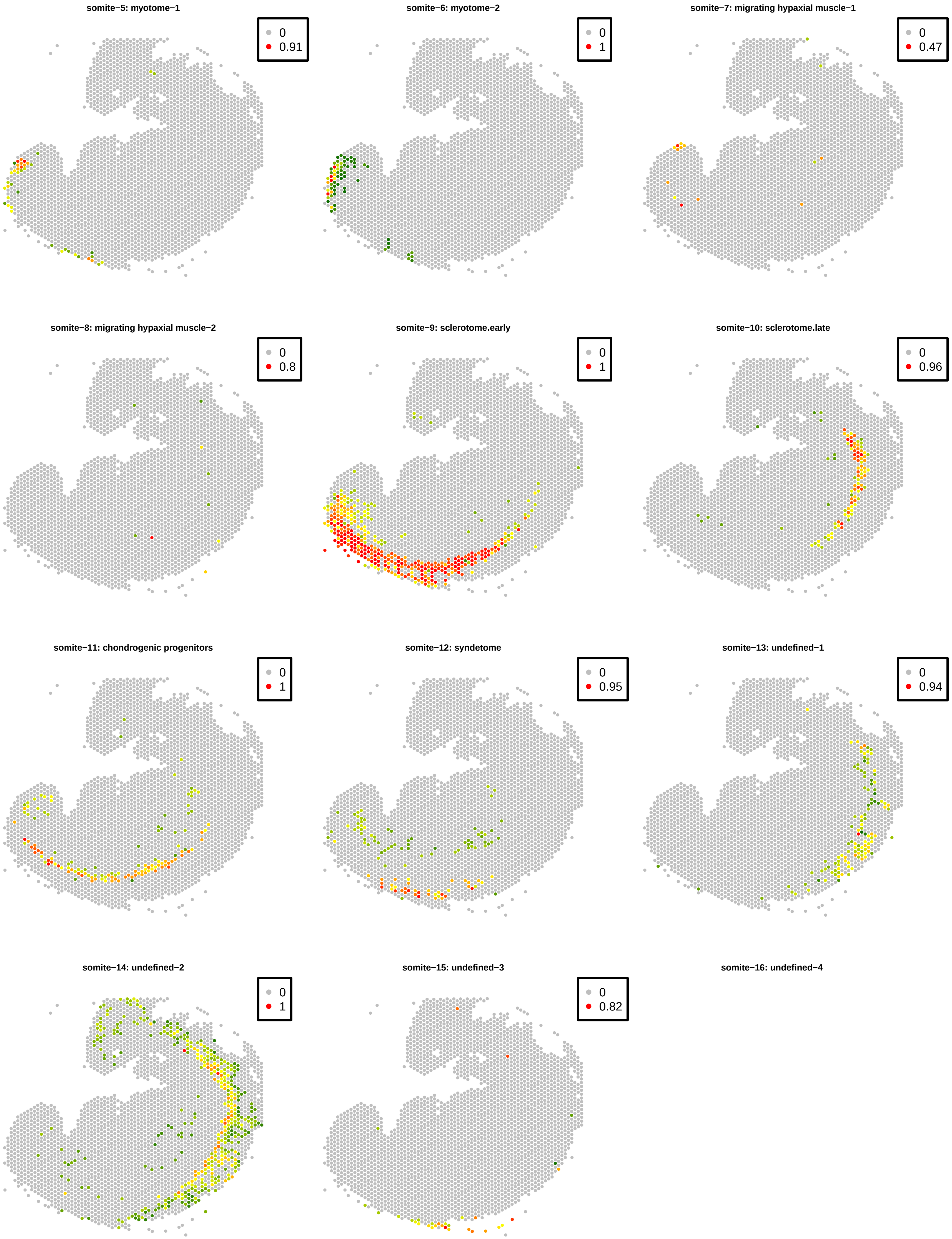

somite−5: myotome−1
0
0.91
somite−6: myotome−2
0
1
somite−7: migrating hypaxial muscle−1
0
0.47
somite−8: migrating hypaxial muscle−2
0
0.8
somite−9: sclerotome.early
0
1
somite−10: sclerotome.late
0
0.96
somite−11: chondrogenic progenitors
0
1
somite−12: syndetome
0
0.95
somite−13: undefined−1
0
0.94
somite−14: undefined−2
0
1
somite−15: undefined−3
0
0.82
somite−16: undefined−4

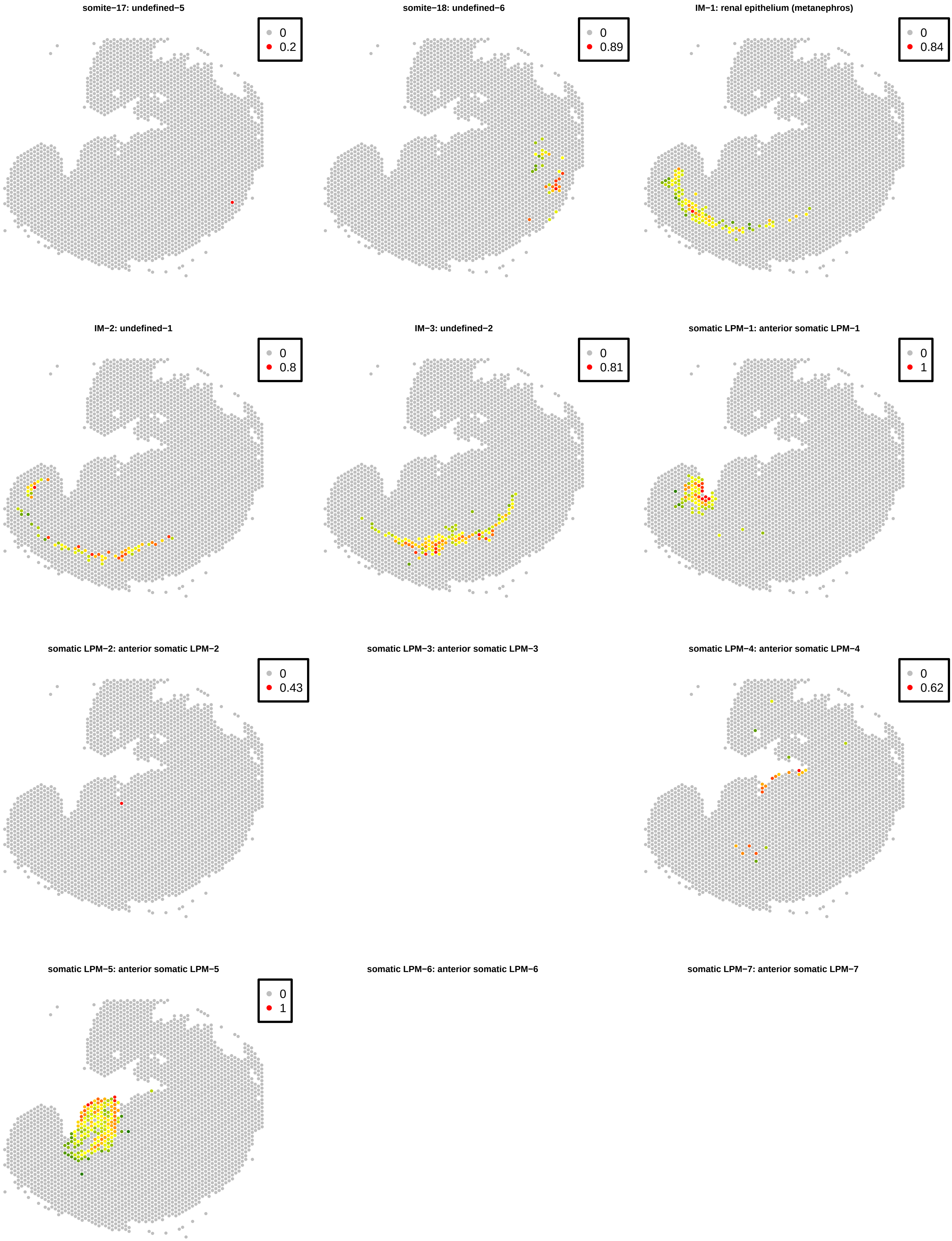
somite−17: undefined−5
0
0.2
somite−18: undefined−6
0
0.89
IM−1: renal epithelium (metanephros)
0
0.84
IM−2: undefined−1
0
0.8
IM−3: undefined−2
0
0.81
somatic LPM−1: anterior somatic LPM−1
0
1
somatic LPM−2: anterior somatic LPM−2
0
0.43
somatic LPM−3: anterior somatic LPM−3
somatic LPM−4: anterior somatic LPM−4
0
0.62
somatic LPM−5: anterior somatic LPM−5
0
1
somatic LPM−6: anterior somatic LPM−6
somatic LPM−7: anterior somatic LPM−7

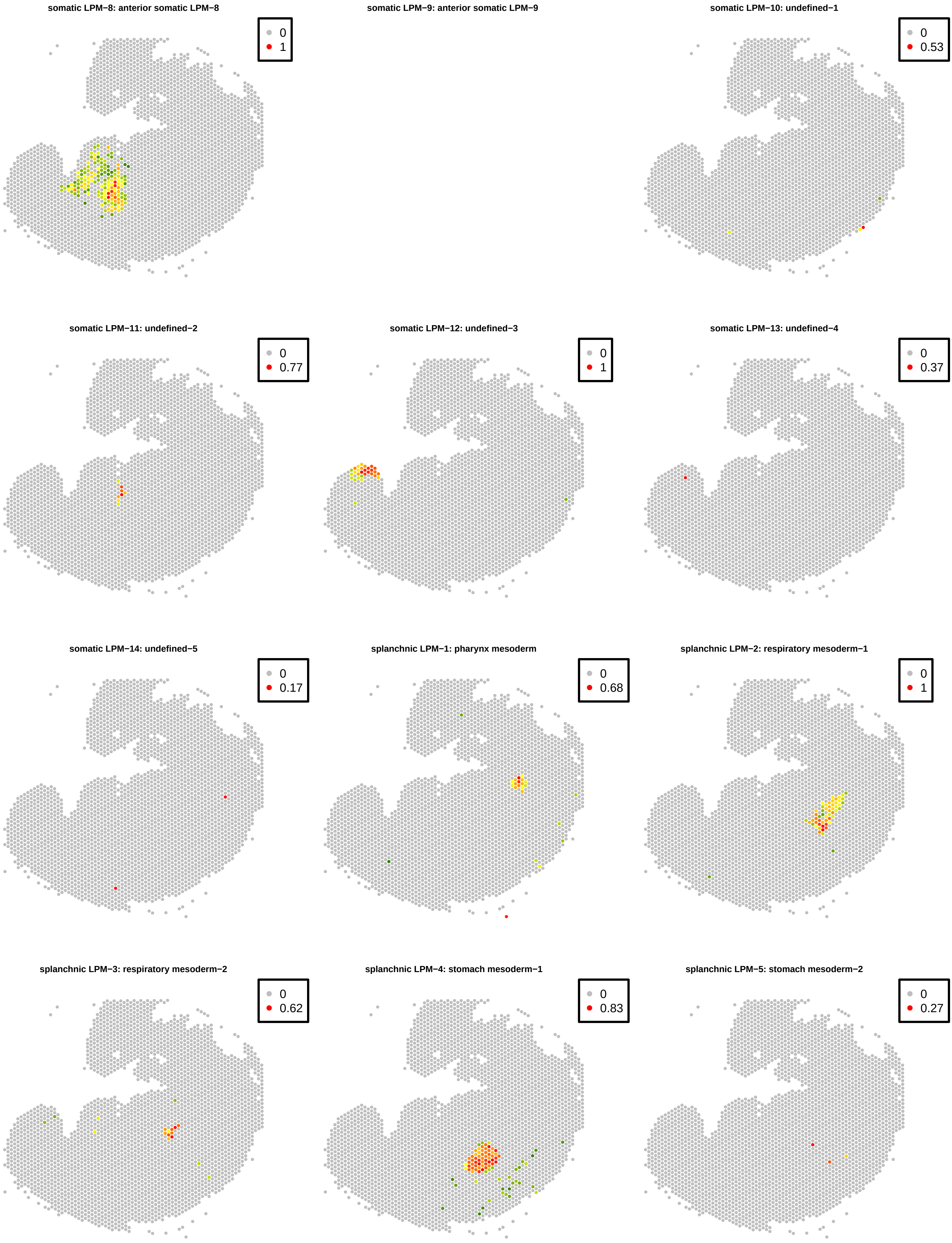
somatic LPM−8: anterior somatic LPM−8
0
1
somatic LPM−9: anterior somatic LPM−9
somatic LPM−10: undefined−1
0
0.53
somatic LPM−11: undefined−2
0
0.77
somatic LPM−12: undefined−3
0
1
somatic LPM−13: undefined−4
0
0.37
somatic LPM−14: undefined−5
0
0.17
splanchnic LPM−1: pharynx mesoderm
0
0.68
splanchnic LPM−2: respiratory mesoderm−1
0
1
splanchnic LPM−3: respiratory mesoderm−2
0
0.62
splanchnic LPM−4: stomach mesoderm−1
0
0.83
splanchnic LPM−5: stomach mesoderm−2
0
0.27

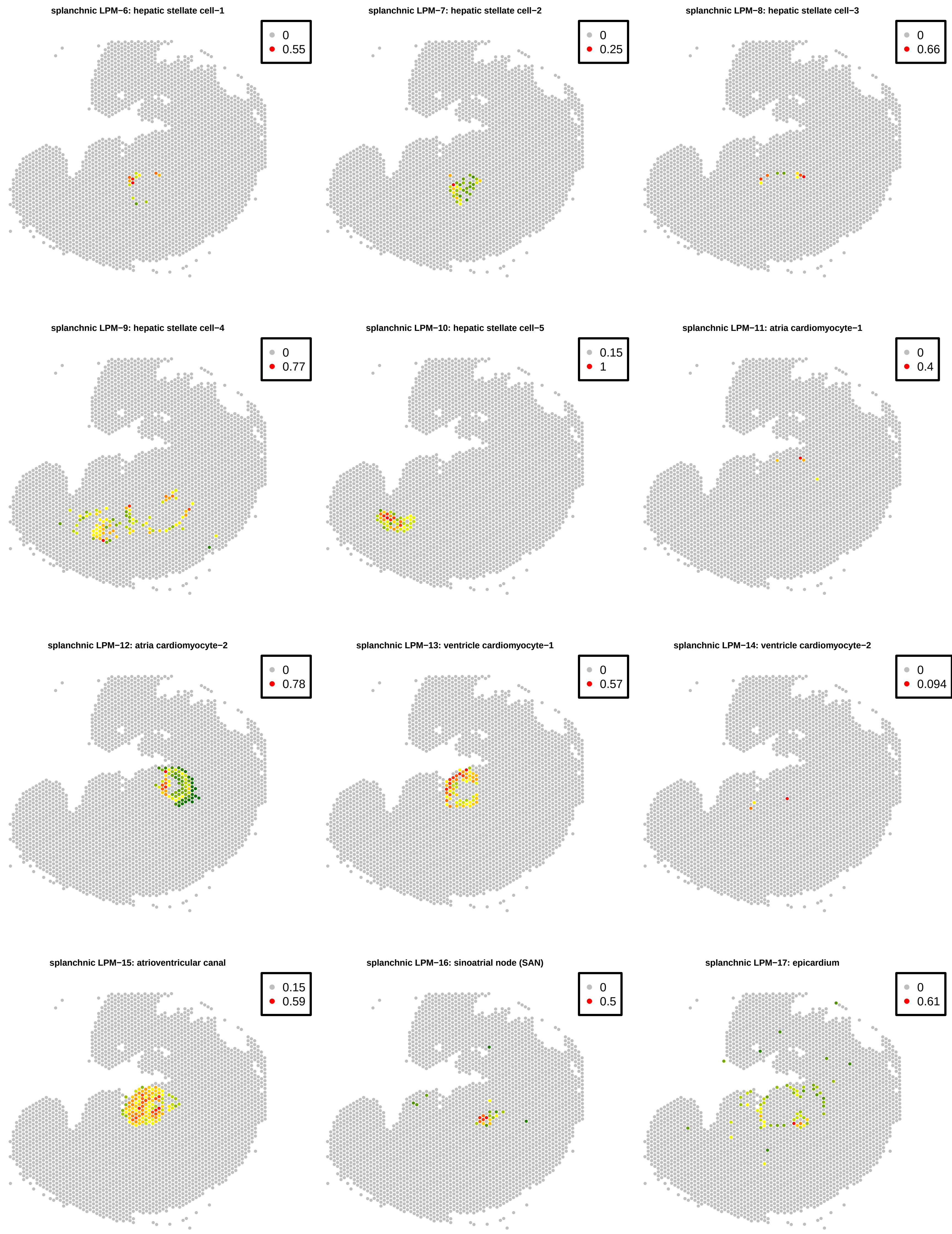

splanchnic LPM−6: hepatic stellate cell−1
0
0.55
splanchnic LPM−7: hepatic stellate cell−2
0
0.25
splanchnic LPM−8: hepatic stellate cell−3
0
0.66
splanchnic LPM−9: hepatic stellate cell−4
0
0.77
splanchnic LPM−10: hepatic stellate cell−5
0.15
1
splanchnic LPM−11: atria cardiomyocyte−1
0
0.4
splanchnic LPM−12: atria cardiomyocyte−2
0
0.78
splanchnic LPM−13: ventricle cardiomyocyte−1
0
0.57
splanchnic LPM−14: ventricle cardiomyocyte−2
0
0.094
splanchnic LPM−15: atrioventricular canal
0.15
0.59
splanchnic LPM−16: sinoatrial node (SAN)
0
0.5
splanchnic LPM−17: epicardium
0
0.61

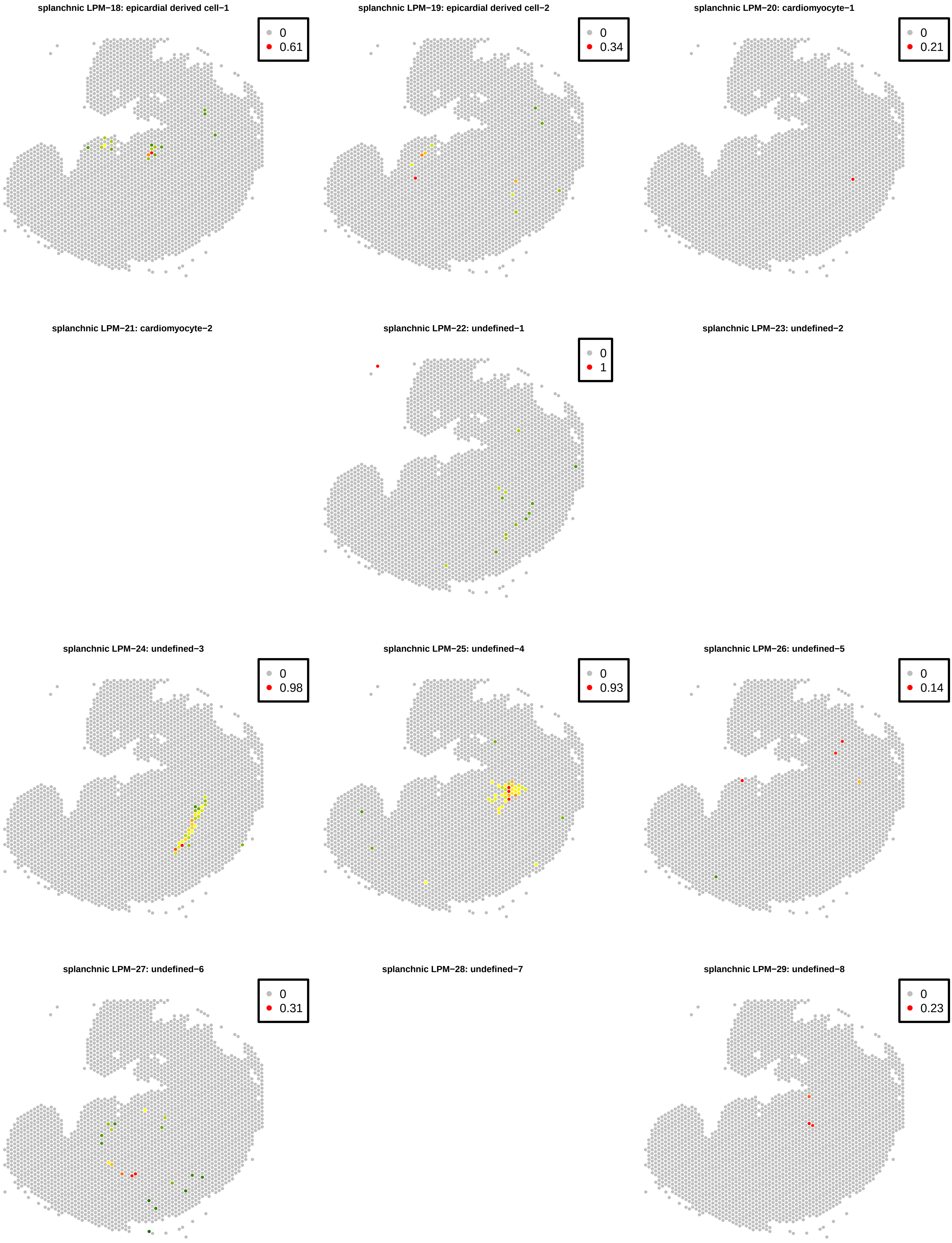
splanchnic LPM−18: epicardial derived cell−1
0
0.61
splanchnic LPM−19: epicardial derived cell−2
0
0.34
splanchnic LPM−20: cardiomyocyte−1
0
0.21
splanchnic LPM−21: cardiomyocyte−2
splanchnic LPM−22: undefined−1
0
1
splanchnic LPM−23: undefined−2
splanchnic LPM−24: undefined−3
0
0.98
splanchnic LPM−25: undefined−4
0
0.93
splanchnic LPM−26: undefined−5
0
0.14
splanchnic LPM−27: undefined−6
0
0.31
splanchnic LPM−28: undefined−7
splanchnic LPM−29: undefined−8
0
0.23

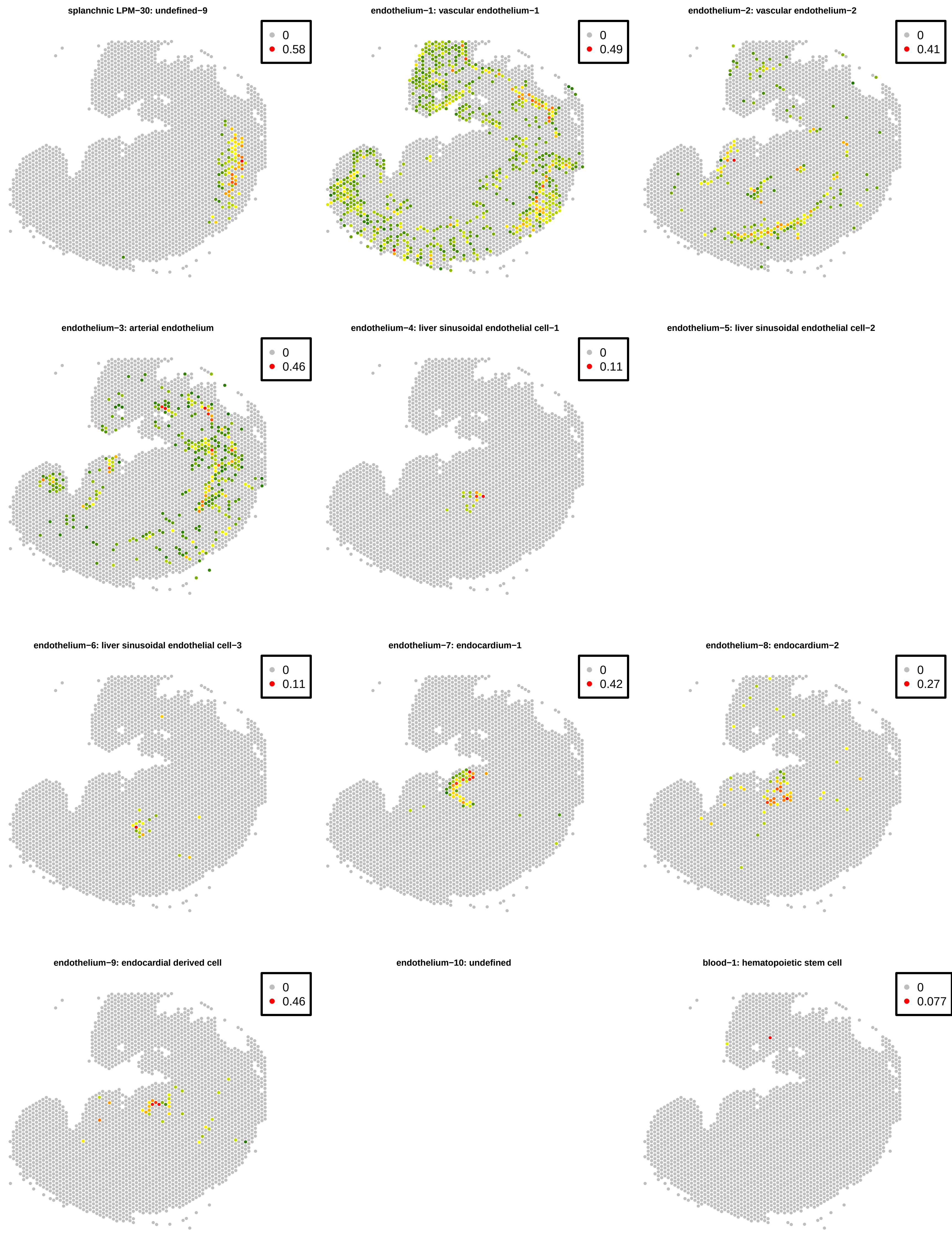
splanchnic LPM−30: undefined−9
0
0.58
endothelium−1: vascular endothelium−1
0
0.49
endothelium−2: vascular endothelium−2
0
0.41
endothelium−3: arterial endothelium
0
0.46
endothelium−4: liver sinusoidal endothelial cell−1
0
0.11
endothelium−5: liver sinusoidal endothelial cell−2
endothelium−6: liver sinusoidal endothelial cell−3
0
0.11
endothelium−7: endocardium−1
0
0.42
endothelium−8: endocardium−2
0
0.27
endothelium−9: endocardial derived cell
0
0.46
endothelium−10: undefined
blood−1: hematopoietic stem cell
0
0.077

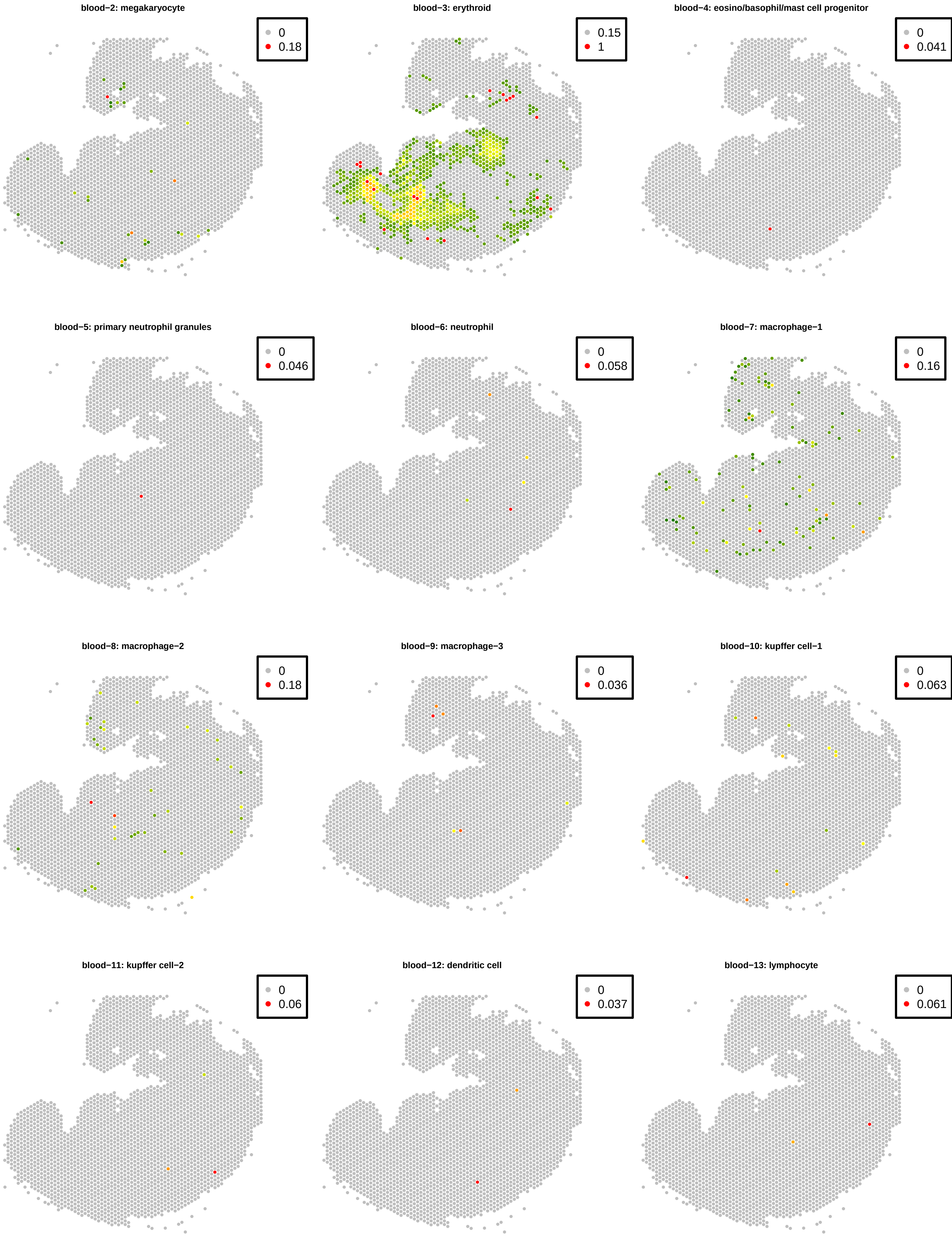
blood−2: megakaryocyte
0
0.18
blood−3: erythroid
0.15
1
blood−4: eosino/basophil/mast cell progenitor
0
0.041
blood−5: primary neutrophil granules
0
0.046
blood−6: neutrophil
0
0.058
blood−7: macrophage−1
0
0.16
blood−8: macrophage−2
0
0.18
blood−9: macrophage−3
0
0.036
blood−10: kupffer cell−1
0
0.063
blood−11: kupffer cell−2
0
0.06
blood−12: dendritic cell
0
0.037
blood−13: lymphocyte
0
0.061

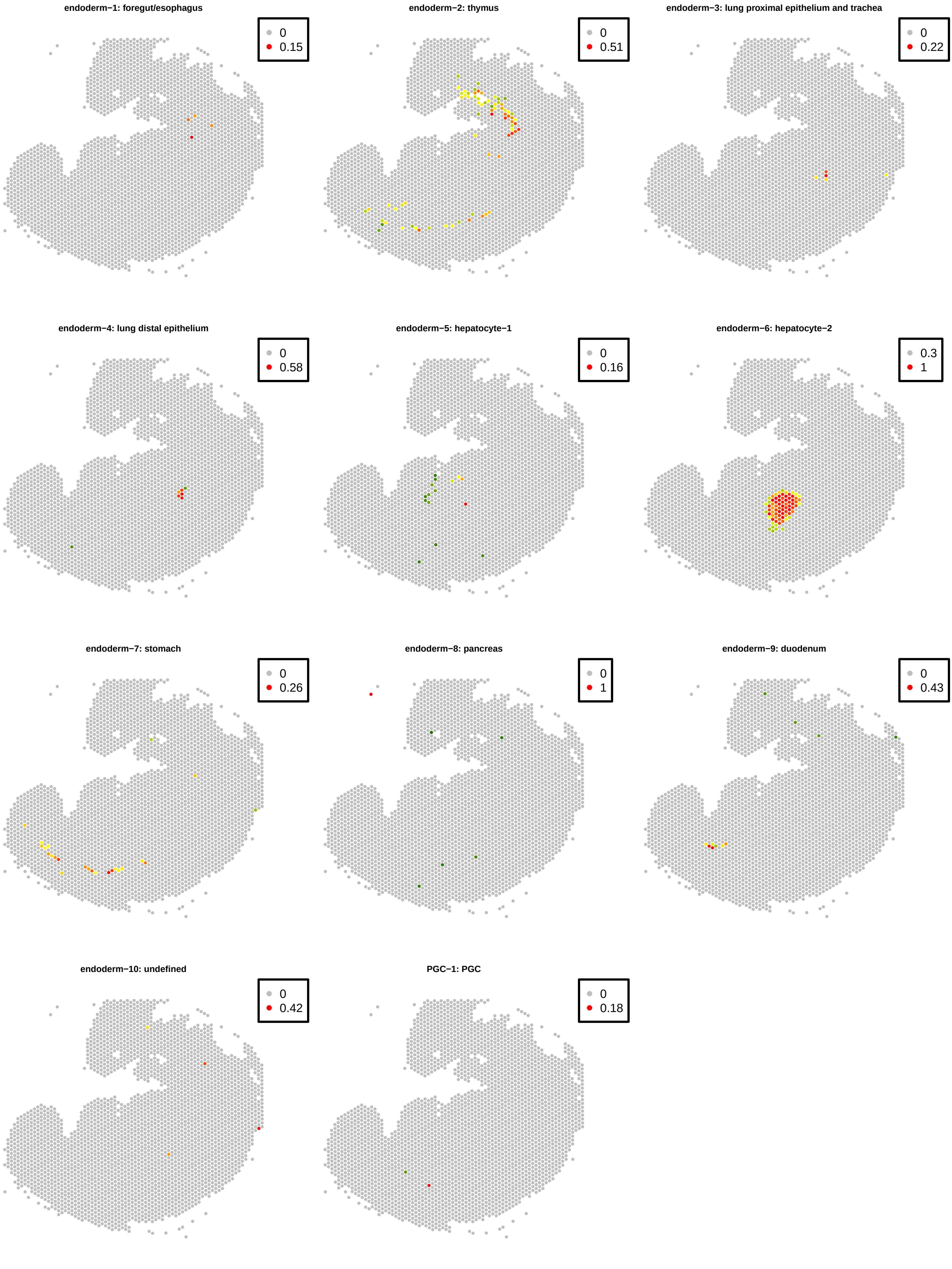
endoderm−1: foregut/esophagus
0
0.15
endoderm−2: thymus
0
0.51
endoderm−3: lung proximal epithelium and trachea
0
0.22
endoderm−4: lung distal epithelium
0
0.58
endoderm−5: hepatocyte−1
0
0.16
endoderm−6: hepatocyte−2
0.3
1
endoderm−7: stomach
0
0.26
endoderm−8: pancreas
0
1
endoderm−9: duodenum
0
0.43
endoderm−10: undefined
0
0.42
PGC−1: PGC
0
0.18
